# Interaction of amisulpride with GLUT1 at the blood-brain barrier. Relevance to Alzheimer’s disease

**DOI:** 10.1101/2023.05.15.540749

**Authors:** Sevda T. Boyanova, Ethlyn Lloyd-Morris, Christopher Corpe, K. Miraz Rahman, Doaa B. Farag, Lee K. Page, Hao Wang, Alice L. Fleckney, Ariana Gatt, Claire Troakes, Gema Vizcay-Barrena, Roland Fleck, Suzanne J. Reeves, Sarah A. Thomas

## Abstract

Blood-brain barrier (BBB) dysfunction may be involved in the increased sensitivity of Alzheimer’s disease (AD) patients to antipsychotics, including amisulpride. Studies indicate that antipsychotics interact with facilitated glucose transporters (GLUT), including GLUT1, and that GLUT1 BBB expression decreases in AD. We tested the hypotheses that amisulpride (charge: +1) interacts with GLUT1, and that BBB transport of amisulpride is compromised in AD.

GLUT1 substrates and inhibitors, and GLUT-interacting antipsychotics were identified by literature review and their physicochemical characteristics summarised. Interactions between amisulpride, and GLUT1 were studied using *in silico* approaches and the human cerebral endothelial cell line, hCMEC/D3. Brain distribution of [^3^H]amisulpride was determined using *in situ* perfusion in wild type (WT) and 5xFamilial AD (5xFAD) mice. With transmission electron microscopy (TEM) we investigated brain capillary degeneration in WT and 5xFAD mice, and human samples. Western blots determined BBB transporter expression in mouse and human.

Literature review revealed that, although D-glucose has no charge, charged molecules can interact with GLUT1. GLUT1 substrates are smaller (184.95±6.45g/mol) than inhibitors (325.50±14.40g/mol), and GLUT-interacting antipsychotics (369.38±16.04). Molecular docking showed beta-D-glucose (free energy binding: -15.39kcal/mol) and amisulpride (-29.04kcal/mol) interact with GLUT1. Amisulpride did not affect [^14^C]D-glucose accumulation in hCMEC/D3. 5xFAD mice showed increased brain [^3^H]amisulpride uptake, and no cerebrovascular space changes compared to WT. TEM revealed brain capillary degeneration in human AD. There was no significant effect of AD on mouse GLUT1 and P-gp BBB expression, and in human GLUT1 expression. In contrast, caudate P-glycoprotein expression was decreased in human AD capillaries versus controls.

This study provides new details about the BBB transport of amisulpride, evidence that amisulpride interacts with GLUT1, and that BBB transporter expression is altered in AD. This suggests that antipsychotics exacerbate the cerebral hypometabolism in AD. Further research into the mechanism of amisulpride transport by GLUT1 is important for improving antipsychotics safety.

## Introduction

Psychosis, which most commonly presents as delusions, is highly prevalent (∼50%) in people with Alzheimer’s disease (AD) (1,2), and is associated with poorer quality of life, a more rapid speed of cognitive and functional decline (2) and greater risk of institutionalisation (3). Safe and effective prescribing of antipsychotic drugs is challenging in older people, particularly those with AD, due to their heightened susceptibility to the side effects of these drugs (4), including sedation, postural hypotension, parkinsonism, and an increased risk of stroke and death (5,6). As a result, antipsychotic use is restricted to those with severe psychosis and/or aggression that have not responded to non-pharmacological approaches. In the UK, the National Institute for Health and Care Excellence guidance advocates use of ‘the lowest possible dose for the shortest possible time’ (7). There is a lack of guidance on the minimum clinically effective dose for individual drugs, or the factors that predict differing response and side effects, although a recent publication, based on clinical data on risperidone use in AD, suggests that increased dementia severity is an independent risk factor for emergent parkinsonism (7).

Amisulpride is a second-generation antipsychotic – a substituted benzamide derivative, which is a highly selective dopamine D2 receptor antagonist (8). It can be prescribed off-label in very late onset (> 60 years) schizophrenia-like psychosis (9), and in patients with AD psychosis (10–12). Therapeutic drug monitoring studies in adults predominantly aged under 65 years have shown that therapeutic striatal D2/3 receptor occupancies between 40-70% are achieved at blood drug concentrations of 100-319 ng/ml; equivalent to 400-800 mg/day (13–15). Older patients with AD psychosis (aged 69-92 years) showed a clinical response to amisulpride and parkinsonian side effects at lower doses (25-75 mg/day) with high D2/3 receptor occupancies in the caudate (41-83%) and lower blood drug concentrations (40-100 ng/ml) than expected (11,12).

These findings suggest that age and/or AD-related changes in central pharmacokinetics contribute to antipsychotic drug sensitivity and implicate the blood-brain barrier (BBB) which controls the entry of drugs to the brain through selective transport pathways. Evidence from our animal studies further supports this hypothesis (16). Since amisulpride is predominantly positively (17) charged at physiological pH, observed changes in the expression and/or function of BBB transporters for organic cations may help explain the increased amisulpride sensitivity in AD patients. In particular, the organic cation transporter 1 (OCT1; SLC22A1) and the plasma membrane monoamine transporter (PMAT; SLC29A4) (16).

Compromised function and expression of other SLC transporters at the BBB, such as the glucose transporter, GLUT1 (SLC2A1), has also been observed in AD (18–20). Consequently, AD brains suffer chronic shortages of energy-rich metabolites (21,22). Importantly, antipsychotic drugs, including risperidone and clozapine, inhibit glucose uptake by transporters, it has been suggested that this could be through direct interaction with GLUT transporters (23,24). In addition, the use of clozapine and another antipsychotic drug, olanzapine, has been associated with the development of type 2 diabetes. One of the proposed mechanisms is through the inhibition of glucose transporters inducing hyperglycaemia (25–27). However, detailed studies of the drugs-transporter interaction, including *in silico* molecular docking studies on the specific molecular interactions of antipsychotics with GLUT1 are rare.

Thus, considering the changes in GLUT1 expression at the BBB and resulting impact on brain metabolism in AD, and the increased sensitivity of AD patients to antipsychotic drugs including risperidone and amisulpride, we wanted to explore the interaction between GLUT1 and amisulpride. To do this we tested the hypotheses that amisulpride interacts with GLUT1 at the BBB, and that expression of BBB transporters and the transport of amisulpride and glucose into the brain will be affected by AD. We utilised a combination of literature review, physicochemical properties analysis, molecular docking approaches, cell culture BBB studies, studies in wild type mice and in an animal model of AD and assessment of tissue from human cases with and without AD. An overview of the methods deployed are presented in Supplementary Fig 1, S1 File. Abstracts of this work have been presented (28).

## Materials and Methods

### Radiolabelled and non-labelled chemicals

Radiolabelled [^14^C]D-glucose (#NEC042X050UC, Lot: 2389266, specific activity 275 mCi/ mmol) and [^3^H]mannitol (#NET101001MC, Lot: 3632303, specific activity 12.3 Ci/ mmol) were purchased from Perkin Elmer. [O-methyl-^3^H]amisulpride (MW 374.8 g/mol; specific activity 77 Ci/mmol; 97% radiochemical purity) was custom tritiated (#TRQ41291 Quotient, UK). [^14^C(U)]Sucrose (MW 359.48 g/mol; specific activity 536 mCi/mmol; 99% radiochemical purity: # MC266) was purchased from Moravek Biochemicals, USA. Non-labelled amisulpride (MW 369.5 g/mol, >LJ98% purity) was purchased from Cayman Chemicals, UK (#71675-85-9), and non-labelled D-glucose (#10117) was purchased from BDH.

### Literature review

Three different groups of molecules, which interacted with GLUT, were identified by literature review. These groups were GLUT1 substrates, GLUT1 inhibitors and GLUT-interacting antipsychotics. The inclusion criteria are explained below.

### Identification of GLUT1 substrates and inhibitors

To identify established substrates and inhibitors of GLUT1 we performed a Pubmed literature search using the parameters ((GLUT1 substrate) OR (GLUT1 inhibitor)) AND (review [Publication Type])) AND ((”1985” [Date - Publication]:”2021” [Date - Publication])) (29) and tabulated the results. In the substrates group we included molecules for which there is *in vitro* evidence that they are transported by GLUT1 and that their uptake into cells is inhibited by established GLUT1 inhibitors, such as cytochalasin B. In the inhibitors group we included molecules that were shown to decrease uptake of GLUT1 substrates *in vitro*, and for which there is a proposed inhibitory mechanism of interaction with GLUT1.

### Identification of antipsychotics that interact with GLUT

We also performed a Pubmed search to identify antipsychotics which interacted with GLUT transporters, using the parameters (((GLUT) AND (antipsychotic)) AND (("1985"[Date - Publication]: "2021"[Date - Publication])) (accessed 04/03/2022). In the antipsychotics group we included both typical and atypical antipsychotics that decreased the uptake of GLUT1 substrates *in vitro*.

### Physicochemical characterisation of GLUT1 substrates and inhibitors, and GLUT-interacting antipsychotics

The physicochemical characteristics of each member of the three groups (i.e. GLUT1 substrates, GLUT1 inhibitors and GLUT-interacting antipsychotics) was obtained from the chemical property databases, DrugBank (30) or MarvinSketch (version 22.9.0, 2022, ChemAxon) (31) and tabulated.

Specifically, information about the structure and molecular weight (MW) was obtained from the DrugBank (30). The mean MW ± SEM, and the median MW of the three groups was then calculated and the results compared.

The gross charge distribution at pH 7.4 of each molecule was then obtained from MarvinSketch. The mean gross charge distribution at physiological pH (± SEM) of the three groups was then calculated and the results compared. The number of microspecies of each molecule at pH7.4 was examined and the percentage prevalence and charge of the top two microspecies was then reported (MarvinSketch). The physicochemical characteristics of the three groups were compared to those obtained for amisulpride.

### *In silico* molecular docking

*In silico* molecular docking was used to examine the molecular level interactions of amisulpride, alpha-D-glucose, beta-D-glucose, and sucrose with GLUT1 using the molecular docking tool GOLD. The main endogenous substrate of GLUT1 is D-glucose, which is a monosaccharide present in the body as anomers: 36% alpha-D-glucose and 64% beta-D-glucose (32). In this molecular docking study alpha-D-glucose and beta-D-glucose were used as positive controls. Sucrose is a plant disaccharide which is thought not to interact with GLUT1 and was used as the negative control.

The GLUT1 Protein Database (PDB) code used was 5EQI – the crystal structure of human GLUT1. The 5EQI crystal structure is for the inward open conformation of the transporter which has been reported to be the most favourable for ligand binding (33).

The results from the molecular docking simulations were expressed as a free energy binding and chem score. A molecule is considered a substrate for a given transporter if their interaction has a free energy binding of ≤5kcal/mol and has a high chem score. The chem score is one of the scoring functions of the molecular docking program. It provides information on the strength of the interaction between ligand and binding sites.

### *In vitro* studies in model of the human BBB (hCMEC/D3 cells)

The human cerebral microvessel endothelial cells/D3 (hCMEC/D3) are an immortalised human adult brain endothelial cell line. This is a well-established and characterised model, regularly used to study the human BBB (16,34). The hCMEC/D3 cell line originated from human brain tissue obtained following surgical excision of an area from the temporal lobe of an adult female with epilepsy (34).

hCMEC/D3 cells were used to study the interaction between GLUT1 and amisulpride. To do this, the expression of a functional GLUT1 was first confirmed using Western blot (WB) and accumulation assays with [^14^C]D-glucose. This was followed by further accumulation assays with [^14^C]D-glucose and non-labelled amisulpride. [^3^H]mannitol was used as a marker of cellular integrity and non-specific binding to the membranes and plastic wear. The hCMEC/D3 cells (passages 30-35) were grown as described previously by Sekhar et al., 2019 (16). All experiments were carried out at King’s College London, in accordance with the guidelines of the Local Ethics Committee and research governance guidelines.

### Western blot studies – expression of GLUT1 in hCMEC/D3 cells

Cells were grown for four to five days in T-75 flasks until they formed a fully confluent monolayer, and then for another two to three days before harvesting and preparing lysates for WB. Lysates were prepared by placing the flask on ice, aspirating the medium and washing the cells in ice-cold phosphate buffered saline (#70011-36 Gibco, Fischer Scientific Ltd, UK). The cells were then incubated on a shaker for 10 minutes in radio-immunoprecipitation assay (RIPA) buffer (#R0278, Sigma, UK) and 1% (v/v) protease inhibitors (#78441, Thermo Fisher Scientific, UK) for cell lysis and protein solubilisation. The cells were then scraped off from the bottom of the flask and transferred to an Eppendorf tube. The tube was then agitated for 20 to 30 minutes at 4°C. Next, the cell suspension was centrifuged (Biofuge Fresco, Heraeus Instruments, UK) for 20 minutes at 11,753 x G at 4°C. The resulting supernatant (protein lysate) was snap frozen in liquid nitrogen. Before the start of the WB procedure, the protein lysates were thawed and the total protein concentration in each lysate was estimated by bicinchoninic acid (BCA) assay using bovine serum albumin standards (Thermo Scientific, UK) as described by Sekhar et al., 2019 (16).

The protein lysates for WB were mixed with sample loading buffer (1:4) (#NP0007 NuPAGE LDS Sample Buffer (4x), Invitrogen, Carlsbad, USA), reducing agent (1:10) (#B0009, Novex by Life Technologies, USA), and RIPA buffer. Once prepared, they were heated at 95°C for 5 minutes. The rest of the WB procedure was performed as described in (16) except, 30 µg of protein was loaded in each well, antibody solutions were made in 5% milk in TBS-T, and washes were performed in TBS-T (for antibodies dilutions see Supplementary Table 1, S1 File). Quantification of protein expression was determined by calculating the intensity ratio of the band of interest and the band of the loading control (GAPDH). Band intensity ratio analysis was conducted using ImageJ software (35).

### Accumulation assays – function of GLUT1 in hCMEC/D3 cells

The cell culture method and experimental design of the accumulation studies are described in the following sections. The cells were grown as already described in the “*In vitro* studies” section. They were split at 80-90% confluence and were seeded at 25,000 cells/cm^2^ in the central 60 wells of 96 well plates for accumulation assay experiments (#10212811, Fisher Scientific, UK). They were grown for four to five days until they formed a fully confluent monolayer, and then for another two to three days before the start of the accumulation assay.

For the control accumulation assay experiments, the hCMEC/D3 cells were incubated in an accumulation buffer (135 mM NaCl, 10 mM HEPES, 5.4 mM KCl, 1.5 mM CaCl_2_H_2_O, 1.2 mM MgCl_2_.H_2_O, 1.1 mM D-glucose - the standard non-labelled D-glucose concentration for the accumulation buffer, and water, pH = 7.4) for 1 h. The buffer also contained [^14^C]D-glucose (3 µM) and [^3^H]mannitol (0.05 µM). [^3^H]Mannitol was used as a marker for non-specific binding, and barrier integrity (36,37). All treatment conditions contained 0.05% DMSO. Incubation was performed in a shaker at 37°C and 120 rotations per minute. Next, the cells were washed with ice-cold phosphate buffered saline (#BR0014G, Oxoid Limited, England) to stop the accumulation process and to remove any radiolabelled molecules and buffer that had not entered the cells. Then the cells were incubated in 1% Triton X (200 µl per well) for 1 h at 37°C, to solubilise the cell membranes, and to free any accumulated radiolabelled compounds via lysis. Half of the cell Triton X lysate from each well was pipetted into a scintillation vial and 4 ml of scintillation fluid (#6013329, Ultima GoldTM, Perkin Elmer, USA) was added. The amount of ^14^C and ^3^H radioactivity in the sample was measured in disintegrations per minute (dpm) on a Tricarb 2900TR liquid scintillation counter. The dpm of each sample were corrected for background dpm. The background dpm was determined from a vial containing 100 µl 1% Triton X and 4 ml liquid scintillation fluid only. The remaining cell lysate in the plate was used for total protein concentration control in each well, determined by BCA analysis (17).

In our self-inhibition studies, investigating the presence of a functional GLUT1 transporter in the hCMEC/D3 cells, the cells were incubated with accumulation buffer, containing [^14^C]D-glucose (3 µM), [^3^H]mannitol (0.05 µM) and 4 mM non-labelled D-glucose. The 4 mM total concentration of non-labelled D-glucose included the standard 1.1mM non-labelled D-glucose normally present in accumulation buffer. Radioactivity uptake was compared to the control condition in which the cells were treated with accumulation buffer of the same composition apart from the non-labelled D-glucose content which was 1.1 mM (the standard non-labelled D-glucose concentration in the accumulation buffer). All treatments contained 0.05% DMSO. After 1 h incubation, the cells were lysed and used for liquid scintillation counting and BCA assay as already described. The non-labelled D-glucose concentration for the treatment condition (4 mM) was selected based on previous reports that normal plasma glucose concentrations are kept in a narrow range between 4 and 8 mM in healthy subjects (38) and on preliminary experiments studying the effect of 4, 6, and 9 mM non-labelled D-glucose on the membrane integrity of hCMEC/D3 cells (see Supplementary Fig 2, S1 File).

In order to assess the effect of amisulpride on the transport of D-glucose through the hCMEC/D3 membrane, non-labelled amisulpride (20, 50, or 100 µM) was added to the accumulation buffer, containing [^14^C]D-glucose (3 µM), [^3^H]mannitol (0.05 µM), and non-labelled D-glucose (1.1 mM - total standard concentration of non-labelled D-glucose in the accumulation buffer). All treatment conditions contained 0.05% of DMSO. The cells were incubated with this mixture for 1 h, the BCA assay and liquid scintillation counting was performed as described previously.

The total cellular accumulation of [^14^C]D-glucose was expressed as a volume of distribution (V_d_; µl/mg). V_d_ was calculated as the ratio of disintegrations per minute (dpm)/mg protein in the Triton X lysate to dpm/ µl of the accumulation buffer. The V_d_ values for [^3^H]mannitol were subtracted from the V_d_ values for [^14^C]D-glucose to correct for cell membrane integrity. This value was then divided by the protein content in each well to correct for protein content. Values were expressed as a mean ± SEM.

Statistical analysis of the results from the self-inhibition studies was performed using unpaired one-tailed Student’s t-test. The results from the studies of the interaction between amisulpride and GLUT1 were analysed using One-way ANOVA. All statistical analysis was performed with GraphPad Prism 9.0.

### Animal model studies in WT and 5xFamilial AD (FAD) mice

The 5xFAD mice are a model of AD which carry human amyloid-β precursor protein (APP) with the Swedish (K670N, M671L), Florida (I716V), and London (V717I) FAD mutations along with human PSEN-1 with two FAD mutations: M146L and L286V under the murine Thy-1 promotor (39). This model has been described as one of the few models that shows several AD hallmarks, including neural loss, neurodegeneration, gliosis, and spatial memory deficits (40).

We confirmed the model phenotype in the following ways: comparing the weight of females and males from the two genotypes; using transmission electron microscopy (TEM) to confirm the presence of amyloid plaques in the brain of 5xFAD mice; comparing the expression of APP in brain capillary enriched pellets isolated from WT and 5xFAD mice. Enrichment of the pellets with brain endothelial cells was confirmed using transferrin receptor 1 (TfR1) as an endothelial cell marker.

### Animal husbandry

All *in vivo* experiments were performed in accordance with the Animal Scientific Procedures Act (1986) and Amendment Regulations 2012 and with consideration to the Animal Research: Reporting of *In Vivo* Experiments (ARRIVE) guidelines. The study was approved by the King’s College London Animal Welfare and Ethical Review Body. The UK government home office project license number was 70/7755.

The mice were housed at King’s College London in groups of three or four under standard conditions (20-22°C, 12 h light/dark cycle) with food and water *ad libitum*. They were housed in guideline compliant cages. Animal welfare was assessed daily by animal care technicians. Animals were identified by earmarks. The experimenter was not blinded to the mouse genotype.

The WT (C57/BL6) mice and the 5xFAD mice (on C57/BL6 background) were between 4.5 and 15 months old. The average lifespan of the C57/BL6 mice has been reported to be 30 months (41). Whereas the 5xFAD mice have been reported to have lifespan of approximately 15 months (42), with some authors reporting a median lifespan of up to 24.6 months (41).

### TEM ultrastructural study

To assess Aβ plaque presence in the brain in WT and 5xFAD mice we used TEM. All studied animals were female. The animals of each genotype were grouped into two age ranges. The groups were WT 4.5-6 months, 5xFAD 6 months, WT 12 months, 5xFAD 12 months with n=2 mice per group. The weight of the animals was between 21.9 g and 26.7 g. For brain dissection, all animals were terminally anaesthetised with intraperitoneal injection of pentobarbital (100 µl/animal) (Fort Dodge, Southampton, UK).

Once each mouse was anaesthetised, it was perfuse-fixed. The left ventricle of the heart was infused with ice-cold phosphate buffered saline and the right atrium sectioned to provide an open circuit. Once the vasculature was free of blood, 4% paraformaldehyde (PFA, #F017/2, TAAB, UK) was infused via the heart. Fixation with 4% PFA was performed for 10 minutes. The brain was then removed, and the frontal cortex was dissected, and cut into 1 mm^3^ samples.

The brain samples were processed, and sectioned for TEM, and then imaged. The samples were incubated overnight at 4°C in TEM fixative, containing 2.5% (v/v) glutaraldehyde in 0.1 M cacodylate buffer (pH 7.3). The tissues were post fixed in 1% (w/v) osmium tetroxide (#O021, TAAB, UK) for 1.5 h at 4°C. Then they were washed and dehydrated through serial graded incubations in ethanol: 10%, followed by 70%, and 100%. The tissue was infiltrated in embedding resin medium (#T028, TAAB, UK) for 4 h at room temperature. The samples were embedded on flat moulds and polymerised at 70°C for 24 h. Ultrathin sections (100-120 nm) were cut on a Leica ultramicrotome (Leica microsystems, Germany) and mounted on mesh copper grids. They were contrasted with uranyl acetate for 2 minutes and with lead citrate for 1 minute. The sections were imaged on an EM-1400 (Plus) transmission microscope operated at 120 kV (JEOL USA, Inc.).

For each frontal cortex sample, a minimum of three images were collected, and amyloid plaques were identified and labelled with reference to electron microscopic atlas of cells, tissues, and organs (43). The dimensions of each image were 3296x2472 pixels.

### Western blot studies

We isolated the brain capillaries of each mouse used in the *in situ* brain perfusion experiments. The mouse brain was *in situ* perfused via the heart with artificial plasma, infused with rabiolabelled amisulpride and radiolabelled sucrose for 10 minutes. Then it was homogenised in capillary depletion buffer (10.9 mM HEPES, 141 mM NaCl, 4 mM KCl, 2.8 mM CaCl_2_ (aqueous solution - 1M), 1 mM MgSO_4_.7H_2_O, 1 mM NaH_2_PO_4_, 10 mM glucose) (brain weight x 3) and 26% dextran (MW 504.4 g/mol) (#J14495.A1, VWR, UK) (brain weight x 4). The homogenate was centrifuged at 5,400 G for 15 min at 4°C. This resulted in separation of an endothelial cell-enriched pellet and a supernatant containing the brain parenchyma and interstitial fluid. Half of the capillary pellets were snap frozen in liquid nitrogen and used for WB analysis to test for TfR1, and BBB transporter expression, including GLUT1, and P-glycoprotein (P-gp).

For WB, the mouse endothelial cell enriched capillary pellets were thawed, homogenised in 250 µl RIPA buffer with added protease inhibitor (1% v/v). The tissue was incubated in the buffer at 4°C for 30 minutes and then centrifuged (Biofuge Fresco, Heraeus Instruments, UK) at 7,999 G for 15 minutes at 4°C. The resulting supernatant was used for WB analysis. The rest of the WB procedure was performed as previously described in this paper. For antibodies used, see Supplementary Table 1, S1 File.

Quantification of protein expression was determined by calculating the intensity ratio of the band of interest and the band of the loading control (GAPDH or tubulin). Band intensity ratio analysis was conducted using ImageJ software (35).

Unpaired two-tailed Student’s t-test was used for statistical analysis of the difference in expression of each transporter studied between the WT and the 5xFAD mice.

### *In situ* brain perfusions

The *in situ* brain perfusion technique allows examination of the movement of slowly moving molecule across the BBB in the absence of systemic metabolism. This method was used to compare [^3^H]amisulpride and [^14^C]sucrose uptake into the brain in WT and in 5xFAD mice.

The mice were terminally anaesthetised with medetomidine hydrochloride (2 mg/kg, Vetoquinol UK Limited) and ketamine (150 mg/kg, Pfizer, UK, and Chanelle, UK), injected intraperitoneally. Heparin was injected intraperitoneally before the perfusion (100 units heparin dissolved in 0.9 % NaCl (aqueous solution) (heparin – batch number: PS40057; NaCl - Sigma Aldrich, Denmark). Two experimental groups were perfused – WT (12-15 months old, n=7, n=4 females, n=3 males) and 5xFAD (12-15 months old, n=7, n=4 males, n=3 females). Weights of the perfused mice were between 16.8 g and 41.5 g. Animals with a weight lower than 25 g were excluded from the perfusion analysis as perfusion at a flow rate of 5.5 ml/min could cause loss of BBB integrity (44) but their capillary lysates were used in WB experiments.

Artificial plasma was used in the perfusion experiments. It contained 117 mM NaCl, 4.7 mM KCl, 2.46 mM MgSO_4_.7H_2_O, 24.8 mM NaHCO_3_, 1.2 mM KH_2_PO_4_, 2.5 mM CaCl_2_ (aqueous solution - 1M), 39 g/L Dextran, 10 mM glucose, 1g/L bovine serum albumin, and it was mixed with Evan’s blue dye (0.0551 g per 1 L of artificial plasma, #E2129-10G, Sigma Life Science, India). It was warmed to 37°C and oxygenated by 95% O_2_/5%CO_2_ gas bubbled through the solution. The artificial plasma also contained [^3^H]amisulpride (6.5 nM) and [^14^C]sucrose (9.4 µM). Perfusion time was 10 minutes.

After the perfusion, the brain was dissected out and weighed. The frontal cortex, striatum, thalamus, and hypothalamus were dissected under a microscope (Leica, Wetzlar, Germany), weighed, and solubilised in Solvable (#6NE9100, PerkinElmer, Inc., USA) for 2-3 days, then they were taken for liquid scintillation counting. These regions were selected to compare with data from other *in situ* brain perfusion experiments and observations from human brain data sets (16) focused on similar areas, including the caudate nucleus and the putamen which are part of the striatum (45).

The rest of the brain was used for capillary depletion as already described. The whole brain homogenate, supernatant (containing brain parenchyma and interstitial fluid), and half of the capillary pellets were also solubilised and used for liquid scintillation counting.

The concentration of [^3^H] or [^14^C] radioactivity present in the brain (disintegrations per minute per gram of tissue – dpm/g) was expressed as a percentage of the concentration of radioactivity detected in the artificial plasma (disintegration per minute per millilitre). The value obtained was named %Uptake showing the radioactivity in ml/g of tissue x 100. The %Uptake values for [^3^H]amisulpride were corrected for vascular space by subtracting the corresponding [^14^C]sucrose Uptake values from the [^3^H]amisulpride Uptake values.

Statistical analysis of the difference between the uptake of [^14^C]sucrose corrected [^3^H]amisulpride into the brain of WT and 5xFAD was performed using unpaired two- tailed Student’s t-test for the capillary pellet, supernatant, and homogenate . The difference between WT and 5xFAD mice in the [^3^H]amisulpride uptake in the brain areas studied was analysed with Mixed-effects analysis with Holm-Sidak post hoc test. The same tests were used to analyse the difference in the [^14^C]sucrose brain uptake between WT and 5xFAD mice.

### Human control and AD tissue studies

We used TEM to examine the brain and brain capillary ultrastructure of an AD case. Brain capillary depletion samples from age-matched human control and AD cases were used to evaluate and compare the total protein levels and the expression levels of TfR1, GLUT1 and P-gp in the two groups.

### Ethics statement

Human tissue brain samples were provided via Brains for Dementia Research (BDR) and were anonymised. Written consent was provided by BDR and the specific BDR reference numbers were: TRID_170, TRID_170 amendment, TRID_265 and TRID_287.

BDR has ethical approval granted by the National Health Service (NHS) health research authority (NRES Committee London-City & East, UK: REC reference:08/H0704/128+5. IRAS project ID:120436). Tissue samples were supplied by The Manchester Brain Bank and the London Neurodegenerative Diseases Brain Bank, both of which are part of the BDR programme, jointly funded by Alzheimer’s Research UK and Alzheimer’s Society. Tissue was received on the basis that it will be handled, stored, used, and disposed of within the terms of the Human Tissue Act 2004. The human samples were collected during February 2012 to February 2019. The human tissue studies were conducted during 1^st^ October 2018 to 29^th^ January 2021. The authors conducting the human tissue analysis studies did not have access to information that could reveal the identity of the human tissue donors.

### TEM ultrastructural study

The tissue used was from one case which was received from the London Neurodegenerative Diseases Brain Bank, Denmark Hill, King’s College London. Case details: BBN002.32856; sex: female; age: 74; post-mortem delay: 19 h; pathological diagnosis: Alzheimer’s disease, modified Braak staging (BrainNet Europe – BNE staging) stage VI.

Samples with size of up to 1 x 5 x 1 mm were dissected from the frontal cortex, caudate and putamen. Each sample was immersed in TEM fixative (2.5% glutaraldehyde in 0.1M cacodylate buffer) and incubated overnight at 4°C. The samples were then post-fixed in 1% (w/v) osmium tetroxide (#O021, TAAB, UK) for 1.5 h at 4°C. Then they were washed and dehydrated through serial graded incubations in ethanol - 10% ethanol, followed by 70%, followed by 100%. The tissue was infiltrated in embedding resin medium (#T028, TAAB, UK) for 4 h at room temperature. Next, the samples were embedded on flat moulds and polymerised at 70°C for 24 h. Ultrathin sections (100-120 nm) were cut on a Leica ultramicrotome (Leica microsystems, Germany) and mounted on mesh copper grids. They were contrasted with uranyl acetate for 2 minutes and with lead citrate for 1 minute. Finally, the sections were imaged on an EM-1400 (Plus) transmission microscope operated at 120 kV (JEOL USA, Inc.). For each section of each brain region (frontal cortex, caudate, putamen), 5 to 10 pictures were examined for pathological changes in the capillary and neurovascular unit. The original dimensions of the images are 3296LJx2472 pixels.

The presence of a single layer of endothelial cell surrounded by a layer of basement membrane was considered to be an intact capillary. The observation of oedematous space and vacuoles around the vessels, vacuolated pericytes, large vacuoles in the endothelial cells or endoplasmic reticulum swelling, as well as oedema around the capillary and multiple layers of basement membrane were considered as signs of pathological changes (46). Neurodegeneration was identified by the presence of lipofuscin granules, loss of myelin compactness, neurite degeneration and fibrillary deposits (47,48).

### Western blot studies

Post-mortem brain capillaries from neurologically healthy individuals and AD cases were used to investigate the expression of transporters. The Braak stage of the control cases was between I, and II (age-related pathology only). For the AD cases, the Braak stage was between IV and VI. See Supplementary Information about the sex, age, post-mortem delay (PMD), clinical diagnosis, and Braak stage of the individual cases (Supplementary Table 2, S1 File).

Brain capillaries isolated from healthy individuals and AD patients were used to investigate the expression levels of BBB transporters of interest by WB analysis. Brain capillaries were isolated after homogenising brain tissue from the frontal cortex or the caudate and carrying out a dextran-based density-gradient centrifugation to produce a capillary enriched pellet. The capillary pellet was then homogenised in capillary depletion buffer (brain weight x 3) and 26% dextran (brain weight x 4). The homogenate was subjected to density gradient centrifugation (5,400 G for 15 minutes at 4°C) to give an endothelial cell-enriched pellet, the resulting supernatant was discarded (49). The pellet was further lysed in 150-200 μl ice-cold RIPA buffer with added protease inhibitors at 4°C, and then centrifuged at 8,000 G for 15 minutes at 4°C.

The protein concentration in each lysate was determined using a BCA assay and the WB procedure was carried out as already described. For antibodies used, see Supplementary Table 1, S1 File.

We used anti-TfR1 antibodies to detect TfR1 - an endothelial cell marker. This way we aimed to confirm that the capillary pellets were enriched in endothelial cells. We also used antibodies to detect GLUT1, and P-gp.

Quantification of protein expression was determined by calculating the intensity ratio of the band of interest and the band of the loading control (GAPDH). Band intensity ratio analysis was conducted using ImageJ software (35).

For statistical analysis, we used two-tailed unpaired Student’s t-test to compare the difference of transporter expression in the frontal cortex and the caudate between control and AD cases.

## Results

### Physicochemical characteristics of amisulpride and glucose

We determined the physicochemical characteristics of amisulpride and D-glucose using DrugBank and MarvinSketch. Amisulpride (chemical abstracts service (CAS) number 71675-85-9) has a MW of 369.48 g/mol, a pKa of 9.37 and exists as two microspecies at physiological pH. The predominant (96.77%) microspecies is positively charged and has a single positive charge at pH 7.4. The other microspecies (3.23%) has no charge (Fig 1). The gross charge distribution at pH 7.4 of amisulpride is +0.968.

**Fig 1.**
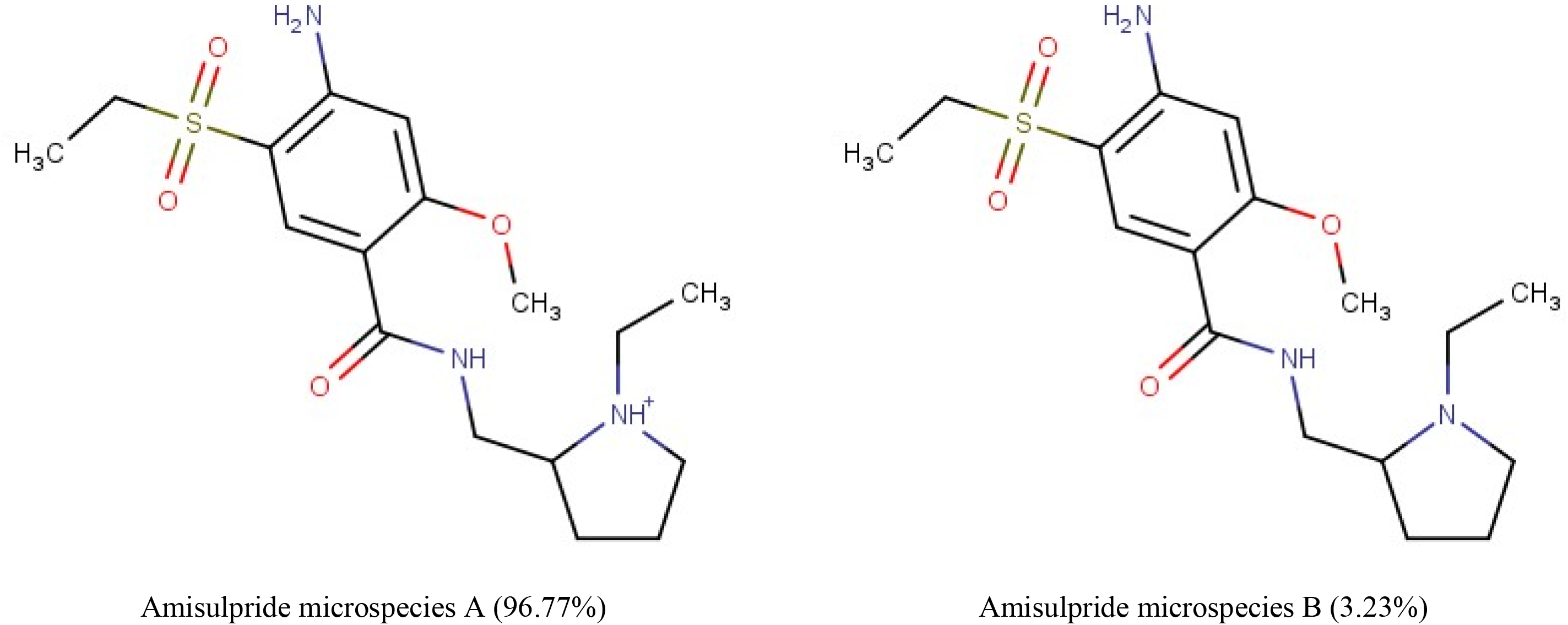
The percentage distribution and chemical structures of the two amisulpride microspecies found at physiological pH. Microspecies A has a single positive charge and Microspecies B has no charge, according to MarvinSketch 22.9.0.

D-glucose has a MW of 180.16 g/mol, and a pKa of 11.8. In solution, at equilibrium, D-glucose exists as two anomers that interconvert spontaneously: ∼36% alpha-D-glucose, and ∼64% beta-D-glucose, with less than 0.01% being present as an open-chain form (linear glucose) (32,50). The major microspecies (99.99%) of both alpha-D-glucose and beta-D-glucose has no charge (Supplementary Table 2, S1 File). The other microspecies (0.01%) of each anomer has a single negative charge at physiological pH. The linear form of D-glucose was found to be 100% neutral. Only alpha- and beta-D-glucose have been included in the GLUT1 substrates group (Supplementary Table 2, S1 File).

### Identification of GLUT1 substrates and inhibitors

Our two PubMed searches identified 99 reviews, and 16 primary research articles, which discussed GLUT1 substrates, GLUT1 inhibitors or antipsychotics which interacted with GLUT transporters and matched our inclusion criteria. In these reviews and primary research articles, 9 GLUT1 substrates, 33 GLUT1 inhibitors, and 11 GLUT-interacting antipsychotics (including 6 typical antipsychotics and 5 atypical antipsychotics) were described and are listed in Supplementary Table 3-5, S1 File. They include the typical antipsychotics: chlorpromazine, fluphenazine, loxapine, pimozide and spiperone. Haloperidol is also included although some studies suggest no interaction with GLUT (23)(51). Atypical antipsychotics include: clozapine, desmethylclozapine, olanzapine, quetiapine and risperidone. Supplementary Table 6 (S1 File) provides specific details about the experimental evidence that indicates that these 11 antipsychotics interact with GLUT. In all cases this interaction was considered to be inhibitory.

### Physicochemical characteristics of published substrates and inhibitors of GLUT1

#### Molecular Weight

The physiochemical characteristics of the identified GLUT1 substrates, GLUT1 inhibitors and the antipsychotics that interact with GLUT are summarised and compared to amisulpride (Fig 2, Table 1, and Supplementary Table 3-5, S1 File). The MW range for the GLUT1 substrates was 164.16 g/mol to 232.28 g/mol, with a mean±SEM of 184.95±6.45 g/mol, the MW range of the GLUT1 inhibitors was 180.16 g/mol to 518.55 g/mol with a mean±SEM of 325.50±14.4 g/mol and the MW range for GLUT interacting antipsychotics was 312.44 to 461.55 g/ mol with a mean±SEM of 369.38±16.04 g/mol (Table 1). One-way ANOVA showed differences in the MW F (2, 50) = 18.72, Tukey’s multiple comparisons test showed significant difference between the MW of GLUT1 substrates and inhibitors (p<0.0001), and between GLUT1 substrates and antipsychotics (p<0.0001) (Fig 2A). No significant difference was observed between the MW of the GLUT1 inhibitors and the group of antipsychotics (Fig 2A).

**Fig 2.**
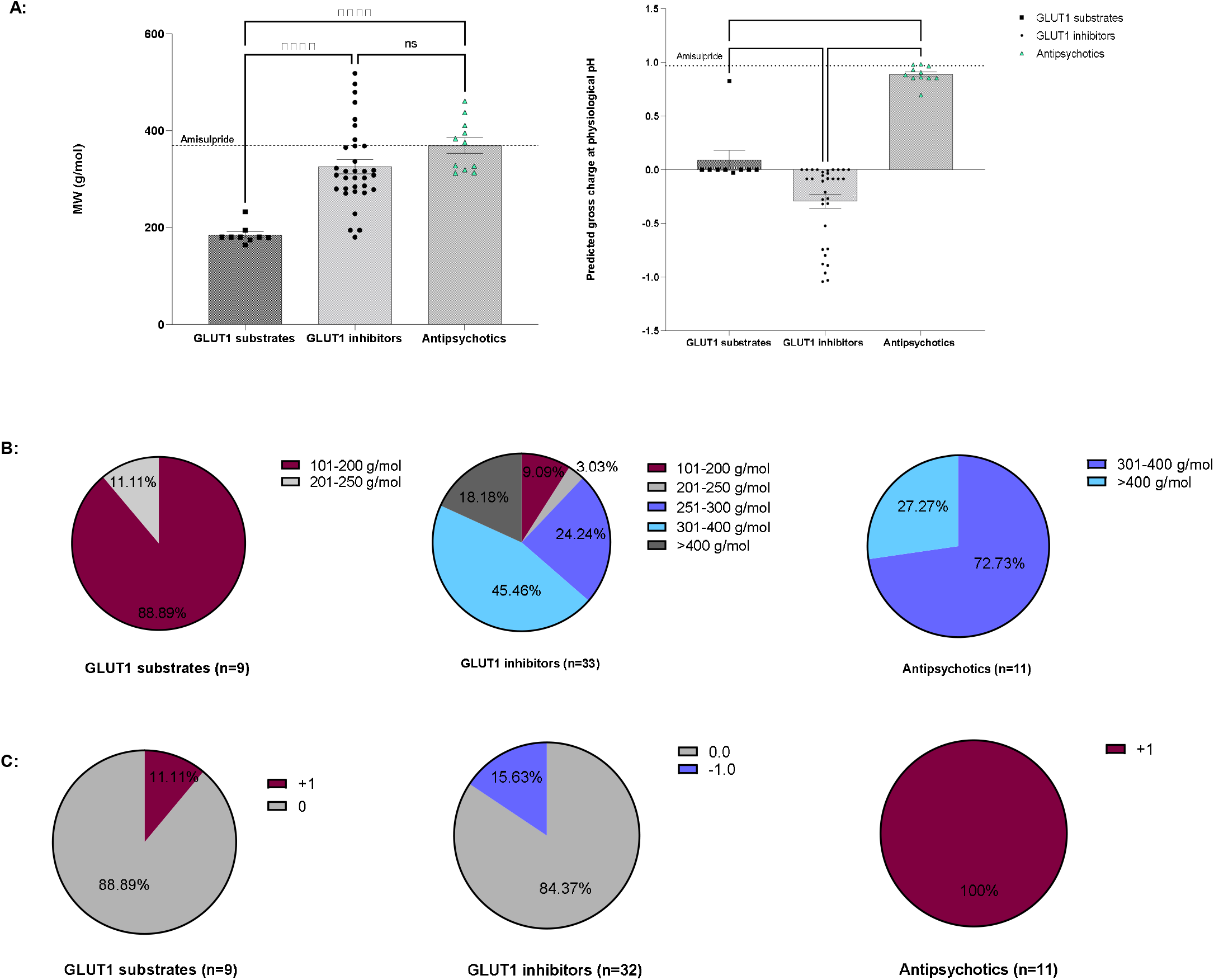
Physicochemical characteristics of GLUT1 substrates and inhibitors. Evaluation of the PubMed database identified 9 substrates and 33 inhibitors of GLUT1, and 11 antipsychotics inhibiting cell entry of GLUT1 substrates. A) Comparison of the molecular weight (g/mol) of substrates, inhibitors of GLUT1, and antipsychotics. Each square, dot, and triangle represents a compound. Comparison of the predicted gross charge of substrates, inhibitors of GLUT1, and antipsychotics at pH=7.4, according to MarvinSketch 22.9.0. Data was analysed using One-way ANOVA, GraphPad Prism 9. B) The pie charts show the molecular weight of the substrates and inhibitors of GLUT1, and of antipsychotics inhibiting cell entry of GLUT1 substrates. C) The pie charts show the charge of the most prevalent microspecies of the GLUT1 substrates, inhibitors, and antipsychotics interacting with GLUT at physiological pH according to MarvinSketch 22.9.0.

**Table 1:** Summary of the MW range, Mean±SEM, Median (g/mol), and gross charge at physiological pH of GLUT1 substrates, inhibitors, and antipsychotics interacting with GLUT.

|  | GLUT1 substrates<br>n=9 | GLUT1 inhibitors<br>n=33 | Antipsychotics<br>n=11 |  |  |
| --- | --- | --- | --- | --- | --- |
|  |  |  | All | Typical n=6 | Atypical n=5 |
| **MW Range (g/mol)** | 164.16-232.28 | 180.16-518.55 | 312.44-461.55 | 318.86-461.55 | 312.44-410.49 |
| **Mean ± SEM (g/mol)** | 184.95±6.45 | 325.50±14.4 | 369.38±16.04 | 386.18±23.42 | 349.21±20.14 |
| **Median (g/mol)** | 180.16 | 308.34 | 375.86 | 385.76 | 326.82 |
| **Gross charge at physiological pH** | +0.089<br>± 0.09 | -0.30<br>±0.06 | +0.89<br>±0.02 | +0.90<br>±0.02 | +0.87<br>±0.05 |

The median for the GLUT1 substrates was 180.16 g/mol, for the GLUT1 inhibitors it was 308.34 g/mol, for the antipsychotics it was 375.86 g/mol (Table 1). We found that 88.89% of the GLUT1 substrates were small molecules with a MW between 101 and 200 g/mol (Fig 2B). The GLUT1 inhibitors showed a wider range of molecular weight with the majority falling (69.7 %) between 250 and 400 g/mol (Figure 2B). The majority (72.73%%) of the antipsychotics affecting GLUT1 substrates uptake were in the range of 301-400 g/mol (Fig 2B). There was no significant difference in the molecular weight between the typical (386.18±23.42) and the atypical antipsychotics (349.21±20.14) (t=1.169, df=9, p= 0.27, two-tailed unpaired t-test).

#### Charge

We investigated the predicted gross charge distribution at physiological pH of the identified substrates and inhibitors of GLUT1 as well as the GLUT interacting antipsychotics (Fig 2A), and also reported the predicted charge and the percentage distribution of the top two microspecies (Supplementary Table3-5, S1 File).

Overall, we found that GLUT1 substrates, GLUT1 inhibitors and GLUT-interacting antipsychotics had an average gross charge distribution at pH 7.4 of +0.089**±**0.09, -0.30**±**0.06, and +0.89**±**0.02, respectively (Fig 2A, and Table 1). There was a significant difference between all three groups (One-way ANOVA F (2,50) = 58.96, p<0.0001, Tukey’s multiple comparisons test GLUT1 substrates vs GLUT1 inhibitors (p=0.0057); GLUT1 substrates vs antipsychotics p<0.0001; GLUT1 inhibitors vs antipsychotics p<0.0001). There was no significant difference in the gross charge between the typical (0.90±0.02) and the atypical (0.87±0.05) antipsychotics (t=0.678, df=9, p= 0.5147, two-tailed unpaired t-test).

Importantly, there were two molecules which could be outliers within their groups. This included the GLUT1 substrate, glucosamine, which had a much greater charge of +0.827, and the atypical antipsychotic, quetiapine, which had a lower charge of +0.696 (Fig 2A).

GLUT1 substrates existed as either one, two or three microspecies at physiological pH. GLUT1 inhibitors could exist as non-ionizable molecules (e.g. mercuric chloride) or as up to 25 microspecies (e.g. morin) at physiological pH. The antipsychotics suggested to interact with GLUT existed at physiological pH as either two, three or four microspecies. This information plus the predicted charge of the top two microspecies (if present) at physiological pH is tabulated in Supplementary Tables 3-5, S1 File. The charge of the major microspecies of each group is presented in the form of pie charts (Fig 2C). The major microspecies of GLUT1 substrates were neutral or had +1 charge, GLUT inhibitors were neutral or had -1 charge and the anti-psychotics which interacted with GLUT all had +1 charge at physiological pH.

### *In silico* molecular docking

*In silico* molecular docking studies revealed that amisulpride could interact with GLUT1 based on the low free energy binding of the interaction: -29.04 kcal/mol, and the high chem score: 26.79. The molecular docking predicted conventional hydrogen bonds between oxygen atoms from amisulpride and the amino acids: threonine (THR) A: 137, and tryptophan (TRP) A: 412, and between hydrogen atom in the NH_2_ group of amisulpride and asparagine (ASN) A: 411. Alkyl interaction between amisulpride and the amino isoleucine (ILE) A: 164, and pi-pi stacking interaction between amisulpride and TRP A: 412 were also observed (Fig 3A).

**Fig 3.**
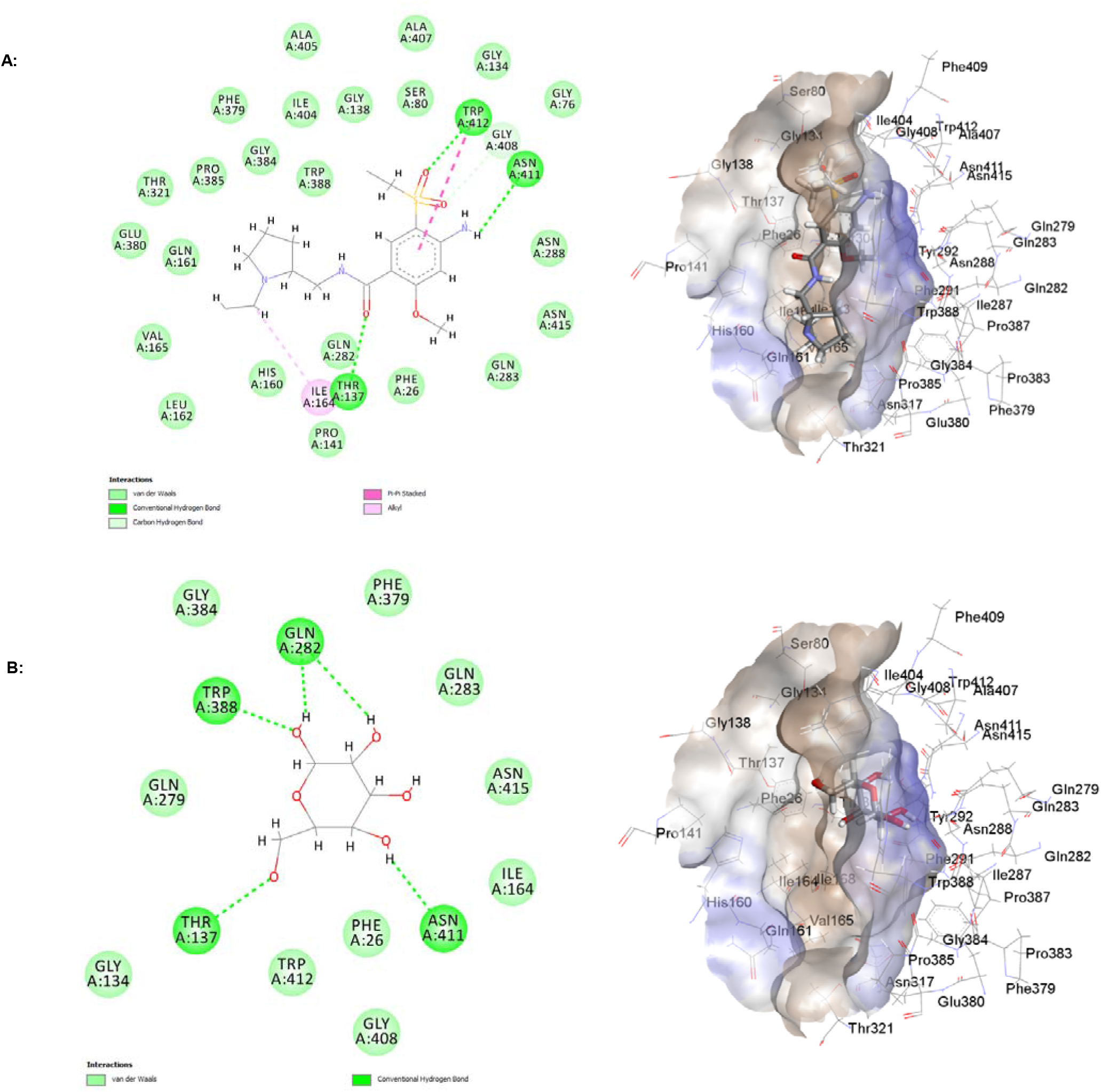
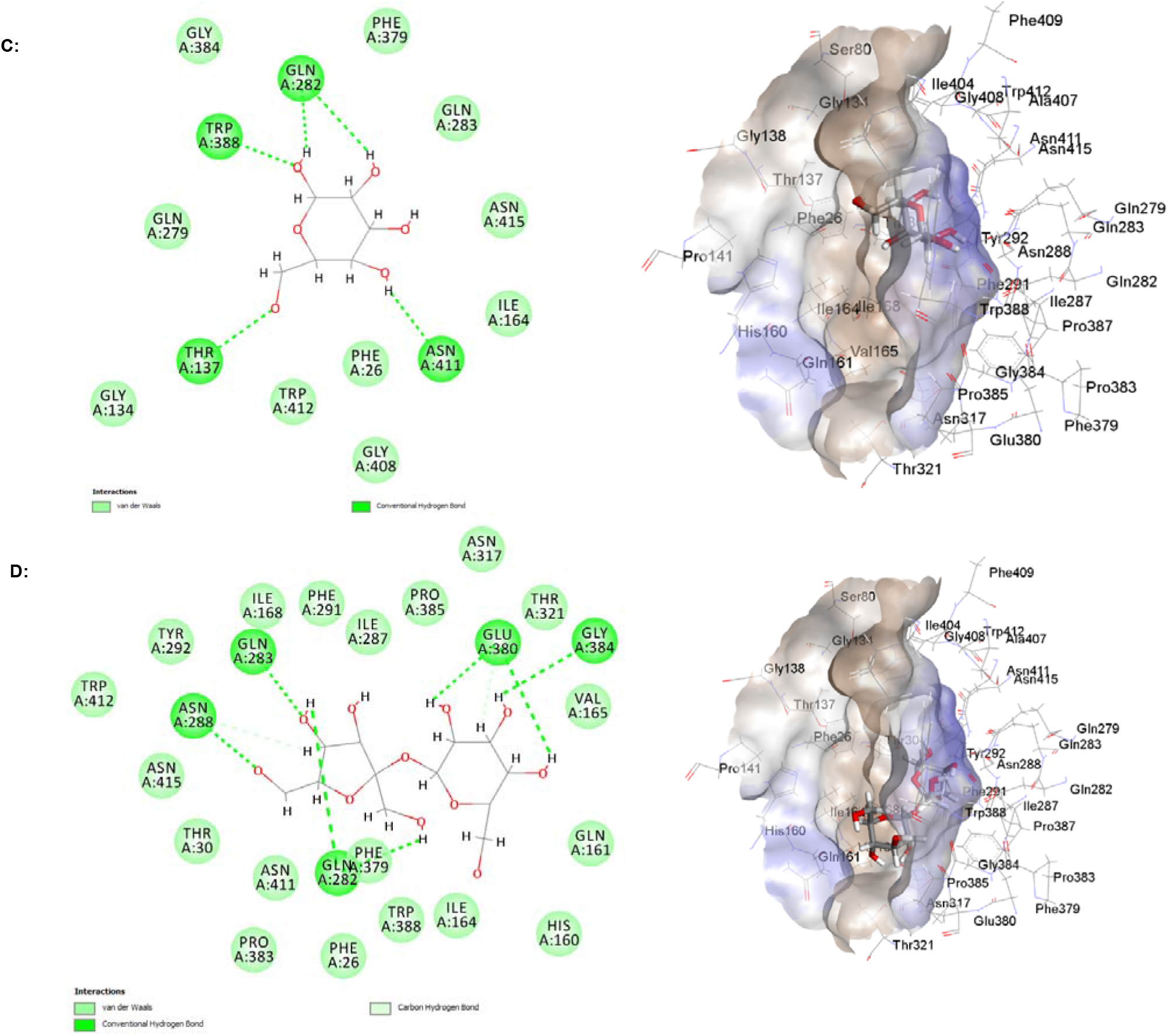
Molecular level interactions of amisulpride, alpha-D-glucose, beta-D-glucose, and sucrose with the binding sites of GLUT1. 2D (left) and 3D (right) representations. In the 2D representations, green dotted lines are used to show hydrogen bonds, pink dotted lines are used to depict hydrophobic interactions. A) Stick representation is used for amisulpride, and line representation for the amino acid residues from GLUT1. B) Stick representation is used for alpha-D-glucose, and line representation for the amino acid residues from GLUT1. C) Stick representation is used for beta-D-glucose, and line representation for the amino acid residues from GLUT1. D) Stick representation is used for sucrose, and line representation for the amino acid residues from GLUT1.

We also studied the interaction of GLUT1 with alpha-D-glucose and beta-D-glucose. This interaction was used as a positive control. Molecular docking study found that both alpha-D-glucose, and beta-D-glucose are substrates for GLUT1 with free binding energy of -15.39 kcal/mol and chem score of 15.29. Conventional hydrogen bonds between alpha-D-glucose and beta-D-glucose, and GLUT1 were observed at TRP A: 388, glutamine (GLN) A: 282, ASN A: 411, and THR A: 137 (Fig 3B and 3C). The negative control sucrose, a disaccharide, showed higher free binding energy and lower chem score than amisulpride, and alpha- and beta-D-glucose (monosaccharide). The free energy binding for the interaction of sucrose with GLUT1 was -8.58 kcal/mol, and the chem score was 6.55 (Fig 3D).

### Expression and Function of GLUT1 in hCMEC/D3 cells

We examined the expression and function of GLUT1 in an established cell model of the human BBB - the hCMEC/D3 immortalised cell line. Verification studies of the experimental design were also performed.

#### Expression of GLUT1 in hCMEC/D3 cells

The hCMEC/D3 cells were found to express the GLUT1 transporter (40-60 kDa). GAPDH was used as a loading control. Lysates from human colon adenocarcinoma (Caco-2) cells were used as a positive control, and lysate from the human lung cancer cell line (Calu-3) or human embryonic kidney cell line (HEK-293) were used as a negative control (Fig 4A).

**Fig 4.**
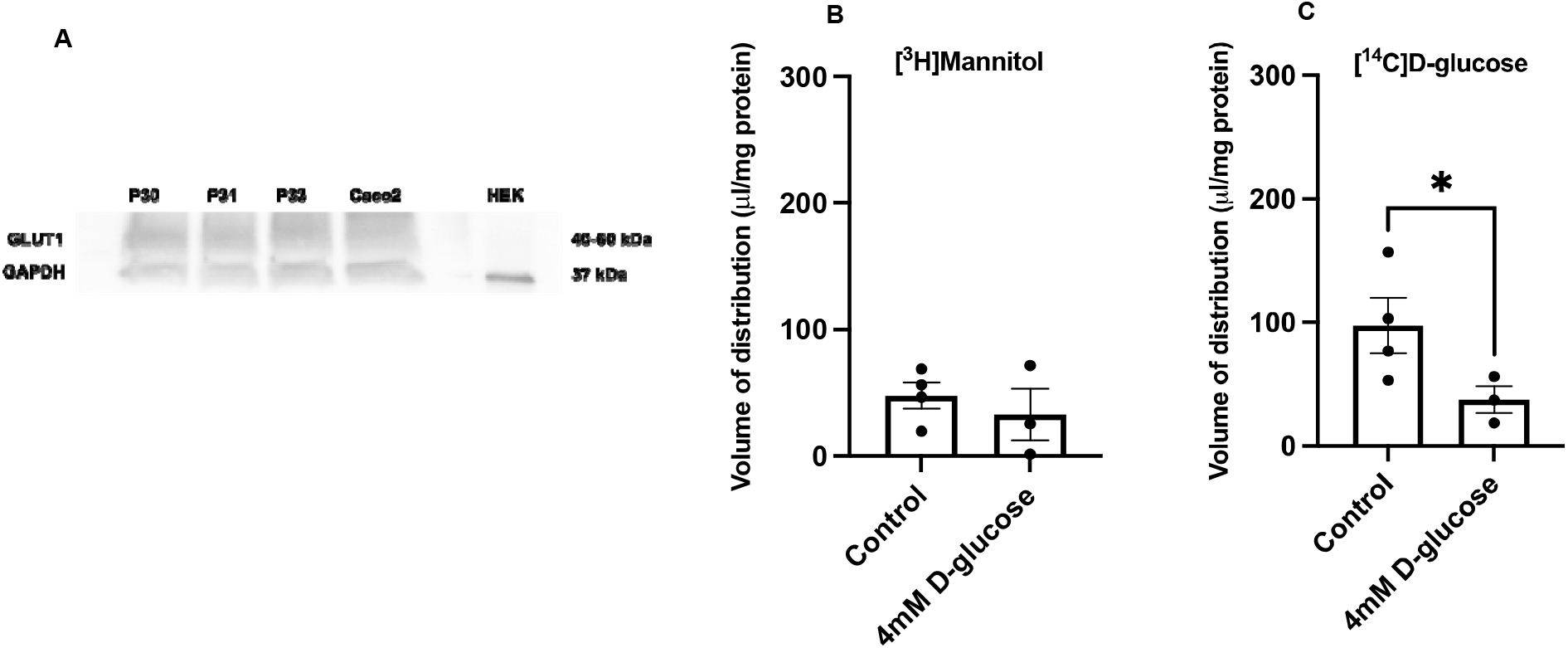
Function of GLUT1 in hCMEC/D3 cells – self inhibition. A) GLUT1 expression in hCMEC/D3 cells. Three passages of hCMEC/D3 cells (P30, 31, 33) (30 µg of protein per well) were tested for GLUT1 (40-60 kDa) expression. The figure is an example membrane of three technical repeats. Caco-2 cell lysate was used as a positive control; HEK-293 cell lysate was used as a negative control. GAPDH (37 kDa) was used as a loading control. Antibodies used: anti-GLUT1 antibody – 1:100 000, #ab115730; anti-GAPDH antibody – 1:2500, #ab9485, Abcam; secondary anti-rabbit IgG, HRP-linked antibody – 1:2000, #7074, Cell Signalling Technology. B) V_d_ of [^3^H]mannitol was not significantly different between the control and the experimental conditions. C) Non-labelled glucose decreased significantly the accumulation of [^14^C]D-glucose ([^3^H]mannitol corrected) in hCMEC/D3 cells after 1 h of incubation. Results are expressed as mean ± SEM, n=3-4 plates, passages (P33, 34, and 35) with five well replicates per treatment in each plate. Data were analysed with an unpaired one-tailed Student’s t-test, using GraphPad Prism 9, each point represents a plate.

#### Accumulation assays – function of GLUT1 in hCMEC/D3 cells

To check that excess concentrations of D-glucose did not affect the membrane integrity of the hCMEC/D3 cells, verification studies were performed. Cells were incubated with [^14^C]D-glucose and [^3^H]mannitol as a control (containing the standard amount of non-labelled D-glucose – 1.1 mM), and with [^14^C]D-glucose, [^3^H]mannitol, and non-labelled D-glucose (4 mM) as a test condition. There was no significant change in the permeability of [^3^H]mannitol (cell permeability marker) when the cells were treated with 4 mM non-labelled D-glucose for 1 h (Fig 4B). Further analysis and studies could therefore be performed using similar D-glucose concentrations.

To investigate the function of GLUT1 in hCMEC/D3 cells, we performed a self-inhibition experiment. Incubation of the cells with 4 mM non-labelled D-glucose for 1 h led to a significant decrease in the V_d_ of [^14^C]D-glucose ([^3^H]mannitol and protein corrected) (from 97.5 ± 22.3 µl/mg to 37.3 ±10.8 µl/mg; t=2.161, df=5, p= 0.0415, unpaired one-tailed Student’s t-test, data is presented as mean ± SEM) (Fig 4C).

When we tested the interaction of amisulpride with GLUT1 in hCMEC/D3 cells, the entry of [^3^H]mannitol (0.05 µM) into the cells was used as a cell integrity marker. There was no significant change in the [^3^H]mannitol permeability when the cells were treated with amisulpride (20, 50, or 100 µM) (Fig 5A). There was also no significant effect on the V_d_ of [^14^C]D-glucose when the cells were treated with amisulpride (20, 50, or 100 µM) (Fig 5B).

**Fig 5.**
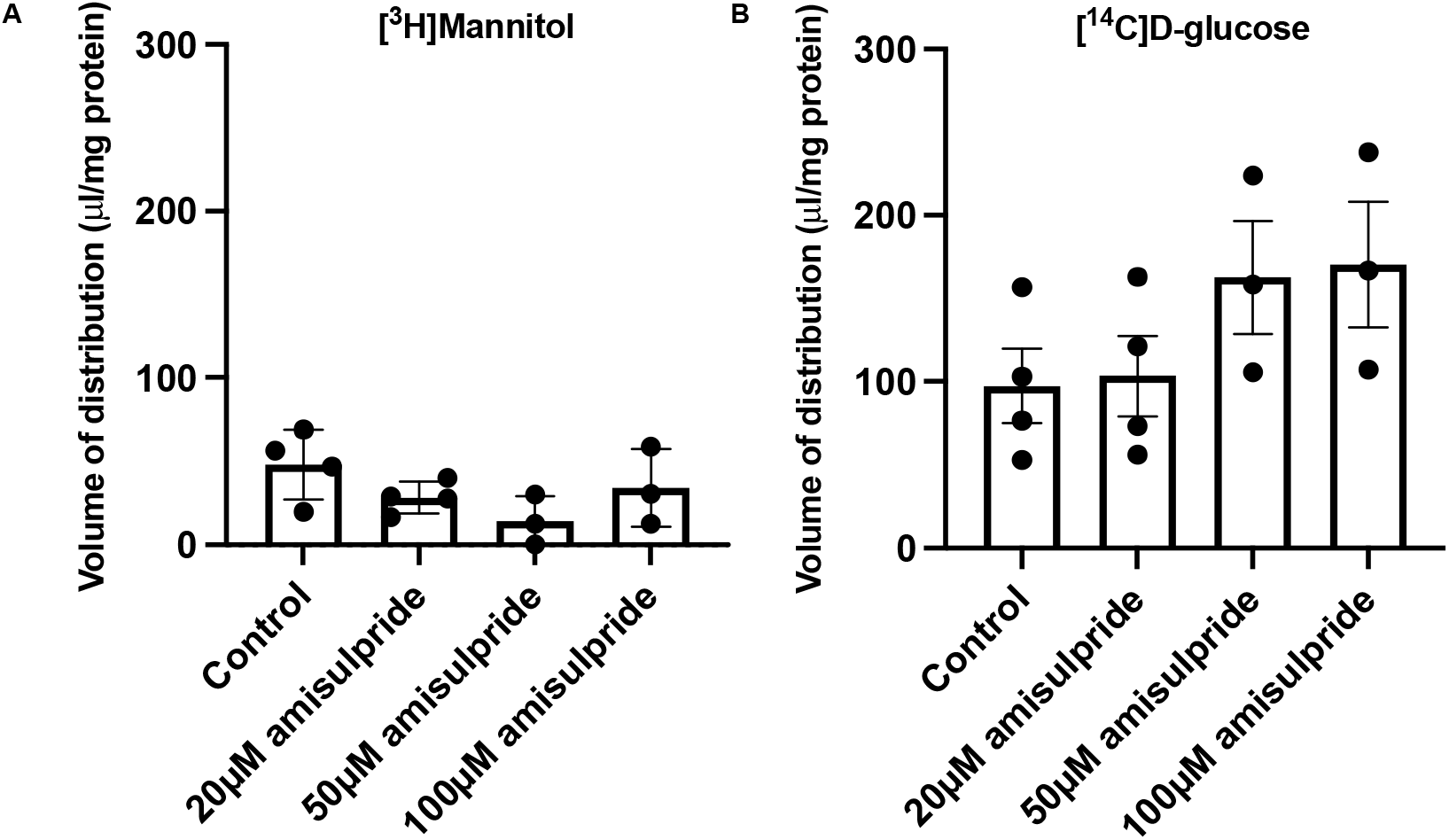
Interaction of amisulpride with GLUT1 in hCMEC/D3 cells. A) Amisulpride at three different concentrations (20, 50, or 100 µM) did not have a significant effect on the accumulation of [^3^H]mannitol in hCMEC/D3 cells after 1 h of incubation. B) Amisulpride at three different concentrations (20, 50, or 100 µM) did not have a significant effect on the accumulation of [^14^C]glucose ([^3^H]mannitol corrected) in hCMEC/D3 cells after 1 h of incubation. Results are expressed as mean ± SEM, n=3-4 plates, passages (P33, 34, and 35) with five well replicates per treatment in each plate. Data were analysed with a One-way ANOVA, using GraphPad Prism 9, each point represents a plate.

### Animal model of AD - validation

#### Mouse weight comparison

We compared the weight of the WT and 5xFAD mice and of the two sexes (age between 12 and 15 months). All data are presented as mean ± SEM. The WT mice had higher average weight than the 5xFAD mice (32.31 ± 2.93 g vs 26.15 ± 2.5 g), and all the 5xFAD mice with weight lower than 25 g were female. We observed that the females had significantly lower weight (g) than the males in the 5xFAD group (19.7 ± 1.46 vs 31 ± 1.61; t=4.983, df=5, p=0.0042; unpaired two-tailed Student’s t-test). In the WT group, weight difference between the sexes was smaller and did not reach significance (Supplementary Fig 3, S1 File).

#### TEM ultrastructural study

We examined frontal cortex samples of WT and 5xFAD mice for the presence of Aβ plaques using TEM. Structures were identified as Aβ plaques by comparison to Aβ plaques which had previously been identified in TEM images (40). Amyloid plaques were not observed in the frontal cortex of the young WT mice (4.5-6 months; weight 25.4 g, 26.1 g). Importantly, structures identical to Aβ plaques previously reported in the literature (40) were present in the 5xFAD mice in both the young (6 months; weight 23.8 g, 23.5g) and the old (12 months 21.9 g, 22 g, data not shown) group (Fig 6).

**Fig 6.**
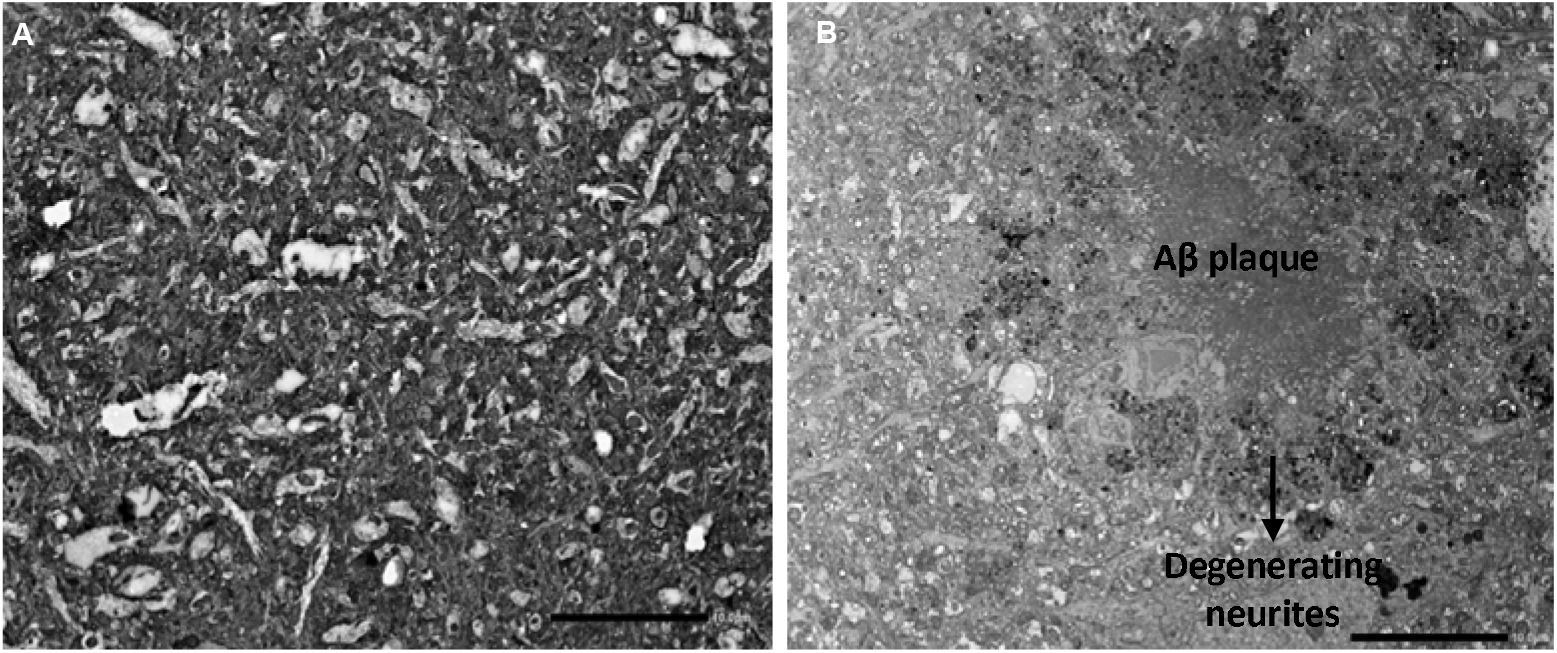
TEM ultrastructural study in WT and 5xFAD mice. TEM image of the frontal cortex of A) WT mouse, which is free of Aβ plaques, and B) 5xFAD mouse, with Aβ plaques. Mouse age – 4.5 – 6 months. Magnification – 1200x, scale bar – 10 µm, n=2 mice per group.

#### Western blot studies

We performed WB for human APP to examine the genotype of each mouse used in this study. APP is the precursor protein from which amyloid-β (Aβ) is cleaved by β-secretase and γ-secretase (52). Our studies confirmed human APP expression was not detectable in the WT mice, but was detectable in the 5xFAD mice cerebral capillaries (See Supplementary Fig 4, S1 File)

We observed TfR1 and transporter expression in the mouse brain endothelial cell lysates from WT and 5xFAD mice used in the *in situ* brain perfusions (See Supplementary Fig 5, S1 File). Thus, these experiments confirmed we have successfully isolated the BBB compartment from the whole brain.

In particular, brain capillaries from WT and 5xFAD mice were found to express GLUT1, PMAT, multi-drug and toxin extrusion protein 1 (MATE1), OCT1, and P-gp. No significant effect of the genotype was found on the expression of these proteins (See Supplementary Figs 5-9, S1 File).

### *In situ* brain perfusions

[^3^H]amisulpride uptake (uncorrected for [^14^C]sucrose) was significantly greater than [^14^C]sucrose uptake in all brain areas studied apart from the hypothalamus, thalamus, and capillary pellet in the WT mice (paired two-tailed t-test, data not shown). In the 5xFAD mice the [^3^H]amisulpride uptake (uncorrected for [^14^C]sucrose) was significantly greater than [^14^C]sucrose uptake in all brain areas studied (paired two-tailed t-test, data not shown).

When comparing WT and 5xFAD mice, there was a trend for an increase in [^3^H]amisulpride uptake into specific brain regions, whole brain homogenate, and capillary pellet in the 5xFAD mice but this failed to reach statistical significance (Fig 7A and 7B). However, there was a statistically significant increase in the uptake of radiolabelled amisulpride in the supernatant (made of brain parenchyma and interstitial fluid) of 5xFAD mice compared to WT mice (t=2.550, df=8, p=0.0342, unpaired two-tailed t-test, by 104.86%). These results were [^14^C]sucrose corrected. None of the regions showed a significant difference in the [^14^C]sucrose uptake between the WT and the 5xFAD mice (Fig 7A and 7B).

**Fig 7.**
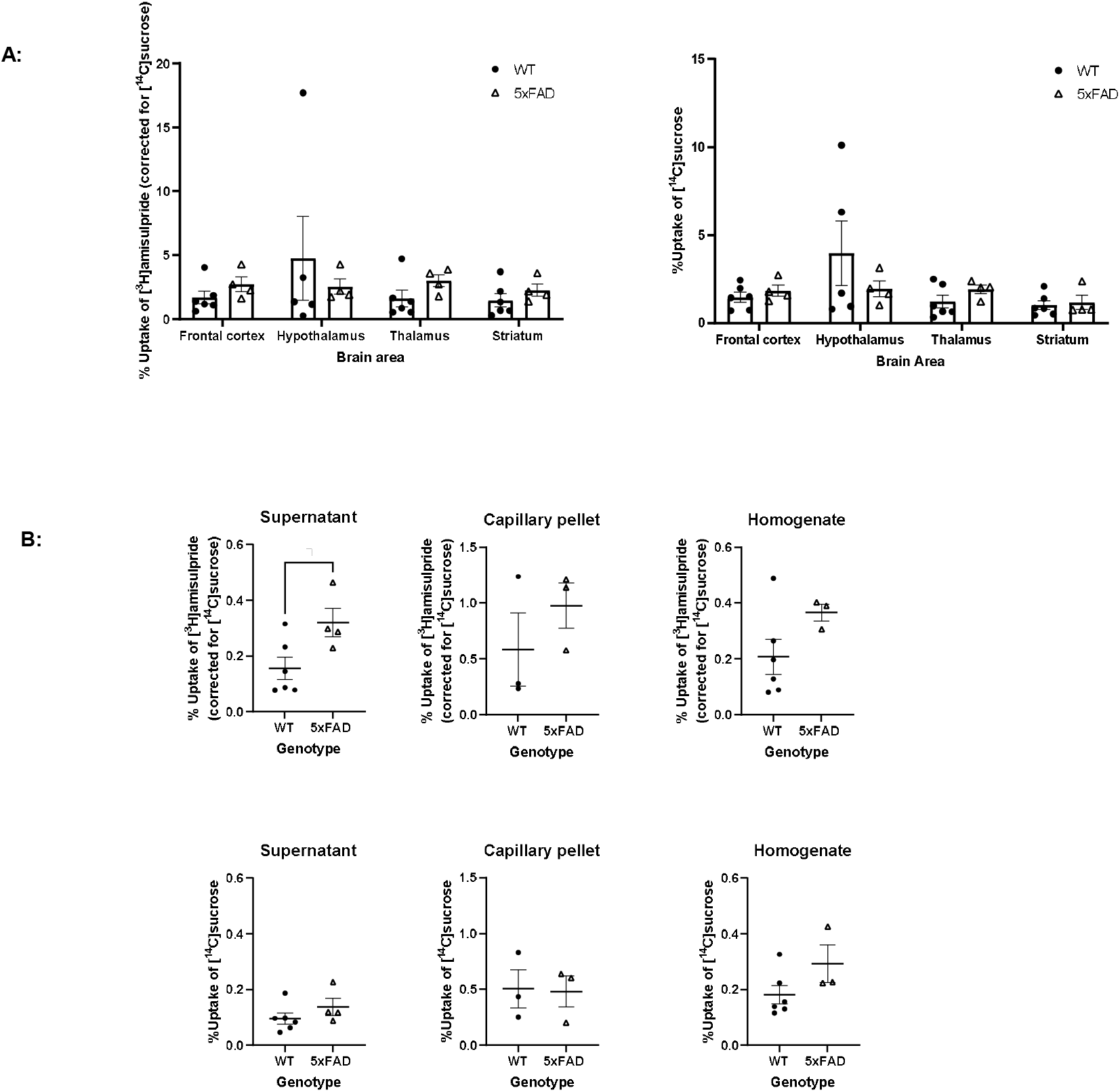
[^3^H]Amisulpride (sucrose corrected) and [^14^C]sucrose uptake into the brain of WT and 5xFAD mice. WT and 5xFAD mice were perfused with [^3^H]amisulpride and [^14^C]sucrose. Perfusion time - 10 minutes, fluid flow rate – 5.5 ml/min. Mouse age: 12-15 months. WT n=6, 5xFAD n=4, apart for the hypothalamus where WT n=5, 5xFAD n=4; the capillary pellet where WT n=3, 5xFAD n=3, and the homogenate where WT n=6, 5xFAD n=3. No significant differences in paracellular permeability and membrane integrity were observed in any region. Each dot represents data from one mouse. All data are expressed as mean ± SEM, Mixed effects analysis and unpaired two-tailed Student’s t-test, GraphPad Prism 9.

### Human control and AD tissue studies

#### TEM ultrastructural study

For each brain region (frontal cortex, caudate, putamen), 10 to 20 pictures were examined, and the endothelial cells, basement membranes, capillary lumens, axons, myelin sheets and markers of degeneration were identified and labelled with reference to Electron Microscopic Atlas of cells, tissues and organs (43). We observed thickened basement membrane, vacuolisation of the endothelial cell, fibrillary deposits, and oedema around brain capillaries in the frontal cortex, caudate and the putamen (Fig 8). Various other features of degeneration were observed in our AD case (BNE stage VI), including: lipofuscin granules in the frontal cortex and the putamen, and myelin degeneration in the caudate (Fig 9).

**Fig 8.**
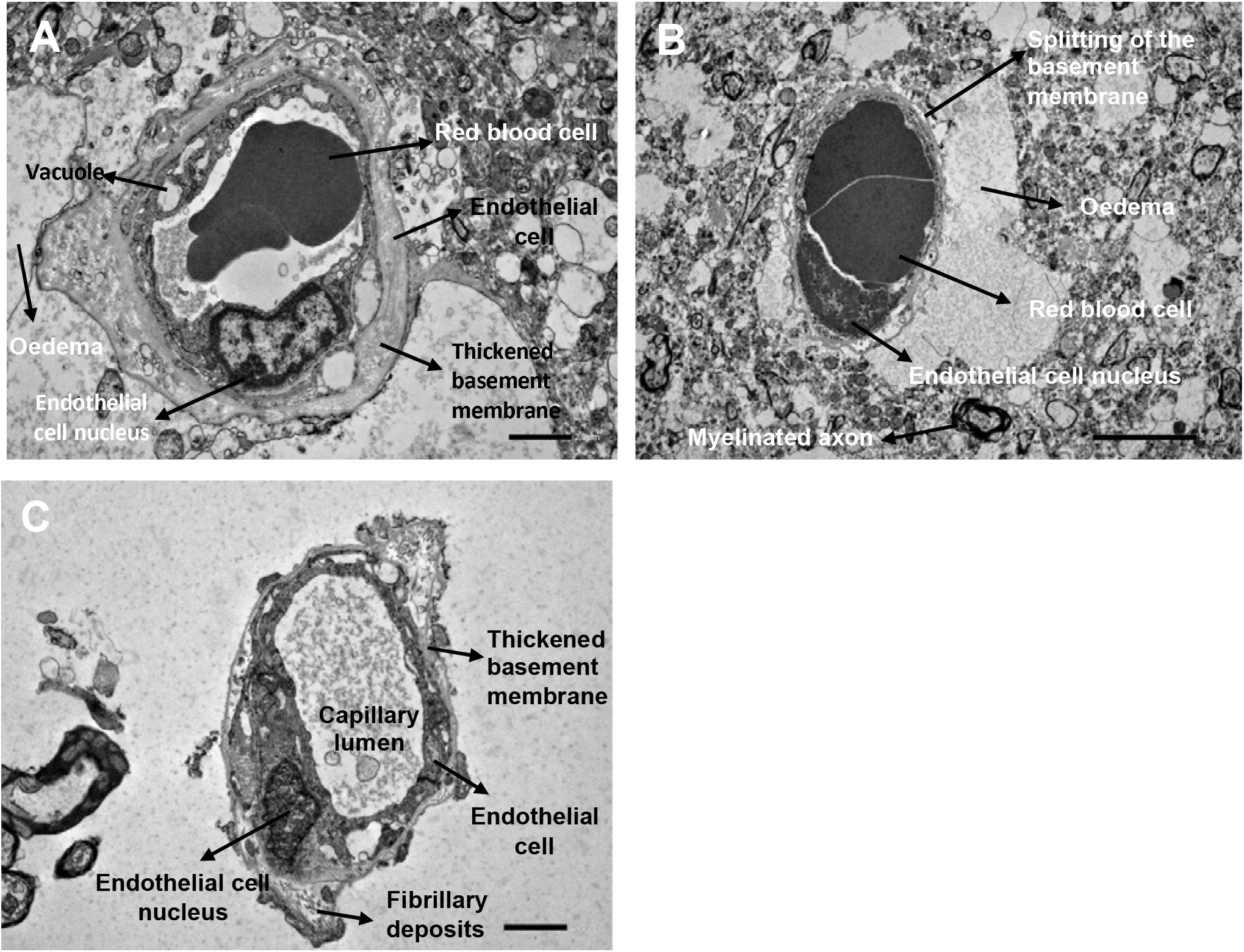
TEM image of capillaries in the frontal cortex (A) putamen (B) and caudate (C) of a human AD case. Frontal cortex and caudate magnification – 3000x scale bar – 2 µm. Putamen magnification – 2000x, scale bar – 5 µm. n=1 case, two sections per brain area were stained and imaged, 5 to 10 images were examined per section. Case number: BBN002.32856; Sex: F; Age: 74; PM delay: 19 h; Alzheimer’s disease, BNE stage VI.

**Fig 9.**
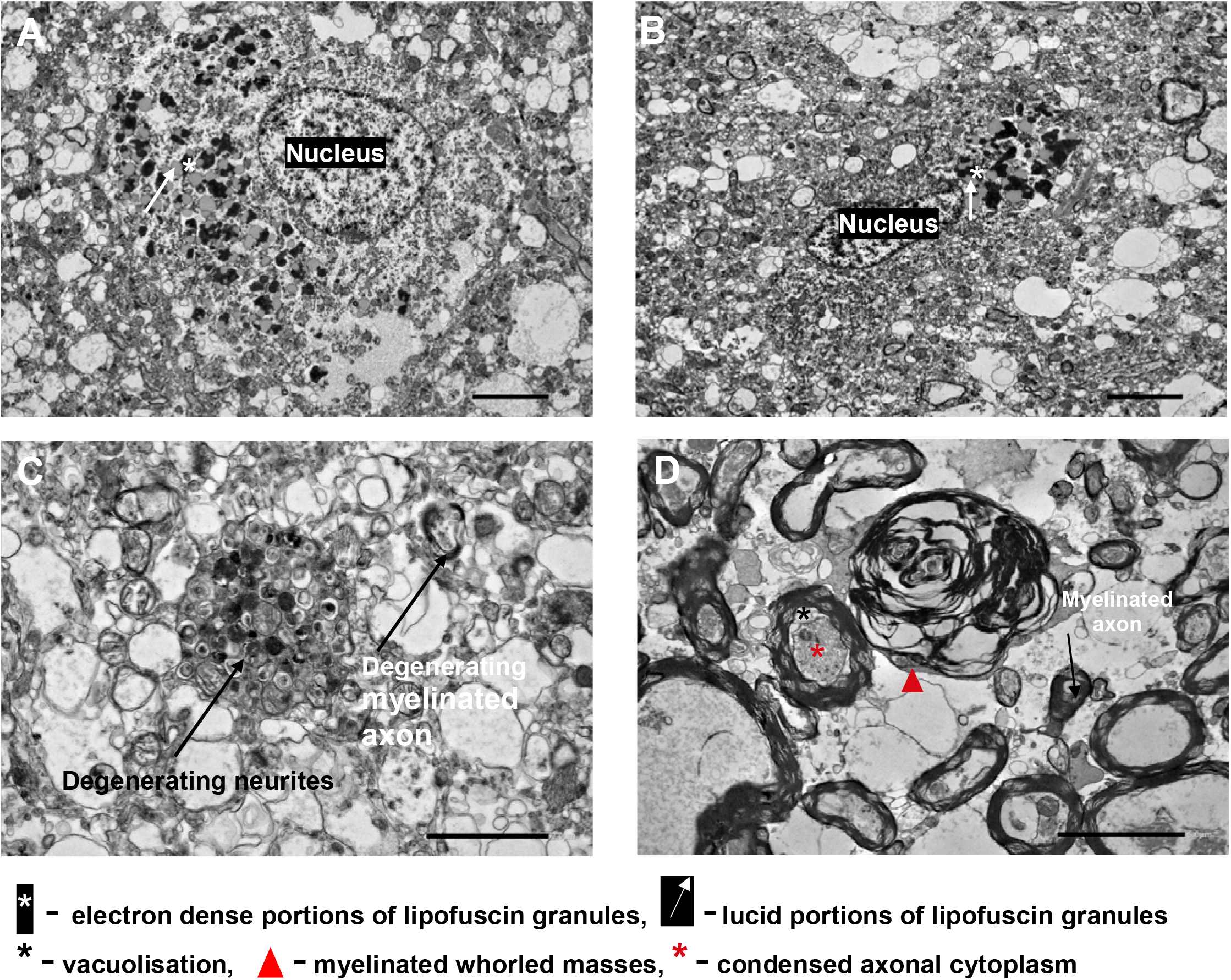
TEM image of the A) frontal cortex (A and C), putamen (B), and caudate (D) of human AD case. A) TEM image of the frontal cortex of human AD case. B) TEM image of the putamen of human AD case. A and B showing electron dense and lucid portions of lipofuscin granules. Magnification – 1500x, scale bar – 5 µm. C) TEM image of degenerating neurites and degenerating myelinated axon in the frontal cortex of a human AD case. Magnification – 6000x, scale bar – 2 µm. D) TEM image of myelin and axon degeneration in the caudate of a human AD case. Magnification – 2500x, scale bar – 5 µm. Case, n=1, two sections per brain area were stained and imaged, 5 to 10 images were examined per section. Case number: BBN002.32856; Sex: F; Age: 74; PM delay: 19 h; Alzheimer’s disease, BNE stage VI.

#### Total protein concentration

A BCA assay was used to compare the total protein concentration in the frontal cortex and caudate capillary lysates of control vs AD cases. For the frontal cortex control cases n=9; AD cases n=9) and caudate (control cases n=9; AD cases n=5), an unpaired two-tailed t-tests showed no significant difference between AD and control (Supplementary Fig 10, S1 File).

#### Western blot studies

TfR1 was detected in the capillary protein lysates from the frontal cortex and caudate of human brains both in the control and in AD cases. There was no significant difference between the two groups in the frontal cortex or the caudate (unpaired two-tailed t-test). The expression of TfR1 in the samples confirms they are endothelial cell enriched (Fig 10).

**Fig 10:**
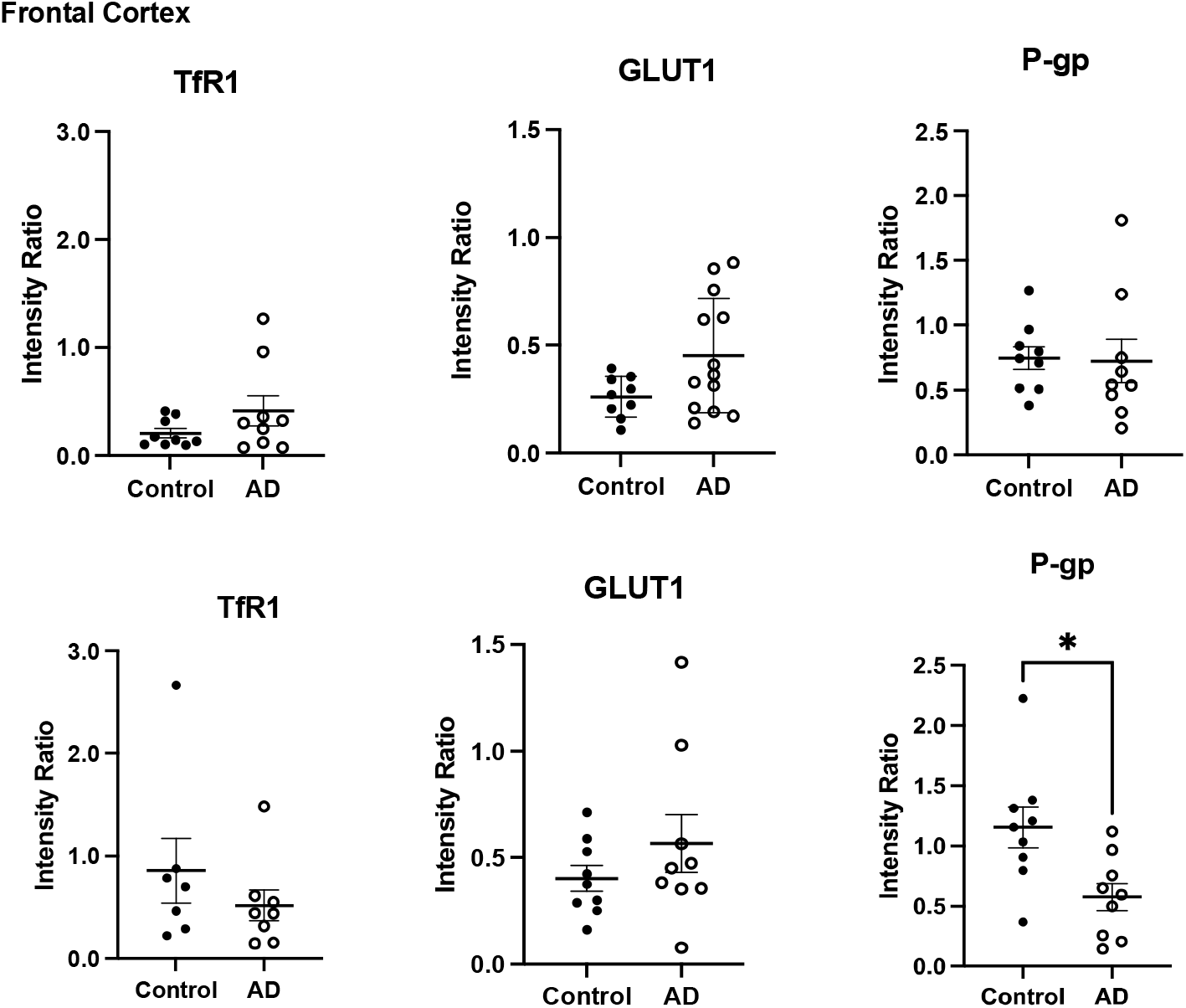
TfR1, GLUT1 and P-gp expression in human frontal cortex and caudate brain capillary lysates. Comparing control and AD expression. Intensity ratio was calculated using the intensity of the band of TfR1, GLUT1 or P-gp, and controlling it for the intensity of the loading control GAPDH. Results are presented as mean±SEM, each dot represents a case, data was analysed using unpaired two-tailed Student’s t-test. All analysis was performed using Image J, Excel, and GraphPad Prism 9.

We studied the expression of GLUT1 in control and AD age-matched cases. We did not see a significant change in the expression of GLUT1 between control and AD in the frontal cortex or the caudate (Fig 10).

We also studied the expression of P-gp in control and AD age-matched cases. The brain areas we examined were the frontal cortex and the caudate. We observed a decrease in P-gp expression in the AD group comparted to control, both in the frontal cortex and the caudate. In the caudate this decrease was significant (t=2.841, df=16, p=0.0118), unpaired two-tailed Student’s t-test (Fig 10; Supplementary Figs 11-13, S1 File).

## Discussion

This study investigated the role of GLUT1 in the transport of the antipsychotic, amisulpride at the BBB, to understand better its role in the hypersensitivity of AD patients to the side effects of antipsychotics compared to healthy aged patients (4). We used an integrative approach to test the hypotheses that amisulpride interacts with GLUT1 at the BBB, and that expression of BBB transporters and the transport of amisulpride and glucose into the brain is affected by AD. Analysis of the published literature allowed us to identify three groups of molecules: GLUT1 substrates, GLUT1 inhibitors and antipsychotics that interacted with GLUT. We then utilized specialist chemical property databases to obtain a clearer picture of the physicochemical characteristics of these molecules and their groups as well as amisulpride. *In silico* docking studies allowed us to explore the specific molecular interactions of amisulpride with GLUT1. WB and *in vitro* accumulation assays in hCMEC/D3 cells were used to confirm the expression of GLUT1 protein in these BBB cells, the presence of functional GLUT1 transporter, and to determine possible interactions between GLUT1 and amisulpride. We also examined the neurovascular unit architecture in WT and 5xFAD mice at an ultrastructural level using TEM and assessed the suitability of the 5xFAD mouse model to study the BBB permeability of amisulpride in AD. WB was used to study the expression of SLC and ABC transporters, including GLUT1 at the BBB in WT and 5xFAD mice. The *in situ* brain perfusion technique allowed us to compare amisulpride uptake into compartments of the brain in WT and 5xFAD mice. We also compared transporter expression in capillaries from human cases with and without Alzheimer’s dementia. TEM was utilised to directly visualise cellular structures in an individual with AD.

### Physicochemical characteristics of GLUT1 substrates and inhibitors

Amisulpride is an atypical antipsychotic and has a MW of 369.48 g/mol. The main microspecies (96.77%) at physiological pH has one positive charge. The other microspecies (3.23%) of amisulpride at physiological pH has no charge. The gross charge distribution at pH 7.4 of amisulpride is +0.968.

Our literature review and analysis of the chemical property databases revealed that GLUT1 substrates are typically neutral, however, we have identified one substrate, D-glucosamine, which has a positive gross charge distribution physiological pH of +0.827. GLUT1 inhibitors are predominately neutral, but we identified inhibitors which had a negative gross charge distribution at pH 7.4 (for example, -1.043 - Lavendustin B).

Importantly, both typical and atypical antipsychotics were identified which impede uptake of GLUT1 substrates *in vitro.* They all have a gross charge distribution which is positive at physiological pH and their main microspecies at physiological pH has a single positive charge, similar to amisulpride. An example is olanzapine for which there is also *in silico* molecular docking data for an inhibitory interaction with a bacterial glucose/H^+^ symporter from *Staphylococcus epidermidis*. It can impede the alternating opening and closing of the substrate cavity necessary for glucose transport (53). Another example is risperidone (24), which has been reported to interact with GLUT, and to inhibit glucose uptake *in vitro* (in rat PC12 cells). Thus, it is plausible that other atypical antipsychotics such as amisulpride could interact with GLUT1.

Although it is important to consider that not all antipsychotics interact with GLUT. For example, sulpiride (CAS 23756-79-8; MW 341.43, gross charge at physiological pH +0.972) and clozapine N-oxide (a clozapine metabolite: CAS 34233-69-7; MW 342.83, gross charge at physiological pH +1.616) were reported to not have a significant effect on glucose uptake *in vitro* (23,51).

When it comes to MW, all the established substrates of GLUT1 are smaller than amisulpride – 164.16 to 232.28 g/mol (MW of the main substrate - glucose is 180.16 g/mol). Amisulpride, has a MW similar to the established GLUT1 inhibitors and to the antipsychotics reported to inhibit uptake of GLUT1 substrates. Amisulpride is therefore more likely to represent a GLUT1 inhibitor than substrate. However, this interpretation is limited by the small number of established GLUT1 substrates, even though a large number of reviews (99) were examined.

First generation antipsychotics are known to inhibit dopaminergic neurotransmission most effectively by blocking about 72% of the D2 dopamine receptors in the brain. They also block noradrenergic, cholinergic, and histaminergic receptors. Whereas, second generation antipsychotics block D2 dopamine receptors and serotonin receptors (5-HT), mainly the 5-HT2A subtype (54).

Interestingly, amisulpride (second generation antipsychotic) shows high and similar affinities for the D2 and D3 dopamine receptor subtypes but it does not have significant affinity to the other receptor subtypes (55).

Although atypical antipsychotics are usually considered to have a broader mechanism of action compared to typical antipsychotics, there were no significant differences in physicochemical characteristics of the two groups or in their reported type of interaction with GLUT1.

### In silico studies

*In silico*, GLUT1 was found to interact with amisulpride with a free energy binding of - 29.04 kcal/mol (the positive control, beta-D-glucose, showed free energy binding of - 15.39 kcal/mol, and the negative control, sucrose, showed much higher free energy binding of -8.58 kcal/mol). Previous studies have shown that interactions with TRP412, TRP388, phenylalanine (PHE) 291, PHE379 and glutamate (GLU) 380 may play critical roles in ligand binding to GLUT1 in the inward open conformation (33). Also, another study reported interactions between cytochalasin B, a competitive inhibitor of glucose exit via GLUT1 (56), and THR137, ASN411, GLN282, ASN288, glycine (GLY) 384 and TRP388 in GLUT1 (57). In addition, ASN411 and TRP412 were found to be important for the binding of inhibitors in the central binding site of GLUT1 (57). Importantly, we see interactions between amisulpride and GLUT1 at some of the reported amino acid residues – TRP412, THR137 and ASN411. The interaction of amisulpride with these specific amino acid residues in GLUT1, taken together with the size of amisulpride compared to GLUT1 substrates and inhibitors, and its positive charge, suggests amisulpride might competitively inhibit glucose delivery to the CNS via GLUT1.

Dysfunction of GLUT1 is likely to cause further dysregulation of the neurovascular unit (NVU), resulting in the loss of BBB integrity as well as the observed altered transporter expression and increased Aβ toxicity in AD. Interestingly, increased neuronal uptake of glucose has been shown to protect against Aβ toxicity, and altering glucose delivery to the brain could influence progression of AD pathology (58). A situation that may be exacerbated by the additional medications AD patients usually receive (59). Antipsychotics are commonly prescribed with antidepressant or sedative drugs (60), i.e., citalopram (2) (MW: 324.4; +1 charge at pH 7.4) (30). It can be assumed that if two of these psychotropics are substrates (or inhibitors) for the same transporter and are prescribed together then this is likely to change individual drug delivery and potentially nutrient delivery to the CNS and contribute to the drug hypersensitivity. In fact, polypharmacy is an important predictor of adverse drug reactions for people with and without dementia (59).

### *In vitro* studies in model of the human BBB (hCMEC/D3 cells)

We confirmed the expression of GLUT1 in the hCMEC/D3 cells with WBs. These results are in line with previous studies. GLUT1 protein expression has been detected in hCMEC/D3 cells and in brain microvascular endothelial cells derived from healthy patient-derived induced pluripotent stem cells (61,62). Other glucose transporters’ mRNA has also been detected in the hCMEC/D3 cells, namely GLUT3 and sodium-dependent glucose transporter 1 (SGLT1) (62,63). Protein expression of GLUT3 and GLUT4, but not SGLT1, has also been reported in hCMEC/D3 cells (61). Nevertheless, there is consensus that GLUT1 is the most highly expressed transporter in the hCMEC/D3 cells and the BBB (64).

*In vitro*, our self-inhibition study using 4 mM non-labelled glucose, showed that 4 mM glucose significantly decreased the uptake of [^14^C]D-glucose into the hCMEC/D3 cells but did not affect membrane integrity as measured with [^3^H]mannitol. This suggests the presence of functional glucose transporter on the hCMEC/D3 cells.

Our *in vitro* experiments looking at the interaction between antipsychotics and GLUT1 showed no effect of micromolar concentrations of amisulpride on the accumulation of [^14^C]D-glucose into the hCMEC/D3 cells. This suggests that clinically relevant concentrations of amisulpride do not inhibit GLUT1 (amisulpride has a plasma Cmax of 0.17-1.19 μM) (4,65). However, these results are difficult to interpret conclusively as the GLUT1 is partially saturated due to the presence of non-labelled glucose in the control accumulation assay buffer. The non-labelled glucose being essential for endothelial cell survival.

Overall, the high affinity (Km = ∼2mM (66)) of the GLUT1 transporter, high expression of GLUT1 at the BBB, excess glucose concentrations in buffer/plasma and the low concentrations of amisulpride could make the interaction between amisulpride and GLUT1 difficult to detect *in vitro.* Amisulpride has been associated with hyperglycemia as a side effect in 1 in 100 people (67). The hyperglycaemia possibly being caused by inhibition of GLUT1 transport. Interestingly, recent studies did not associate its use with a higher prevalence of diabetes (68)

Cell culture studies have shown that another second-generation antipsychotic - clozapine, inhibits glucose uptake (at 20 µM) in PC12 rat cells after a 30-minute incubation. Exposure to clozapine beyond 24 h (at concentrations up to 20 µM), there was a significant increase in the cellular expression of GLUT1 and GLUT3 (23).

Later studies have confirmed that both clozapine and risperidone interact with glucose transporters and inhibit glucose transport. It has been suggested that this could be mediated by directly binding to glucose transporters and allosterically modulating either a glucose binding side or the conformational change in the protein conformation required for transport (24). Importantly, risperidone which is the only antipsychotics licensed for use in AD, shows treatment response and emergent side effects at very low doses and low plasma concentrations (7), similar to amisulpride (10). Also, clozapine is used at low doses and at correspondingly low plasma concentrations in the treatment of psychosis in Parkinson’s disease patients (69).

### Animal model studies in WT and 5xFamilial AD (FAD) mice

In order to study the effect of AD on the BBB and how this contributes to the increased sensitivity of AD patients to antipsychotics, we need good preclinical models. In this study we utilized the 5xFAD mice. The significantly lower weight of the female mice, compared to the male mice of the 5xFAD genotype which we observed, matches with previous reports that female mice develop the pathology earlier than male mice (70). This could be related to increased expression of the Thy1 promoter which drives the transgenes in the 5xFAD mice and has an oestrogen response element (71) resulting in generation of higher levels of Aβ (72).

#### TEM ultrastructural study

Using TEM, we detected Aβ plaques in the frontal cortex of 5xFAD mice from 6 months of age. This is in agreement with previous data, reporting amyloid deposition in these mice from the age of 2 months (40) and with our Western blot studies which confirmed APP expression.

#### *In situ* brain perfusions

When we studied the permeability of [^3^H]amisulpride across the BBB in a whole animal *in vivo*, we observed low permeability of the drug across the BBB in WT and in 5xFAD mice (both groups 12-15 months of age).

Interestingly, as measured by [^14^C]sucrose (MW 359.48 g/mol), there was no significant change in vascular integrity between age-matched WT and 5xFAD mice in all areas that we investigated even though there was a trend for increased permeability in the 5xFAD mice in the frontal cortex, thalamus, supernatant, and homogenate. Importantly, there was significantly higher uptake of [^3^H]amisulpride (corrected for [^14^C]sucrose) in the supernatant of 5xFAD mice compared to WT mice. An earlier animal study has also reported difference in the brain uptake of amisulpride in an AD mouse model compared to WT, without significant change in the [^14^C]sucrose uptake, i.e. without significant change in BBB integrity (16). The 3xTg mouse model of AD had increased [^3^H]amisulpride uptake in the frontal cortex, but not the occipital cortex, compared to WT mice at 24 months of age (age matched) (16). Thus, suggesting that the increased uptake of amisulpride in the brain of the 5xFAD mice could be related to regional BBB changes in transporter expression associated with AD.

#### Western blot studies

A decrease in GLUT1 expression in brain samples has been observed previously using WB in 5xFAD mice compared to WT at 9 months of age (73). This study used whole brain samples and pooled the samples from all of their WT mice together and from all of their 5xFAD mice together (73). In our study we confirmed the expression of GLUT1 in 5xFAD and WT mouse capillaries at 12-15 months of age. We did not observe significant difference between the genotypes, this is likely related to the fact that we did not pool our samples but analysed each mouse separately which would have increased variability.

Other studies using the 5xFAD mouse model have reported a decrease in the expression of P-gp and GLUT1 in the cortex capillaries of 6-month-old 5xFAD mice, compared to age-matched WT mice (74). We did not detect any effect of the genotype; this could be because we used the whole mouse brain to prepare the capillary samples and to perform WB, and the expression changes could be region-specific.

Finally, expression of PMAT, MATE1, OCT1, and P-gp has been reported in the BBB of 3xTg mice. No difference in expression levels was observed when they were compared to age matched WT mice (16). These results are in line with our observations in the 5xFAD model. This could be because both studies used the whole brain to do capillary isolation, whereas the AD-associated transporter expression changes might be regional, as seen in human control and AD cases (16).

### Human control and AD tissue studies

#### TEM ultrastructural study

In our AD case we were able to identify brain capillaries, myelinated axons and neurites using TEM. Our images showed signs of NVU, and brain degeneration such as swollen basement membrane and oedema around the capillaries, lipofuscin granules, degenerating neurites, degenerating myelin, and fibrillary depositions.

Previous EM studies have also reported abnormalities in the brain capillaries, associated with AD, these include, splitting and duplication of the basement membrane, reduction of the length of the tight junctions, morphological alterations of the mitochondria of the endothelial cells, the pericytes and the perivascular astrocytic processes. The number of the pinocytic vesicles was substantially increased in the endothelium of the brain capillaries in AD in comparison with age-matched controls. Thus, it has been suggested that abnormalities in the brain capillaries may result in the release of neurotoxic factors and abnormal Aβ homeostasis in the brain and contribute to AD pathology (75).

#### Western blot studies

Expression of the endothelial marker TfR1 in our human brain capillary lysates confirms that we have isolated brain capillary pellets enriched in endothelial cells. There was no significant change in TfR1 expression between control and AD samples observed.

Diminished glucose uptake has been reported in the hippocampus, parietotemporal cortex and/or posterior cingulate cortex in individuals at genetic risk for AD (76), positive family history (77) and/or mild or no cognitive impairment who develop AD (78,79). Reduced levels of GLUT1 in cerebral microvessels have also been reported in AD in the caudate nucleus (80), frontal cortex (protein was decreased but not mRNA) (19), and the hippocampus (18). It has been suggested that GLUT1 deficiency can contribute to the disease process, acting in tandem with Aβ to initiate or amplify vascular damage and Aβ accumulation (81). We did not observe a significant change in GLUT1 expression in the frontal cortex and caudate between controls and AD patients. This could be related to the insufficient selectivity of the antibody, the need of more cases, or the heterogeneity in the group of cases.

P-gp is ubiquitously and abundantly expressed in the brain capillaries (82). Importantly, protein expression has been reported to decrease significantly in the prefrontal cortex of AD patients, compared to healthy ageing controls (61 to 100 years old) (83). In line with other studies, we observed a decrease in the P-gp expression in the caudate capillaries of AD patients, compared to healthy controls. Importantly, P-gp protects the brain from potentially toxic substances and has been reported to extrude Aβ from the brain (84). It has been suggested that downregulation of P-gp could allow pharmaceuticals into the central nervous system and may increase the accumulation of Aβ (83).

## Conclusion

In conclusion, a literature review identified GLUT1 substrates, GLUT1 inhibitors and antipsychotics that interacted with GLUT. Physicochemical characterization of these groups using chemical property databases established that amisulpride had similar properties to the GLUT-interacting antipsychotics group. Our *in silico* molecular docking studies revealed that amisulpride interacts with GLUT1 and could potentially affect glucose delivery to the CNS. We also have *in vitro* evidence for the presence of functional glucose transporter in the hCMEC/D3 cells line. However, we could not detect any interaction of amisulpride with GLUT1 in this assay. This is possibly because glucose being essential for endothelial cell survival limits the sensitivity of this assay for exploring antipsychotic interaction with GLUT1 *in vitro*. TEM and WB analysis validated the 5xFAD mouse model for our study. The *in situ* brain perfusion studies showed limited entry of amisulpride across the BBB in both WT and 5xFAD mice, and an increased uptake into the brain of the 5xFAD mice compared to WT mice, although the cerebrovascular space was similar in both genotypes. Our WB work with P-gp further confirms that transporter expression at the human BBB is altered in AD, although a significant difference was not observed for GLUT1 expression in our cases. It is possible that amisulpride competitively inhibits glucose entry via GLUT1, which may further compromise the neurovascular unit and increase BBB permeability, and therefore increase central drug access, and contribute to the amisulpride sensitivity observed in AD. This research further confirms the national guidance that antipsychotic drugs should only be prescribed at the lowest dose possible for the shortest durations in AD. It is also plausible that, in the longer term, the impact on energy delivery to the brain may lead to further cellular degeneration. The implications of our findings extend to other antipsychotic drugs.

## List of abbreviations

3xTgAD: triple transgenic,
Aβ: amyloid beta,
AD: model;
5xFAD: five times familial Alzheimer’s disease mouse model;
ABC: Adenosine triphosphate binding cassette;
AD: Alzheimer’s disease;
APP: amyloid precursor protein;
ASN: asparagine;
BBB: blood–brain barrier;
BCA: bicinchoninic acid;
BDR: Brain for Dementia Research;
dpm: disintegrations per minute;
CAS: chemical abstracts service number;
FBS: foetal bovine serum;
GLN: glutamine;
GLU: glutamate;
GLUT1: glucose transporter 1;
GLY: glycine;
hCMEC/D3: immortalized human cerebral microvessel endothelial cell line;
HEK cells: human embryonic kidney cells;
MATEs: multi-drug and toxin extrusion proteins;
MW: molecular weight;
OCT: organic cation transporters;
NVU: neurovascular unit;
PFA: paraformaldehyde;
P-gp: P-glycoprotein;
PHE: phenylalanine;
PMAT: plasma membrane monoamine transporter;
PMD: post-mortem delay;
RIPA: radio-immunoprecipitation assay;
SGLT1: sodium-dependent glucose transporter 1;
SLC: solute carrier;
TEM: transmission electron microscopy;
TfR1: transferrin receptor 1;
THR: threonine;
TRP: tryptophan;
V_d_: volume of distribution;
WB: Western blot;
WT: wild type.

## Declarations

### Ethics approval and consent to practice

All *in vivo* animal experiments were performed in accordance with the Animal Scientific Procedures Act (1986) and Amendment Regulations 2012 and with consideration to the Animal Research: Reporting of *In Vivo* Experiments (ARRIVE) guidelines. The study was approved by the King’s College London Animal Welfare and Ethical Review Body. The UK government home office project license number was 70/7755.

BDR has ethical approval granted by the National Health Service (NHS) health research authority (NRES Committee London-City & East, UK: REC reference:08/H0704/128+5. IRAS project ID:120436). Tissue samples were supplied by The Manchester Brain Bank and the London Neurodegenerative Diseases Brain Bank, which are both part of the BDR programme, jointly funded by Alzheimer’s Research UK and Alzheimer’s Society. Tissue was received on the basis that it will be handled, stored, used, and disposed of within the terms of the Human Tissue Act 2004.

## Consent for publication

This research was funded in whole, or in part, by the Wellcome Trust [080268]. For the purpose of Open Access, the author has applied a CC BY public copyright licence to any Author Accepted Manuscript version arising from this submission.

## Availability of data and materials

The datasets supporting the conclusions of this article are included within the article and supplementary S1 File.

## Acknowledgements

We acknowledge the support of Professor D.M. Mann from the Manchester Brain Bank who provided some of the human brain tissue samples used in this study. We would also like to thank Ms S. Selvackadunco from the London Neurodegenerative Diseases Brain Bank for her assistance with acquiring the human brain samples used in our WB and TEM studies. We also acknowledge the support of Professors I. Romero, B. Weksler, and P. Couraud who provided the hCMEC/D3 cell line under MTA.

## Supporting Information

- S1 File.docx
- 6 Supplementary Tables
- 13 Supplementary Figures

## References

1 Jeste D V, Jin H, Golshan S, Mudaliar S, Glorioso D, Fellows I, et al. Discontinuation of quetiapine from an NIMH-funded trial due to serious adverse events. Am J Psychiatry. 2009 Aug;166(8):937–8.

2. Jeste D V, Blazer D, Casey D, Meeks T, Salzman C, Schneider L, et al. ACNP White Paper: update on use of antipsychotic drugs in elderly persons with dementia. Neuropsychopharmacology [Internet]. 2008 Apr;33(5):957–70. Available from: http://dx.doi.org/10.1038/sj.npp.1301492

3. Murray PS, Kumar S, Demichele-Sweet MAA, Sweet RA. Psychosis in Alzheimer’s disease. Biol Psychiatry [Internet]. 2014 Apr;75(7):542–52. Available from: http://dx.doi.org/10.1016/j.biopsych.2013.08.020

4. Reeves S, Bertrand J, D’Antonio F, McLachlan E, Nair A, Brownings S, et al. A population approach to characterise amisulpride pharmacokinetics in older people and Alzheimer’s disease. Psychopharmacology (Berl) [Internet]. 2016 Sep;233(18):3371–81. Available from: http://dx.doi.org/10.1007/s00213-016-4379-6

5. Ballard C, Howard R. Neuroleptic drugs in dementia: benefits and harm. Nat Rev Neurosci [Internet]. 2006 Jun;7(6):492–500. Available from: http://dx.doi.org/10.1038/nrn1926

6. Schneider LS, Tariot PN, Dagerman KS, Davis SM, Hsiao JK, Ismail MS, et al. Effectiveness of atypical antipsychotic drugs in patients with Alzheimer’s disease. N Engl J Med [Internet]. 2006 Oct;355(15):1525–38. Available from: http://dx.doi.org/10.1056/NEJMoa061240

7. Reeves S, Bertrand J, Uchida H, Yoshida K, Otani Y, Ozer M, et al. Towards safer risperidone prescribing in Alzheimer’s disease. British Journal of Psychiatry. 2021 May 1;218(5):268–75.

8. Schlösser R, Gründer G, Anghelescu I, Hillert A, Ewald-Gründer S, Hiemke C, et al. Long-Term Effects of the Substituted Benzamide Derivative Amisulpride on Baseline and Stimulated Prolactin Levels. Neuropsychobiology. 2002;46(1):33–40.

9. Reeves S, Eggleston K, Cort E, McLachlan E, Brownings S, Nair A, et al. Therapeutic D2/3 receptor occupancies and response with low amisulpride blood concentrations in very late-onset schizophrenia-like psychosis (VLOSLP). Int J Geriatr Psychiatry. 2018;33(2):396–404.

10. Reeves S, Bertrand J, D’Antonio F, McLachlan E, Nair A, Brownings S, et al. A population approach to characterise amisulpride pharmacokinetics in older people and Alzheimer’s disease. Psychopharmacology (Berl) [Internet]. 2016 Sep;233(18):3371–81. Available from: http://dx.doi.org/10.1007/s00213-016-4379-6

11. Reeves S, McLachlan E, Bertrand J, D’Antonio F, Brownings S, Nair A, et al. Therapeutic window of dopamine D2/3 receptor occupancy to treat psychosis in Alzheimer’s disease. Brain. 2017;140(4):1117–27.

12. Clark-Papasavas C, Dunn JT, Greaves S, Mogg A, Gomes R, Brownings S, et al. Towards a therapeutic window of D2/3 occupancy for treatment of psychosis in Alzheimer’s disease, with [18F]fallypride positron emission tomography. Int J Geriatr Psychiatry [Internet]. 2014 Oct;29(10):1001–9. Available from: http://dx.doi.org/10.1002/gps.4090

13. Hiemke C, Baumann P, Bergemann N, Conca A, Dietmaier O, Egberts K, et al. AGNP Consensus Guidelines for Therapeutic Drug Monitoring in Psychiatry: Update 2011. Pharmacopsychiatry. 2011 Sep;44(6):195–235.

14. Lako IM, van den Heuvel ER, Knegtering H, Bruggeman R, Taxis K. Estimating dopamine DLJ receptor occupancy for doses of 8 antipsychotics: a meta-analysis. J Clin Psychopharmacol. 2013 Oct;33(5):675–81.

15. Sparshatt A, Taylor D, Patel MX, Kapur S. Amisulpride - dose, plasma concentration, occupancy and response: implications for therapeutic drug monitoring. Acta Psychiatr Scand. 2009 Dec;120(6):416–28.

16. Sekhar GN, Fleckney AL, Boyanova ST, Rupawala H, Lo R, Wang H, et al. Region-specific blood–brain barrier transporter changes leads to increased sensitivity to amisulpride in Alzheimer’s disease. Fluids Barriers CNS. 2019 Dec 17;16(1):38.

17. dos Santos Pereira JN, Tadjerpisheh S, Abu Abed M, Saadatmand AR, Weksler B, Romero IA, et al. The poorly membrane permeable antipsychotic drugs amisulpride and sulpiride are substrates of the organic cation transporters from the SLC22 family. AAPS J. 2014 Nov;16(6):1247–58.

18. Horwood N, Davies DC. Immunolabelling of hippocampal microvessel glucose transporter protein is reduced in Alzheimer’s disease. Virchows Arch. 1994;425(1):69–72.

19. Mooradian AD, Chung HC, Shah GN. GLUT-1 expression in the cerebra of patients with Alzheimer’s disease. Neurobiol Aging. 18(5):469–74.

20. Kalaria RN, Harik SI. Reduced glucose transporter at the blood-brain barrier and in cerebral cortex in Alzheimer disease. J Neurochem. 1989 Oct;53(4):1083–8.

21. Hooijmans CR, Graven C, Dederen PJ, Tanila H, van Groen T, Kiliaan AJ. Amyloid beta deposition is related to decreased glucose transporter-1 levels and hippocampal atrophy in brains of aged APP/PS1 mice. Brain Res. 2007 Nov 21;1181:93–103.

22. Merlini M, Meyer EP, Ulmann-Schuler A, Nitsch RM. Vascular β-amyloid and early astrocyte alterations impair cerebrovascular function and cerebral metabolism in transgenic arcAβ mice. Acta Neuropathol. 2011 Sep;122(3):293–311.

23. Dwyer DS, Pinkofsky HB, Liu Y, Bradley RJ. Antipsychotic drugs affect glucose uptake and the expression of glucose transporters in PC12 cells. Prog Neuropsychopharmacol Biol Psychiatry. 1999 Jan;23(1):69–80.

24. Ardizzone TD, Bradley RJ, Freeman AM, Dwyer DS. Inhibition of glucose transport in PC12 cells by the atypical antipsychotic drugs risperidone and clozapine, and structural analogs of clozapine. Brain Res [Internet]. 2001 Dec 27;923(1–2):82–90. Available from: http://www.ncbi.nlm.nih.gov/pubmed/11743975

25. Lean MEJ, Pajonk FG. Patients on Atypical Antipsychotic Drugs. Diabetes Care. 2003 May 1;26(5):1597–605.

26. Dwyer DS, Donohoe D. Induction of hyperglycemia in mice with atypical antipsychotic drugs that inhibit glucose uptake. Pharmacol Biochem Behav. 2003 May;75(2):255–60.

27. Dwyer DS, Pinkofsky HB, Liu Y, Bradley RJ. Antipsychotic drugs affect glucose uptake and the expression of glucose transporters in PC12 cells. Prog Neuropsychopharmacol Biol Psychiatry. 1999 Jan;23(1):69–80.

28. Boyanova S, Wang H, Fleckney AL, Gatt A, Farag DB, Rahman KM, et al. Heightened sensitivity of people with Alzheimer’s disease to the side effects of antipsychotic drug amisulpride may be mediated through an interaction with glucose transporter 1 at the bloodLJbrain barrier. In: Alzheimer’s Dement 2020;16(Suppl3):e047395. 2020.

29. Bethesda (MD): National Library of Medicine (US) NC for BI. PubMed. https://pubmed.ncbi.nlm.nih.gov/ Accessed 16 March 2021.

30. Wishart DS, Feunang YD, Guo AC, Lo EJ, Marcu A, Grant JR, et al. DrugBank 5.0: a major update to the DrugBank database for 2018. Nucleic Acids Res. 2018;46(D1):D1074–82.

31. MarvinSketch version 22.9.0. ChemAxon http://chemaxon.com Accessed April 2022 and April 2023.

32. Oliva L, Fernandez-Lopez JA, Remesar X, Alemany M. The Anomeric Nature of Glucose and Its Implications on Its Analyses and the Influence of Diet: Are Routine Glycaemia Measurements Reliable Enough? J Endocrinol Metab. 2019;9(3):63–70.

33. Almahmoud S, Wang X, Vennerstrom JL, Zhong HA. Conformational Studies of Glucose Transporter 1 (GLUT1) as an Anticancer Drug Target. Molecules. 2019 Jun 7;24(11).

34. Weksler BB, Subileau EA, Perrière N, Charneau P, Holloway K, Leveque M, et al. Blood-brain barrier-specific properties of a human adult brain endothelial cell line. FASEB J. 2005 Nov;19(13):1872–4.

35. Schneider CA, Rasband WS, Eliceiri KW. NIH Image to ImageJ: 25 years of image analysis. Vol. 9, Nature Methods. 2012. p. 671–5.

36. Wolff A, Antfolk M, Brodin B, Tenje M. In Vitro Blood-Brain Barrier Models-An Overview of Established Models and New Microfluidic Approaches. J Pharm Sci. 2015 Sep;104(9):2727–46.

37. Cloyd JC, Snyder BD, Cleeremans B, Bundlie SR, Blomquist CH, Lakatua DJ. Mannitol pharmacokinetics and serum osmolality in dogs and humans. J Pharmacol Exp Ther. 1986 Feb;236(2):301–6.

38. Sprague JE, Arbeláez AM. Glucose counterregulatory responses to hypoglycemia. Pediatr Endocrinol Rev. 2011 Sep;9(1):463–73; quiz 474–5.

39. Richard BC, Kurdakova A, Baches S, Bayer TA, Weggen S, Wirths O. Gene Dosage Dependent Aggravation of the Neurological Phenotype in the 5XFAD Mouse Model of Alzheimer’s Disease. J Alzheimers Dis. 2015;45(4):1223–36.

40. Oakley H, Cole SL, Logan S, Maus E, Shao P, Craft J, et al. Intraneuronal beta-amyloid aggregates, neurodegeneration, and neuron loss in transgenic mice with five familial Alzheimer’s disease mutations: potential factors in amyloid plaque formation. J Neurosci. 2006 Oct 4;26(40):10129–40.

41. Rae EA, Brown RE. The problem of genotype and sex differences in life expectancy in transgenic AD mice. Neurosci Biobehav Rev. 2015 Oct;57:238–51.

42. Szu J, Jullienne A, Obenaus A, Territo PR, Consortium M. Lifespan neuroimaging of the 5xFAD mouse model of Alzheimer’s disease: Evolution of metabolic and vascular perturbations. Alzheimer’s & Dementia. 2020 Dec 7;16(S4).

43. Jastrow H. Dr. Jastrows electron microscopic atlas. http://www.drjastrow.de/WAI/EM/EMAtlas.html Accessed 10 October 2019.

44. Mason BL, Pariante CM, Jamel S, Thomas SA. Central Nervous System (CNS) Delivery of Glucocorticoids Is Fine-Tuned by Saturable Transporters at the Blood-CNS Barriers and Nonbarrier Regions. Endocrinology. 2010 Nov 1;151(11):5294–305.

45. Báez-Mendoza R, Schultz W. The role of the striatum in social behavior. Front Neurosci. 2013;7.

46. Garbuzova-Davis S, Rodrigues MCO, Hernandez-Ontiveros DG, Louis MK, Willing AE, Borlongan C v, et al. Amyotrophic lateral sclerosis: a neurovascular disease. Brain Res. 2011 Jun 29;1398:113–25.

47. Merlo S, Nakayama ABS, Brusco J, Rossi MA, Carlotti CG, Moreira JE. Lipofuscin Granules in the Epileptic Human Temporal Neocortex with Age. Ultrastruct Pathol. 2015;39(6):378–84.

48. Pickett EK, Rose J, McCrory C, McKenzie CA, King D, Smith C, et al. Region-specific depletion of synaptic mitochondria in the brains of patients with Alzheimer’s disease. Acta Neuropathol. 2018;136(5):747–57.

49. Sanderson L, Dogruel M, Rodgers J, de Koning HP, Thomas SA. Pentamidine movement across the murine blood-brain and blood-cerebrospinal fluid barriers: effect of trypanosome infection, combination therapy, P-glycoprotein, and multidrug resistance-associated protein. J Pharmacol Exp Ther. 2009 Jun;329(3):967–77.

50. Ouellette RJ, Rawn JD. Carbohydrates. In: Organic Chemistry. Elsevier; 2018. p. 889–928.

51. Dwyer DS, Liu Y, Bradley RJ. Dopamine receptor antagonists modulate glucose uptake in rat pheochromocytoma (PC12) cells. Neurosci Lett. 1999 Oct;274(3):151–4.

52. Chow VW, Mattson MP, Wong PC, Gleichmann M. An Overview of APP Processing Enzymes and Products. Neuromolecular Med. 2010 Mar 24;12(1):1–12.

53. Babkin P, George Thompson AM, Iancu C v, Walters DE, Choe JY. Antipsychotics inhibit glucose transport: Determination of olanzapine binding site in Staphylococcus epidermidis glucose/H(+) symporter. FEBS Open Bio. 2015;5:335–40.

54. Chokhawala K, Stevens L. Antipsychotic Medications [Internet]. In: StatPearls [Internet]. Treasure Island (FL): StatPearls Publishing; 2023 [cited 2023 Apr 6]. Available from: https://www.ncbi.nlm.nih.gov/books/NBK519503/

55. Natesan S, Reckless GE, Barlow KBL, Nobrega JN, Kapur S. Amisulpride the ‘atypical’ atypical antipsychotic — Comparison to haloperidol, risperidone and clozapine. Schizophr Res. 2008 Oct;105(1–3):224–35.

56. Devés R, Krupka RM. Cytochalasin B and the kinetics of inhibition of biological transport. A case of asymmetric binding to the glucose carrier. Biochimica et Biophysica Acta (BBA) - Biomembranes. 1978 Jul;510(2):339–48.

57. Kapoor K, Finer-Moore JS, Pedersen BP, Caboni L, Waight A, Hillig RC, et al. Mechanism of inhibition of human glucose transporter GLUT1 is conserved between cytochalasin B and phenylalanine amides. Proc Natl Acad Sci U S A. 2016 Apr 26;113(17):4711–6.

58. Niccoli T, Cabecinha M, Tillmann A, Kerr F, Wong CT, Cardenes D, et al. Increased Glucose Transport into Neurons Rescues Aβ Toxicity in Drosophila. Current Biology. 2016 Sep;26(17):2291–300.

59. Kable A, Fullerton A, Fraser S, Palazzi K, Hullick C, Oldmeadow C, et al. Comparison of Potentially Inappropriate Medications for People with Dementia at Admission and Discharge during An Unplanned Admission to Hospital: Results from the SMS Dementia Study. Healthcare. 2019 Jan 9;7(1):8.

60. Orsel K, Taipale H, Tolppanen AM, Koponen M, Tanskanen A, Tiihonen J, et al. Psychotropic drugs use and psychotropic polypharmacy among persons with Alzheimer’s disease. European Neuropsychopharmacology. 2018 Nov;28(11):1260–9.

61. Al-Ahmad AJ. Comparative study of expression and activity of glucose transporters between stem cell-derived brain microvascular endothelial cells and hCMEC/D3 cells. Am J Physiol Cell Physiol. 2017 Oct 1;313(4):C421–9.

62. Ohtsuki S, Ikeda C, Uchida Y, Sakamoto Y, Miller F, Glacial F, et al. Quantitative targeted absolute proteomic analysis of transporters, receptors and junction proteins for validation of human cerebral microvascular endothelial cell line hCMEC/D3 as a human blood-brain barrier model. Mol Pharm. 2013 Jan 7;10(1):289–96.

63. Weksler B, Romero IA, Couraud PO. The hCMEC/D3 cell line as a model of the human blood brain barrier. Fluids Barriers CNS. 2013 Mar 26;10(1):16.

64. Meireles M, Martel F, Araújo J, Santos-Buelga C, Gonzalez-Manzano S, Dueñas M, et al. Characterization and modulation of glucose uptake in a human blood-brain barrier model. J Membr Biol. 2013 Sep;246(9):669–77.

65. Coukell AJ, Spencer CM, Benfield P, Carpenter Tr WT. DRUG EVALUATION Amisulpride eNS A Review of its Pharmacodynamic and Pharmacokinetic Properties and Therapeutic Efficacy in the Management of Schizophrenia. Vol. 6, Drugs. 1996.

66. Burant CF, Sivitz WI, Fukumoto H, Toshiaki K, Nagamatsu S, Susumo S, et al. Mammalian Glucose Transporters: Structure and Molecular Regulation. Vol. 47, RECENT PROGRESS IN HORMONE RESEARCH: Proceedings of the 1990 Laurentian Hormone Conference. 1991.

67. Zentiva. Patient Information Leaftlet: Amisulpride. https://www.medicines.org.uk/emc/files/pil.3965.pdf Accessed 13 January 2022.

68. Holt RIG. Association Between Antipsychotic Medication Use and Diabetes. Curr Diab Rep. 2019 Oct 2;19(10):96.

69. Ruggieri S, de Pandis MF, Bonamartini A, Vacca L, Stocchi F. Low dose of clozapine in the treatment of dopaminergic psychosis in Parkinson’s disease. Clin Neuropharmacol. 1997 Jun;20(3):204–9.

70. Forner S, Kawauchi S, Balderrama-Gutierrez G, Kramár EA, Matheos DP, Phan J, et al. Systematic phenotyping and characterization of the 5xFAD mouse model of Alzheimer’s disease. Sci Data. 2021;8(1):270.

71. Sadleir KR, Eimer WA, Cole SL, Vassar R. Aβ reduction in BACE1 heterozygous null 5XFAD mice is associated with transgenic APP level. Mol Neurodegener. 2015 Jan 7;10:1.

72. Maarouf CL, Kokjohn TA, Whiteside CM, Macias MP, Kalback WM, Sabbagh MN, et al. Molecular Differences and Similarities Between Alzheimer’s Disease and the 5XFAD Transgenic Mouse Model of Amyloidosis. Biochem Insights. 2013;6:1–10.

73. Ahn KC, Learman CR, Dunbar GL, Maiti P, Jang WC, Cha HC, et al. Characterization of Impaired Cerebrovascular Structure in APP/PS1 Mouse Brains. Neuroscience. 2018;385:246–54.

74. Park R, Kook SY, Park JC, Mook-Jung I. Aβ1-42 reduces P-glycoprotein in the blood-brain barrier through RAGE-NF-κB signaling. Cell Death Dis. 2014 Jun 26;5:e1299.

75. Baloyannis SJ. Brain capillaries in Alzheimer’s disease. Hell J Nucl Med. 18 Suppl 1:152.

76. Ossenkoppele R, Prins ND, van Berckel BN. Amyloid imaging in clinical trials. Alzheimers Res Ther. 2013;5(4):36.

77. Mosconi L, Rinne JO, Tsui WH, Murray J, Li Y, Glodzik L, et al. Amyloid and metabolic positron emission tomography imaging of cognitively normal adults with Alzheimer’s parents. Neurobiol Aging. 2013 Jan;34(1):22–34.

78. Mosconi L, Mistur R, Switalski R, Tsui WH, Glodzik L, Li Y, et al. FDG-PET changes in brain glucose metabolism from normal cognition to pathologically verified Alzheimer’s disease. Eur J Nucl Med Mol Imaging. 2009 May;36(5):811–22.

79. Landau SM, Harvey D, Madison CM, Koeppe RA, Reiman EM, Foster NL, et al. Associations between cognitive, functional, and FDG-PET measures of decline in AD and MCI. Neurobiol Aging. 2011 Jul;32(7):1207–18.

80. Simpson IA, Chundu KR, Davies-Hill T, Honer WG, Davies P. Decreased concentrations of GLUT1 and GLUT3 glucose transporters in the brains of patients with Alzheimer’s disease. Ann Neurol. 1994 May;35(5):546–51.

81. Winkler EA, Nishida Y, Sagare AP, Rege S v, Bell RD, Perlmutter D, et al. GLUT1 reductions exacerbate Alzheimer’s disease vasculo-neuronal dysfunction and degeneration. Nat Neurosci. 2015 Apr;18(4):521–30.

82. Löscher W, Langer O. Imaging of P-glycoprotein function and expression to elucidate mechanisms of pharmacoresistance in epilepsy. Curr Top Med Chem. 2010;10(17):1785–91.

83. Chiu C, Miller MC, Monahan R, Osgood DP, Stopa EG, Silverberg GD. P-glycoprotein expression and amyloid accumulation in human aging and Alzheimer’s disease: preliminary observations. Neurobiol Aging. 2015 Sep;36(9):2475–82.

84. Chai AB, Leung GKF, Callaghan R, Gelissen IC. P-glycoprotein: a role in the export of amyloid-β in Alzheimer’s disease? FEBS J. 2020;287(4):612–25.

