## Supplementary material for "Interaction of amisulpride with GLUT1 at the blood-brain barrier. Relevance to Alzheimer’s disease": S1 File

**Supporting Information**

**S1 File**

**Supplementary Table 1:** Primary and secondary antibodies used for Western blotting with the proteins they detect and their predicted molecular weight (MW). They were all made up in TBS-T with 5% milk. Actual band sizes may differ due to post translational modifications or cleavages. Validation data is available from the Abcam website (1), Cell Signalling Technology website (2) and Alomone labs website (3). Validation data is available from the BIOSS USA antibodies website (4). Verification data is available from ThermoFisherScientific (5).

| Protein | Primary antibody | Secondary antibody (WB) | MW |
| --- | --- | --- | --- |
| APP/Aβ | Purified anti-β-Amyloid, 1-16 Antibody (Biolegend; #803004; RRID: AB_2715854), Dilution: 1:1000 | Rabbit Anti-Mouse IgG H&L (HRP) Abcam; #ab6728; RRID: AB_955440),  Dilution: 1:3000 | 100 kDa/4 kDa |
| TfR1 | Mouse Transferrin Receptor Monoclonal Antibody (H68.4), (Thermo Fisher; #13-6800; RRID: AB_2533029),  Dilution: 1:1000 | Rabbit Anti-Mouse IgG H&L (HRP) (Abcam, #ab6728; RRID: AB_955440),  Dilution: 1:3000 | 190 kDa |
| PMAT  (SLC29A4) | Rabbit polyclonal reactive to human, mouse and rat (Bioss Antibodies; #bs-4176R; RRID: AB_11108960),  Dilutions:  1:600 for mouse capillaries | Goat anti-rabbit (IgG)-HRP (Cell Signalling; #7074S; RRID: [AB_2099233](http://antibodyregistry.org/AB_2099233)) Dilution:  1:1000 | 58 kDa |
| MATE1  (SLC47A1) | Rabbit polyclonal reactive to mouse and rat (Alonome; # ANT-131; RRID: AB_2751020)  Dilution:  1:1000 for mouse capillaries | Goat anti-rabbit (IgG)-HRP (Cell Signalling; #7074S; RRID: [AB_2099233](http://antibodyregistry.org/AB_2099233)) Dilution:  1:1000 | 75 kDa |
| OCT1  (SLC22A1) | Rabbit polyclonal reactive to human, predicted to work with mouse, rat, rabbit, etc. (Abcam, #ab55916; RRID: AB_882579) Dilution: 1:667 | Goat anti-rabbit (IgG)-HRP (Cell Signalling; #7074S; RRID: [AB_2099233](http://antibodyregistry.org/AB_2099233)) Dilution:  1:1000 | 61 kDa |
| GLUT1 (SLC2A1) | Recombinant Anti-Glucose Transporter GLUT1 antibody [EPR3915] (Abcam, #ab115730; RRID:AB_10903230) Dilution: 1:100 000 | Goat anti-rabbit (IgG)-HRP (Cell Signalling; #7074S; RRID: [AB_2099233](http://antibodyregistry.org/AB_2099233)) Dilution:  1:2000 | 40-60 kDa |
| MCT1 (SLC16A1) | Mouse polyclonal Monocarboxylic acid transporter 1 antibody (Abcam; #ab90582  RRID: AB_2050317)  Dilution: 1:1000 | Rabbit Anti-Mouse IgG H&L (HRP) (Abcam, #ab6728; RRID: AB_955440),  Dilution: 1:2000 | 40-50 kDa  54 kDa (predicted) |
| P-gp  (MDR1, ABCB1, CD243) | Rabbit monoclonal [EPR10364-57] reactive to mouse, rat, human (Abcam; #ab170904; RRID: AB_2687930),  Dilution:  1:1000 | Goat anti-rabbit (IgG)-HRP (Cell Signalling; #7074S; RRID: [AB_2099233](http://antibodyregistry.org/AB_2099233)) Dilution:  1:1000 (for mouse capillaries); 1:2000 (for human capillaries) | 140 kDa |
| GADPH | Rabbit polyclonal reactive to mouse, rat, chicken, dog, human, etc. (Abcam, #ab9485; RRID: AB_307275)  Dilution:  1:2500 – 1:1000 | Goat anti-rabbit (IgG)-HRP (Cell Signalling; #7074S; RRID: [AB_2099233](http://antibodyregistry.org/AB_2099233)) Dilution:  1:1000-1:2000 | 37kDa |

Supplementary Table 2: Case number, Braak stage, age, sex, and post-mortem delay (PMD) of the control and AD cases used in the WB study.

Human brain tissue samples were supplied by The London and Manchester Brain Banks, which are part of the Brains for Dementia Research programme, jointly funded by Alzheimer’s Research UK and Alzheimer’s Society. The cases were divided into two groups based on Braak staging. The mean±SEM age of cases with a Braak stage 0-II was 87.0±2.5 years (healthy controls), it was not significantly different to the age of cases with a Braak-stage of V-VI, which was 81.0±2.1 years (AD affected individuals) (unpaired two-tailed t-test, t=1.812, df=21, p=0.0844). The mean post-mortem delay was 50.4±6.9 h (healthy controls) versus 69.2±9.3 h (AD affected individuals). These values are not significantly different to each other (unpaired two-tailed t-test, t=1.449, df=21, p=0.1620). The mean brain weight in the healthy controls group was 1253.0±55.3 g, and the mean brain weight in the AD group was 1095±42.7 g, these values were not significantly different (t=2.007, df=13, p=0.0660). The mean post-mortem pH was 6.5±0.1 (healthy controls) and 6.1±0.1 (AD affected individuals). These values are significantly different to each other (unpaired two-tailed t-test, t=2.959, df=18, p=0.0084). Three brain regions (frontal cortex, caudate nucleus, and putamen) were obtained from each of the cases. The majority of information tabulated was obtained directly from the brain banks. The information in brackets was obtained from the tissue database (6). When two values were present for any characteristic of the samples, the value in brackets was used to calculate mean and SEM.

| Case number | Sex | Age at death (years) | PMD (hours) | Clinical Diagnosis | Pathological diagnosis | Braak or  (BNE) stage | Alleles of APOE gene: e2, e3 or e4 | Brain weigh (g) | Average brain pH | TRID |
| --- | --- | --- | --- | --- | --- | --- | --- | --- | --- | --- |
| BBN002_30176 | Female | 96 | 72 | Control | control case- minimal Alzheimer’s disease pathology (BNE/ Braak) stage 1 with amyloid angiopathy | (I) | 3,3 |  | 6.1 | 265 |
| BBN002_30171 | Female | 86 | 27 | Control | control case-age related changes and very mild Alzheimer's disease pathology (modified Braak/BNE stage 2) | (II) | 3,3 |  |  | 265 |
| BBN002_30069 | Female | 98 | 76 | Control | control (AD modified Braak (BNE) stage 2 with moderate amyloid angiopathy | (II) | 3,3 |  | 6.34 | 265 |
| BBN_24666 | Female | 74 | 66 | Control | Braak stage 2 consistent with ageing, control | II |  |  | 6.59 | 265 |
| BBN_24297 | Female | 83 | 39 | Control | Tau braak stage 2 in keeping with ageing, control | II | 3,4 | 1166 | 6.52 | 265 |
| BBN002_26151 | Male | 82 | 26 | Control | Control (very mild Alzheimer disease pathology- modified Braak BNE stage 2) | (II) | 3,4 | 1409 | 6.64 | 265 |
| BBN_23398 | Male | 80(83) | 31 | Control | Tau Braak Stage 2 consistent with ageing process, control | II | 3,3 | 1184 | 5.95 | 265 |
| BBN_14408 | Male | 90 | 45(70) | Control | Mild age-related changes (control brain) - AD modified Braak stage + with mild focal amyloid angiopathy | (0-I) | 2,3 | 1252 | 6.9 | 265 |
| BBN_11072 | Male | 91 | 47 | Control | Consistent with ageing, Braak stage 2 | II | 3,3 |  | 6.98 | 265 |
| Mean±SEM |  | **87.0±2.5** | **50.4±6.9** |  |  |  |  | **1253.0±55.3** | **6.5±0.1** |  |
| BBN_24530 | Female | 91 | 131 | AD | Mild non-amyloid SVD Braak tangle stage VI CERAD high density of neuritic plaques Braak LB stage 0. Alzheimer's disease. Mild SVD. | VI | 3,4 | 1000 | 6.02 | 170 |
| BBN_19609 | Male | 75(74) | 62 | AD | Braak tangle stage VI CERAD high density of neuritic plaques Braak LB stage 0  Thal phase not assessed.  Alzheimer's disease. | VI | 3,3 | 970 | 6.05 | 170 |
| BBN_24943 | Male | 71 | 96 | AD | Mild non-amyloid SVD Braak tangle stage VI CERAD high density of neuritic plaques Braak LB stage 0 Moderate/severe arteriosclerosis in occipital white matter Moderate/severe occipital leptomeningeal CAA Thal phase not assessed. Alzheimer's disease. Mild SVD | VI | 3,4 | 1163 | 6.21 | 170 |
| BBN_25109 | Female | 79 | 97 | AD | Moderate non-amyloid SVD Braak tangle stage VI CERAD high density of neuritic plaques Braak LB stage 0 Thal phase 5 Moderate arteriolar CAA in occipital leptomeninges Moderate/severe arteriosclerosis in occipital white matter Moderate/severe occipital leptomeningeal CAA. Alzheimer's disease. Moderate to severe SVD | VI | 4,4 | 948 | 6.54 | 170 |
| BBN_25921 | Male | 81 | 68 | AD | Mild non-amyloid SVD  Mild arteriolar Aß-CAA  Braak tangle stage V  CERAD high density of neuritic plaques  Braak LB stage 0. Alzheimer's disease. Mild SVD. | V-VI | 3,4 | 1318 | 6.03 | 170 |
| BBN_11027 | Female | 85 | 73 | AD | moderate to severe SVD, v. Mild DLB | V-VI | 3,4 | 1030 |  | 265 |
| BBN_3472 | Male | 73 | 107 | AD | mod SVD | VI | 4,4 | 1180 |  | 265 |
| BBN005_29239 | Male | 83 | 49 | AD | Transitional DLB | VI | 3,4 | 1330 | 6.41 | 265 |
| BBN005_26134 | Female | 95 | 108 | AD | CAA, Secondary TDP43, Marked SVD | VI | 3,4 | 938 | 5.99 | 265 |
| BBN002_29842 | Male | 81 | 30 | AD | Alzheimer's disease BNE stage 6  mild cerebral amyloid angiopathy (capillary type) | VI | 3,3 |  | 5.94 | 265 |
| BBN002_29634 | Male | 72 | 41 | AD | Alzheimer's disease (modified Braak/BNE stage 6) with extensive and capillary amyloid angiopathy and significant cerebrovascular pathology | (VI) | 3,3 |  | 5.91 | 265 |
| BBN002_30072 | Female | 82 | 65.5 | AD | Alzheimer's disease BNE stage 6, Moderate small vessel disease.  Amygdala only Lewy body pathology | (VI) |  |  | 5.63 | 265 |
| BBN002_32856 (WB and TEM) | Female | 74 | 19 | AD  Pneumonia | Alzheimer's disease, BNE stage VI with moderate amyloid angiopathy type 1, moderate cerebrovascular disease VCING mod. | (VI) |  | 1145 | 6.36 | 287 |
| BBN002_35217 | Female | 93 | 22 | AD | Cerebrovascular disease (probable vascular Parkinsonism) and Alzheimer disease (BNE stage IV) with extensive amyloid angiopathy and limbic TDP43 pathology and unusual p62 positive neuronal intranuclear inclusions | (IV) |  | 1018 | 6.13 | 287 |
| Mean±SEM |  | **81.0±2.1** | **69.2±9.3** |  |  |  |  | **1095.0±42.7** | **6.1±0.1** |  |

Supplementary Table 3: Molecules identified as substrates of GLUT1 by literature review are tabulated. We present their chemical abstracts service (CAS) number, structure, MW (g/mol), predicted gross charge at physiological pH, the physiological charge of the top two microspecies of each molecule at physiological pH, and total number of microspecies at physiological pH. The specific references which identified each molecule as a GLUT1 substrate are also tabulated. The MW and structure of the molecules was obtained from the Drug Bank website (7). The charge, and microspecies information was obtained via MarvinSketch version 22.9.0 (8).

| **GLUT1 Substrate (with reference)**  **IUPAC**  **CAS** | **Structure** | **MW (g/mol)**  **(DrugBank)** | **Gross charge distribution at pH 7.4 (MarvinSketch)** | **Physiological charge, % distribution of**  **the top two microspecies, and total number of microspecies at pH 7.4 (MarvinSketch)** |
| --- | --- | --- | --- | --- |
| 3-O-methyl-d-glucose (9)  CAS: 146-72-5 | 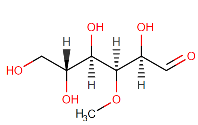 | 194.18 | 0 | 0 (100%)  Total number of microspecies at pH 7.4: n=1 |
| D-Galactose (10)  Or (3R,4S,5R,6R)-6-(hydroxymethyl)oxane-2,3,4,5-tetrol  CAS: 10257-28-0 | 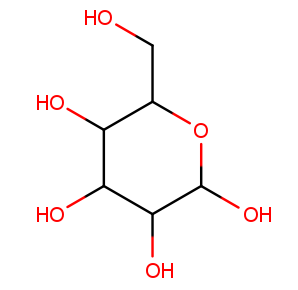 | 180.16 | 0 | 0 (99.99%)  -1 (0.01%)  Total number of microspecies at pH 7.4: n=2 |
| D-Glucosamine  (10)  CAS: 3416-24-8 | 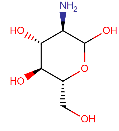 | 179.17 | +0.827 | +1 (82.74%)  0 (17.18%)  Total number of microspecies at pH 7.4: n=3 |
| Alpha-D-glucose (11)  CAS: 492-62-6 | 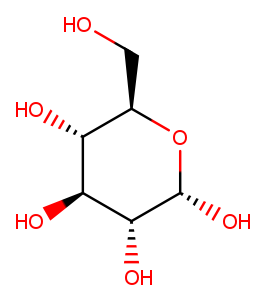 | 180.16 | -0.00 | 0 (99.99%)  -1 (0.01%)  Total number of microspecies at pH 7.4: n=2 |
| Beta-D-glucose (11)  CAS: 492-61-5 | 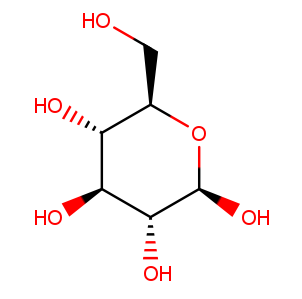 | 180.16 | -0.00 | 0 (99.99%)  -1 (0.01%)  Total number of microspecies at pH 7.4: n=2 |
| L-dehydroascorbic acid (DHA) (12)  CAS: 490-83-5 | 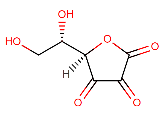 | 174.11 | -0.030 | 0 (57.47%)  -1 (42.53%)  Total number of microspecies at pH 7.4: n= 2 |
| D-Mannose  (10)  CAS: 530-26-7 | 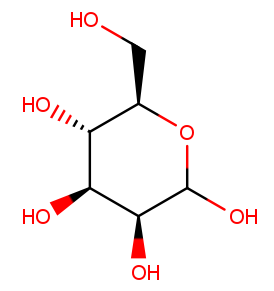 | 180.16 | -0.00 | 0 (99.99%)  -1 (0.01%)  Total number of microspecies at pH 7.4: n=2 |
| Melatonin (13)  CAS: 73-31-4 | 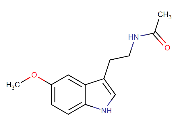 | 232.28 | -0.00 | 0 (100%)  Total number of microspecies at pH 7.4: n=1 |
| 2-Deoxy-d-glucose  CAS: 154-17-6 | 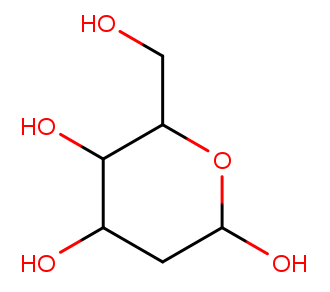 | 164.16 | 0 | 0 (100%)  Total number of microspecies at pH 7.4: n=1 |

Supplementary Table 4: Molecules identified as inhibitors of GLUT1 by literature review are tabulated. We present their CAS number, structure, MW (g/mol), predicted gross charge at physiological pH, the physiological charge of the top two microspecies of each molecule, and total number of microspecies at physiological pH. The specific references which identified each molecule as a GLUT1 inhibitor are also tabulated. The MW and structure (except where stated) of the molecules was obtained from the Drug Bank website (7). The charge, and microspecies information was obtained via MarvinSketch version 22.9.0 (8).

| **GLUT1 Inhibitor (with reference)**  **IUPAC**  **CAS** | **Structure** | **MW (g/mol)**  **(DrugBank)** | **Gross charge distribution at pH 7.4 (MarvinSketch)** | **Physiological charge, % distribution of**  **the top two microspecies, and total number of microspecies at pH 7.4 (MarvinSketch)** |
| --- | --- | --- | --- | --- |
| Bay-876 (14)  CAS: 1799753-84-6 | 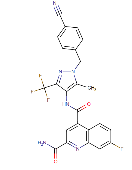 | 496.43 | -0.000 | 0 (100%)  Total number of microspecies at pH 7.4: n=1 |
| Biochanin A (15)  CAS: 491-80-5 | 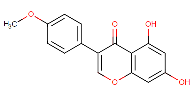 | 284.27 | -0.271 | 0 (73.55%)  -1 (22.58%)  Total number of microspecies at pH 7.4: n=4 |
| Caffeine (15)  CAS: 58-08-2 | 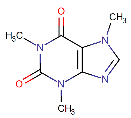 | 194.19 | 0 | 0 (100%)  Total number of microspecies at pH 7.4: n=1 |
| Curcumin (15)  CAS: 458-37-7 | 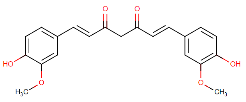 | 368.38 | -0.036 | 0 (96.06%)  -1 (0.76%)  Total number of microspecies at pH 7.4: n=5 |
| Cytochalasin B (13)  CAS: 14930-96-2 | 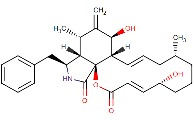 | 479.62 | -0.000 | 0 (100%)  Total number of microspecies at pH 7.4: n=1 |
| Epigallocatechin-3-gallate  (EGCG) (16)  CAS: 989-51-5 | 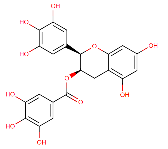 | 458.37 | -0.279 | 0 (78.88%)  -1 (13.86%)  Total number of microspecies at pH 7.4: n=19 |
| Fasentin (17)  CAS: 392721-37-8 | 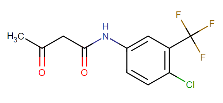 | 279.64 | -0.001 | 0 (99.95%)  -1 (0.05%)  Total number of microspecies at pH 7.4: n=2 |
| Forskolin (9)  CAS: 66575-29-9 | 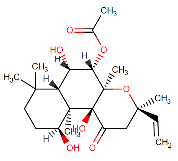 | 410.50 | -0.000 | 0 (99.99%)  -1 (0.01%)  Total number of microspecies at pH 7.4: n=2 |
| Genistein (15)  CAS: 446-72-0 | 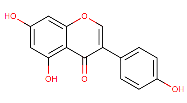 | 270.24 | -0.320 | 0 (71.57%)  -1 (3.54%)  Total number of microspecies at pH 7.4: n=8 |
| Gossypol (15)  CAS: 20300-26-9 | 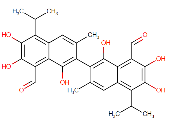 | 518.55 | -0.316 | 0 (79.18%)  -1 (12.17%)  Total number of microspecies at pH 7.4: n=13 |
| Mercuric chloride (HgCl_2_) (9)  Or dichloromercury  CAS: 7487-94-7 | 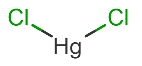 | 271.50 | 0.000 | No ionisable atoms found |
| Isorhamnetin  or 3’-methoxquercetin (15)  CAS: 480-19-3 | 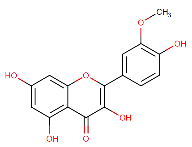 | 316.26 | -0.745 | 0 (40.55%)  -1 (30.27%)  Total number of microspecies at pH 7.4: n=15 |
| Kaempferol (15)  CAS: 520-18-3 | 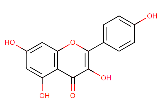 | 286.24 | -0.740 | 0 (41.73%)  -1 (26.15%)  Total number of microspecies at pH 7.4: n=15 |
| Lavendustin A (15)  CAS: 125697-92-9 | 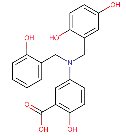 | 381.38 | -1.032 | -1 (97.67%)  -2 (1.45%)  Total number of microspecies at pH 7.4: n= 6 |
| Lavendustin B (15)  CAS: 125697-91-8 | 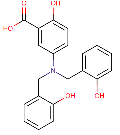 | 365.38 | -1.043 | -1 (97.08%)  -2 (2.88%)  Total number of microspecies at pH 7.4: n=4 |
| Methyl-2,5-dihydroxycinnamate (15)  CAS: 63177-57-1 | 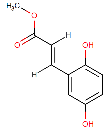 | 194.19 | -0.008 | 0 (99.18%)  -1 (0.5%)  Total number of microspecies at pH 7.4: n=3 |
| Morin (15)  CAS: 480-16-0 | 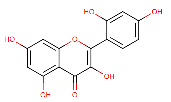 | 302.24 | -0.893 | -1 (37.02%)  0 (28.03%)  Total number of microspecies at pH 7.4: n=25 |
| Myricetin (15)  CAS: 529-44-2 | 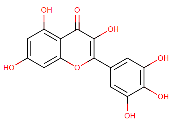 | 318.23 | -0.881 | -1 (36.93%)  0 (29.85%)  Total number of microspecies at pH 7.4: n=23 |
| Nordihydroguaiaretic acid (NDGA)(15)  CAS: 500-38-9 | 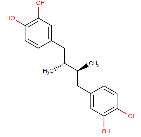 | 302.36 | -0.023 | 0 (98.46%)  -1 (0.49%)  -1 (0.49%)  Total number of microspecies at pH 7.4: n=5 |
| Pentoxifylline (15)  CAS: 6493-05-6 | 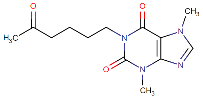 | 278.31 | 0 | 0 (100%)  Total number of microspecies at pH 7.4: n=1 |
| Phloretin (9,15)  CAS: 60-82-2 | 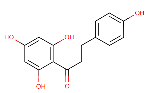 | 274.27 | -0.210 | 0 (73.18%)  -1 (18.81%)  Total number of microspecies at pH 7.4: n=7 |
| Quercetin (16)  CAS: 117-39-5 | 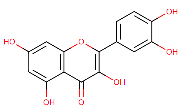 | 302.24 | -0.800 | 0 (36.69%)  -1 (32.90%)  Total number of microspecies at pH 7.4: n=19 |
| Resveratrol (15)  CAS: 501-36-0 | 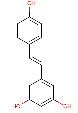 | 228.24 | -0.107 | 0 (92.41%)  -1 (5.11%)  Total number of microspecies at pH 7.4: n=5 |
| Rhamnetin or 7-Methoxyquercetin (15)  CAS: 90-19-7 | 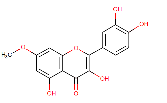 | 316.26 | -0.524 | 0 (51.50%)  -1 (38.59%)  Total number of microspecies at pH 7.4: n=12 |
| STF-31 (13,16)  Or 4-[[(4-tert-butylphenyl)sulfonylamino]methyl]-N-pyridin-3-ylbenzamide  CAS: 724741-75-7 | 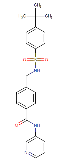 | 423.53 | -0.000 | 0 (99.81%)  -1 (0.10%)  Total number of microspecies at pH 7.4: n=3 |
| Theophylline (15)  CAS: 58-55-9 | 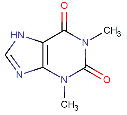 | 180.16 | -0.084 | 0 (72.23%)  -1 (27.77%)  Total number of microspecies at pH 7.4: n=2 |
| Tyrphostin B44 (15)  CAS: 133550-32-0 | 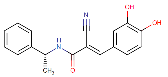 | 308.34 | -0.085 | 0 (91.47%)  -1 (6.51%)  Total number of microspecies at pH 7.4: n=3 |
| Tyrphostin B46 (15)  CAS: 133550-34-2 | 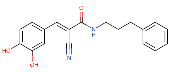 | 322.36 | -0.085 | 0 (91.47%)  -1 (6.51%)  Total number of microspecies at pH 7.4: n=3 |
| Tyrphostin B48 (15)  CAS: 133550-35-3 | 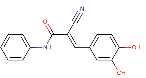 | 280.28 | -0.086 | 0 (91.45%)  -1 (6.53%)  Total number of microspecies at pH 7.4: n=3 |
| Tyrphostin B50 (15)  CAS: 133550-37-5 | 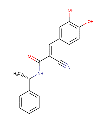(MarvinSketch 22.9) | 308.34 | -0.085 | 0 (91.47%)  -1 (6.51%)  Total number of microspecies at pH 7.4: n=3 |
| Tyrphostin B56 (15)  CAS: 133550-41-1 | 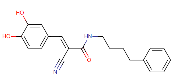 | 336.39 | -0.085 | 0 (91.47%)  -1 (6.51%)  Total number of microspecies at pH 7.4: n=3 |
| Tyrphostin AG879 (15)  CAS: 148741-30-4 | 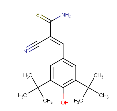 (MarvinSketch 22.9) | 316.46 | -0.963 | -1 (96.33%)  0 (3.67%)  Total number of microspecies at pH 7.4: n=2 |
| WZB117 (13,17)  Or  3-Fluoro-1,2-phenylene bis(3-hydroxybenzoate)  CAS: 1223397-11-2 | 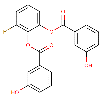 | 368.32 | -0.055 | 0 (96.31%)  -1 (1.83%)  -1 (1.83%)  Total number of microspecies at pH 7.4: n=4 |

Supplementary Table 5: The antipsychotics suggested to interact with GLUT in our literature review are tabulated. We present their CAS number, structure, MW (g/mol), predicted gross charge at physiological pH, the physiological charge of the top two microspecies of each molecule, and total number of microspecies at physiological pH. The specific references are also tabulated. The MW and structure of the molecules was obtained from the Drug Bank website (7). The charge, and microspecies information was obtained via MarvinSketch version 22.9.0 (8).

| **Type of antipsychotic** | **Antipsychotics affecting GLUT1 substrates transport through GLUT**  **IUPAC**  **CAS** | **Structure** | **MW (g/mol)**  **(DrugBank)** | **Gross charge distribution at pH 7.4 (MarvinSketch)** | **Physiological charge, % distribution of**  **the top two microspecies, and total number of microspecies at pH 7.4 (MarvinSketch)** |
| --- | --- | --- | --- | --- | --- |
| **Typical**  **(First generation)** | Chlorpromazine (18–21)  CAS: 50-53-3 | 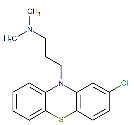 | 318.86 | +0.984 | +1 (98.43%)  0 (1.57%)  Total number of microspecies at pH 7.4: n=2 |
|  | Fluphenazine (18,20–22)  CAS: 69-23-8 | 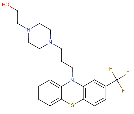 | 437.52 | +0.932 | +1 (78.84%)  +1 (15.39%)  Total number of microspecies at pH 7.4: n=4 |
|  | Haloperidol (18–20,23)  CAS: 52-86-8 | 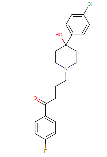 | 375.86 | +0.862 | +1 (86.19%)  0 (13.81%)  Total number of microspecies at pH 7.4: n=2 |
|  | Loxapine (19,20,22)  CAS: 1977-10-2 | 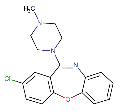 | 327.81 | +0.853 | +1 (85.30%)  0 (14.70%)  Total number of microspecies at pH 7.4: n=2 |
|  | Pimozide (20,21)  CAS: 2062-78-4 | 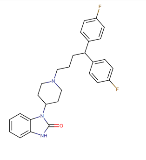 | 461.55 | +0.906 | +1 (90.59%)  0 (9.41%)  Total number of microspecies at pH 7.4: n=2 |
|  | Spiperone (21)  CAS: [749-02-0](https://commonchemistry.cas.org/detail?cas_rn=749-02-0) | 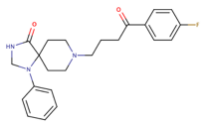 | 395.48 | +0.884 | +1 (88.39%)  0 (11.61%)  Total number of microspecies at pH 7.4: n=2 |
| **Atypical**  **(Second generation)** | Clozapine (18–23)  CAS: 5786-21-0 | 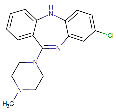 | 326.82 | +0.853 | +1 (85.27%)  0 (14.69%)  Total number of microspecies at pH 7.4: n=3 |
|  | Desmethylclozapine (19,20,22)  CAS: 6104-71-8 | 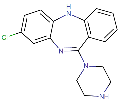 | 312.80 | +0.964 | +1 (96.35%)  0 (3.61%)  Total number of microspecies at pH 7.4: n=3 |
|  | Olanzapine (20,23–25)  CAS: 132539-06-1 | 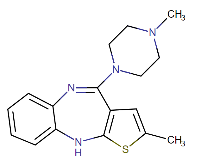 | 312.44 | +0.853 | +1 (85.27%)  0 (14.69%)  Total number of microspecies at pH 7.4: n=3 |
|  | Quetiapine (19,20,23)  CAS: 111974-69-7 | 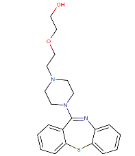 | 383.51 | +0.696 | +1 (69.63%)  0 (30.37%)  Total number of microspecies at pH 7.4: n=2 |
|  | Risperidone (19,20,22,23)  CAS: 106266-06-2 | 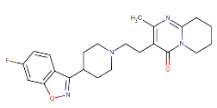 | 410.49 | +0.979 | +1 (97.87%)  0 (2.13%)  Total number of microspecies at pH 7.4: n=2 |

**Supplementary Table 6:** Details characterising the interaction between GLUT, and antipsychotics identified to inhibit GLUT substrates uptake in cells. The table shows IC50 for inhibition of GLUT substrates uptake *in vitro*, the shortest incubation time with the drugs which shows effects on GLUT substrates uptake *in vitro*, and possible mechanism for the observed inhibition. First-generation antipsychotics are dopamine receptor antagonists and are known as typical antipsychotics. Second-generation antipsychotics are serotonin-dopamine antagonists and are also known as atypical antipsychotics.

| **Type of antipsychotic** | **Antipsychotic** | **IC50 (μM)** | **Incubation time for effect on glucose uptake (minutes)** | **Evidence for interaction with GLUT** |
| --- | --- | --- | --- | --- |
| **Typical**  **(First generation)** | Chlorpromazine | 25 (18) | 30(18,20) | * (22) |
|  | Fluphenazine | (18) - 20 (20) | 1-30 (18,20,21) | *  Effect is also seen in L6 myoblast cells (18,22) |
|  | Haloperidol | >200 (19)  >150 (20) | 30 (18,20) | Decrease of glucose uptake in PC12 cells, suggests interaction with GLUT proteins; however, reports show of low potency to no significant effect (18,20) |
|  | Loxapine | 18(22) – 50 (19) (20) | 1-30 (20,22) | Non-competitive inhibitor of glucose transport,  might block glucose transport in a manner similar to cytochalasin B by directly binding to GLUTs and allosterically modulating either a glucose binding site or the conformational changes in the protein  conformation required for transport (22) |
|  | Pimozide | 2-5 (21) | 1-30 (20,21) | Decreases glucose uptake in PC12 cells (20,21) |
|  | Spiperone | No data | 30 (21) | Decreases glucose uptake in PC12 cells (21) |
| **Atypical**  **(Second generation)** | Clozapine | 20 (22) -50 (20) | 1-30 (18,20,22) | *  Effect is also seen in L6 myoblast cells (20,22)  Might block glucose transport in a manner similar to cytochalasin B by directly binding to GLUTs and allosterically modulating either a glucose binding site or the conformational changes in the protein  conformation required for transport (22) |
|  | Desmethylclozapine (metabolite of clozapine) | 15 (22) -20 (20) | 1-30 (20,22) | Inhibits glucose transport in PC12 cells in glucose sensitive manner might block glucose transport in a manner similar to cytochalasin B by directly binding to GLUTs and allosterically modulating either a glucose binding site or the conformational changes in the protein conformation required for transport (22) |
|  | Olanzapine | 80-100 (20)  0.9±0.1 mM (glucose transporter from Staphylococcus epidermidis) (25) | 15 s (25)  30 (20,25)  72 h (24) | * Effect is also seen in L6 myoblast cells (20)  Decreased glucose uptake by olanzapine may be the consequence of conformational alterations at the GLUT1 (study in peripheral blood mononuclear cells) (24)  *In silico* molecular docking data for inhibition of bacterial glucose/H+ symporter from Staphylococcus epidermidis (GlcPSe) (25) |
|  | Quetiapine | 50 (19) - 60 (20) | 30 (20) | Inhibit glucose uptake in dose dependent manner in PC12 cells and in L6 cells, one suggested mechanism is through inhibition of glucose transport (20) |
|  | Risperidone | 35 (22) - 40 (20) | 1-30(20,22) | Non-competitive inhibitor of glucose transport might block glucose transport in a manner similar to cytochalasin B by directly binding to GLUTs and allosterically modulating either a glucose binding site or the conformational changes in the protein conformation required for transport (22) |

***** Rapid decrease of glucose uptake in rat pheochromocytoma (PC12) cells (expressing GLUT1 and GLUT3), suggests direct interaction with GLUT proteins.

**An overview of the experimental design of the study**

**Literature review of substrates and inhibitors of GLUT1, and antipsychotics interacting with GLUT**

- Identification of substrates and inhibitors of GLUT1
- Identification of typical and atypical antipsychotics interacting with GLUT, including GLUT1

**Physicochemical characterisation of substrates and inhibitors of GLUT1, and antipsychotics interacting with GLUT**

- Characterisation of the molecular weight, charge, and microspecies prevalence of the identified substrates, inhibitors, and antipsychotics

***In silico* modelling study**

Characterisation of the molecular interactions between GLUT1, glucose, amisulpride and sucrose.

***In vitro* human cell culture model of BBB (hCMEC/D3)**

**Accumulation assays**

Confirmation of presence of active glucose transporter in hCMEC/D3 cells

Investigation of the interaction between amisulpride and GLUT1 in hCMEC/D3 cells

**Western blot**

Confirmation of GLUT1 expression in hCMEC/D3 cells

***In vivo* mouse model of AD (5xFAD)**

**TEM**

Assessment of Aβ plaque absence or presence in the brain of WT and 5xFAD mice

***In situ* brain perfusions**

Assessment of the BBB integrity in WT and 5xFAD mice

Study of BBB permeability to amisulpride in WT and 5xFAD mice

**Brain tissue and isolated brain capillaries from human control and AD cases**

**TEM**

Examine neurovascular unit structure and brain degeneration in human AD case

**Western blot**

Confirm BBB expression of transporters of interest, including GLUT1 in WT and 5xFAD mice

Compare transporter expression between the genotypes

**Western blot**

Confirm expression of transporters of interest, including GLUT1 at the BBB of control and AD cases

Compare transporter expression between control and AD cases

Supplementary Figure: 1: An overview of the experimental design of the study is shown in this flow chart. Literature review identified the transporter of interest (GLUT1) and the associated substrates and inhibitors. This was followed by physicochemical characterization of the GLUT1 substrates and inhibitors. This information provided the basis for the *in silico*, *in vitro* and *in vivo* experimental design described.

**Effect of unlabelled glucose and amisulpride on hCMEC/D3 cell membrane-integrity-[^3^H]mannitol**

There was no statistically significant effect of non-labelled glucose at a total concentration of 4 mM, 6 mM or 9 mM glucose or 100 μM non-labelled amisulpride on cell membrane integrity as measured by [^3^H]mannitol. However, individual wells in the 9 mM non-labelled glucose and the 100 μM non-labelled amisulpride condition had very high V_d_s, suggesting decreased cell membrane integrity in those wells. As a result, 4 mM non-labelled glucose was selected to test saturable transport of glucose in subsequent accumulation assays.

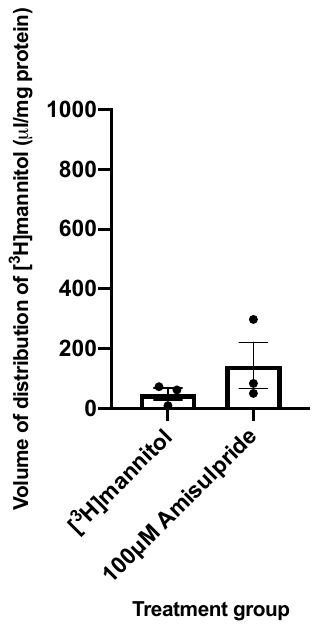

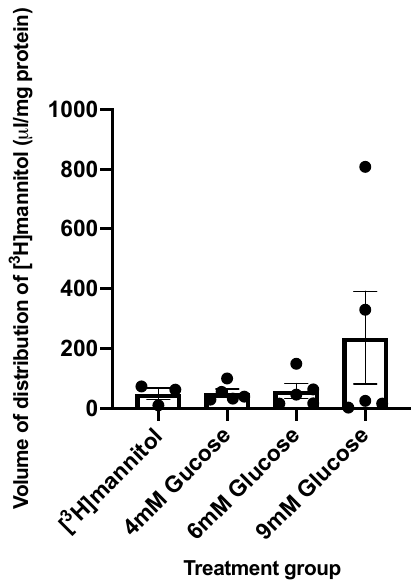

Supplementary Figure 2: Testing the effect of different doses of non-labelled glucose and amisulpride on membrane integrity (V_d_ for [^3^H]mannitol, corrected for protein concentration) in hCMEC/D3 cells. [^3^H]mannitol condition and 100μM amisulpride condition contained 1.1mM non-labelled glucose. Other conditions contained 4, 6 or 9mM non-labelled glucose as indicated. Statistics not performed due to n = 1. Each point (•) represents a technical replicate (well). Error bars represent SEM.

**Weight comparison between WT and 5xFAD mice, and between females and males of WT genotype, as well as females and males of 5xFAD genotype**

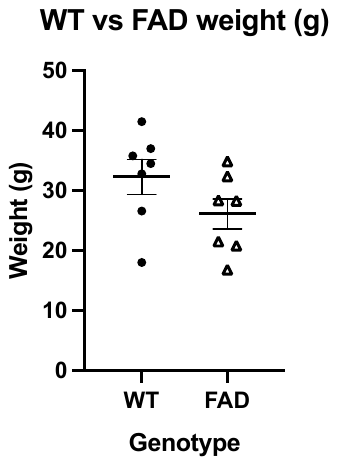

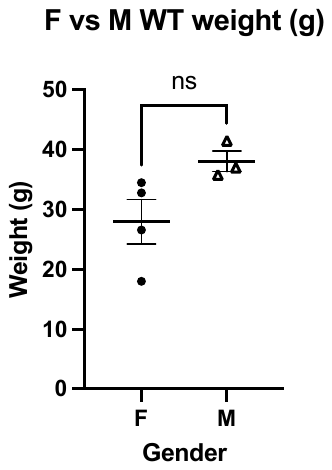

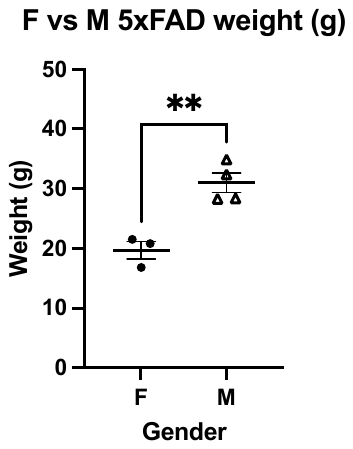

**Supplementary Figure 3:** Weight comparison between WT and 5xFAD mice, and between females and males of WT genotype, as well as females and males of 5xFAD genotype. Data is presented as mean±SEM, analysed using unpaired two-tailed Student's t-test. GraphPad Prism 9.0. Each dot represents a mouse (age 12 to 15 months).

**Representative Western Blots in WT and 5xFAD mice**

**APP WBs**

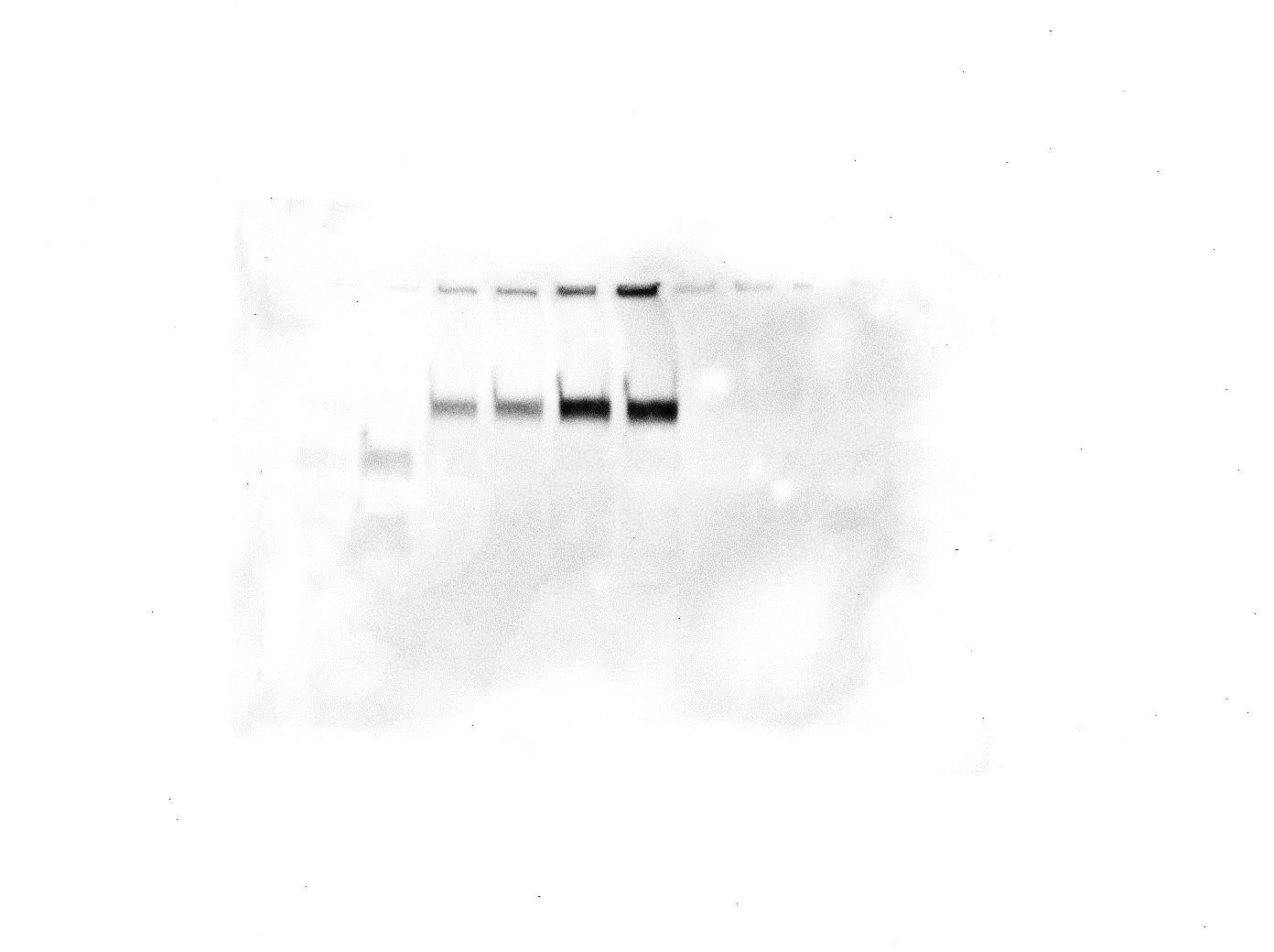

**WT**

**3xTg**

**M8**

**M7**

**M3**

**M2**

**M6**

**M5**

**M4**

**100 kDa - APP**

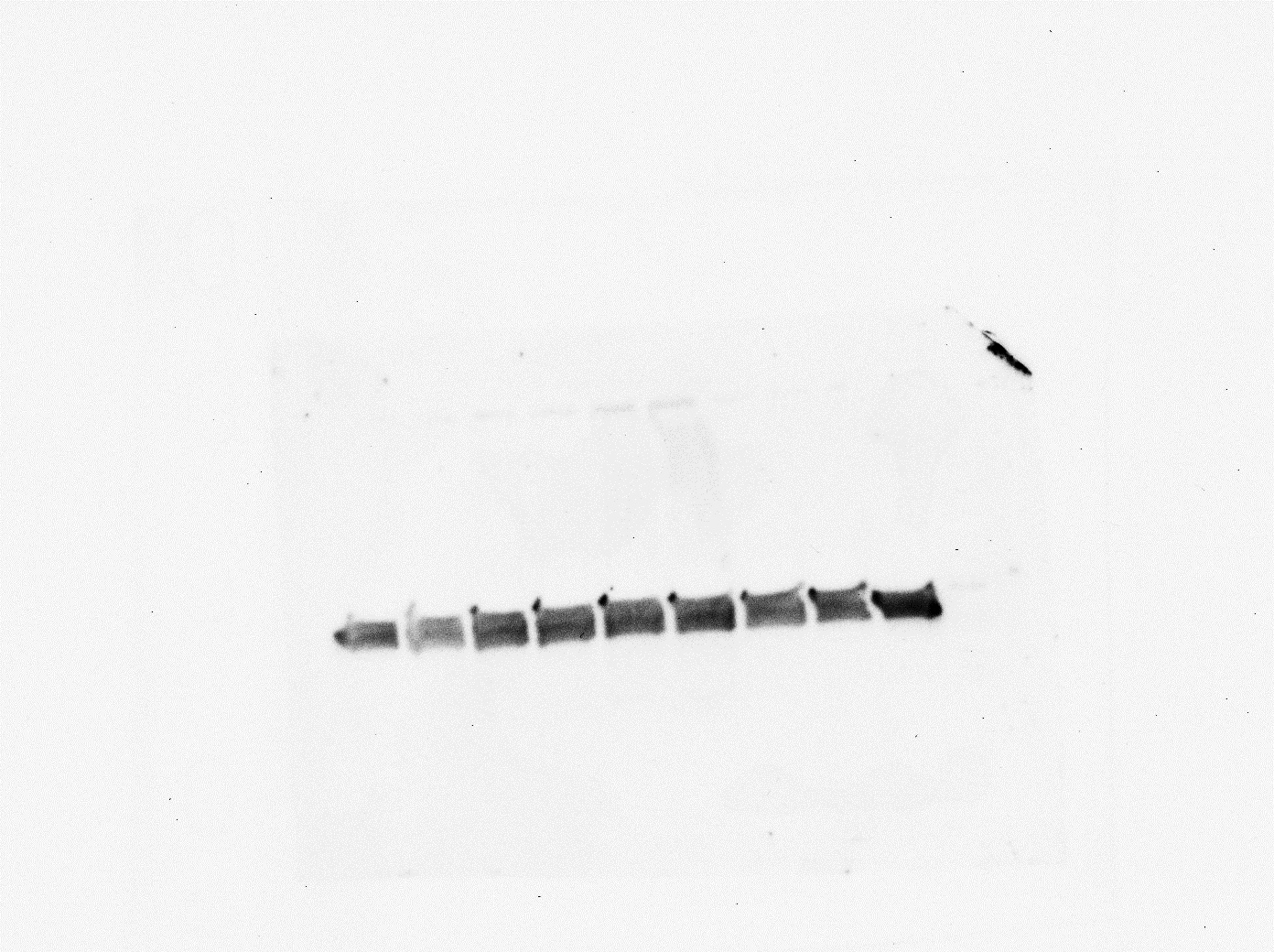

**37 kDa - GAPDH**

**M5**

**M4**

**M6**

**M2**

**WT**

**3xTg**

**M9**

**M3**

**M1**

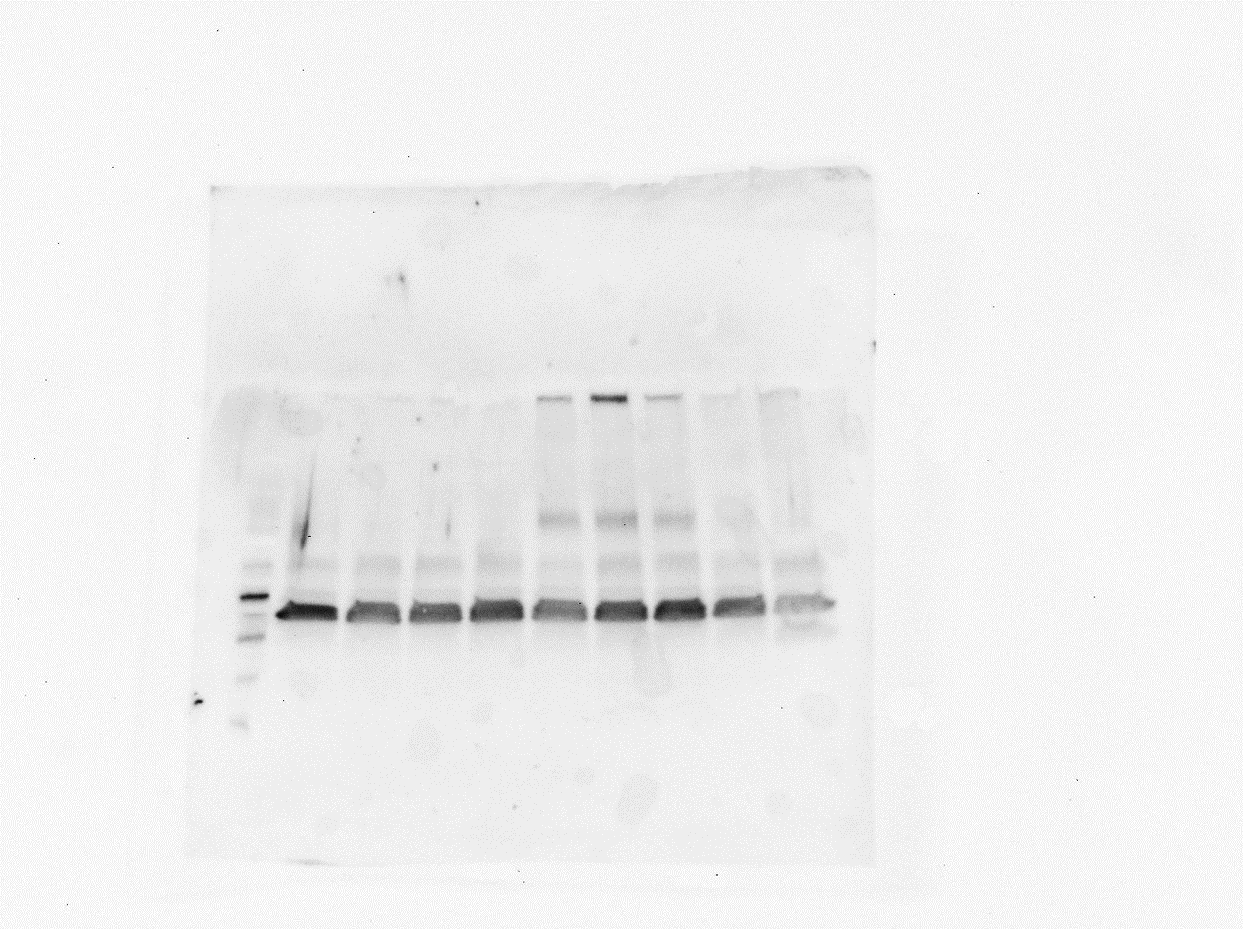

**37 kDa - GAPDH**

**100 kDa - APP**

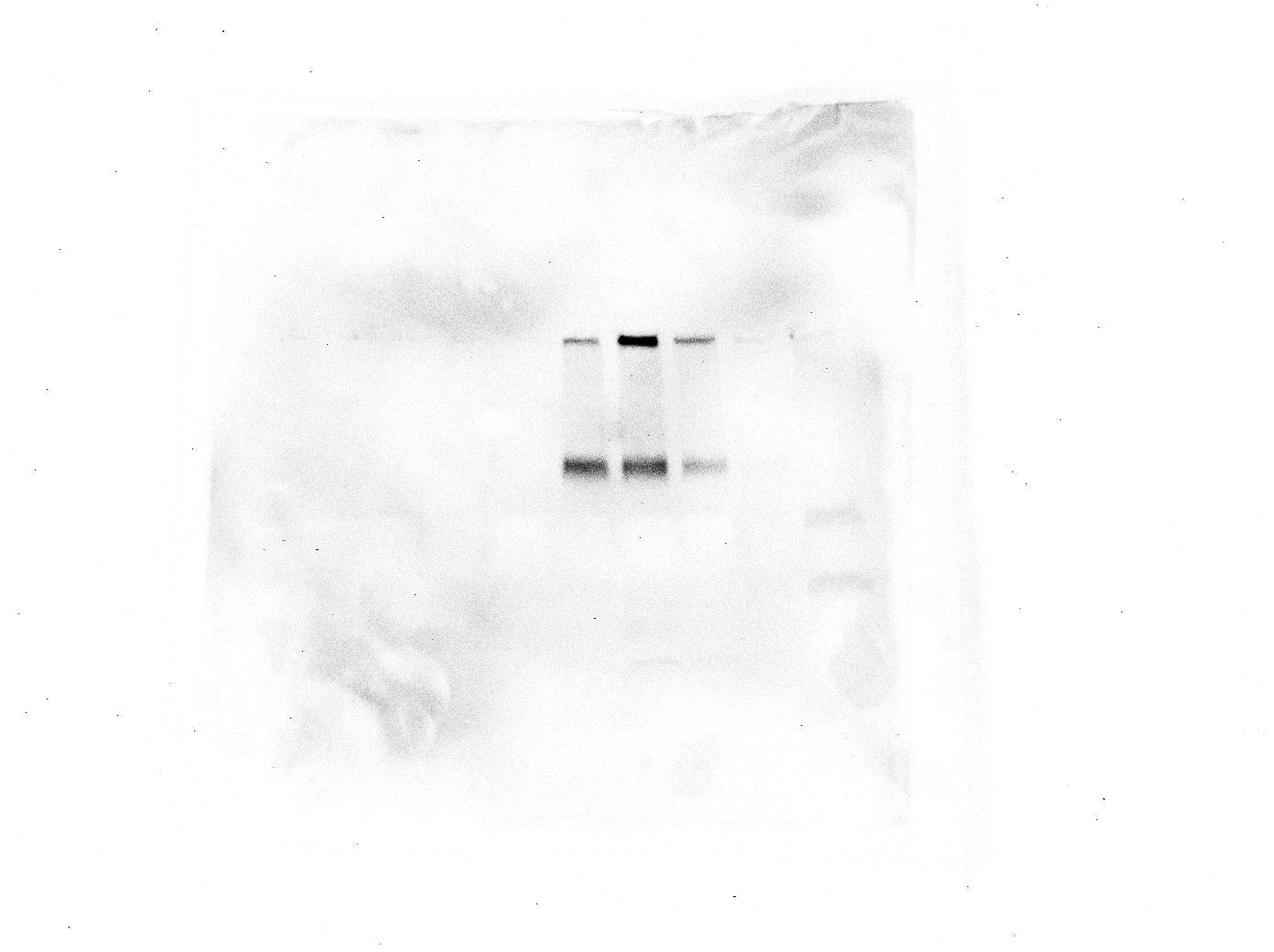

Supplementary Figure 4: Capillary lysates (40 µg per well were loaded) from WT and 5xFAD mice stained with anti-beta Amyloid 1-16 antibody (# 803004, Biolegend, RRID: AB_2715854, detecting APP and Aβ) (n=3 WT and n=4 5xFAD, one membrane M2-8; n=4 WT and n=3 5xFAD, second membrane, M9 and M1-6). GAPDH (37 kDa) was used as a loading control. For the anti-beta amyloid antibody, a secondary anti-mouse antibody was used – 1:3000 (#ab6728, Abcam, UK, RRID:AB_955440). For GAPDH, a secondary anti-rabbit IgG, HRP-linked antibody was used – 1:1000 (#7074, Cell Signalling Technology). Brain capillary lysates from a known WT and 3xTg mice were used as a negative and a positive control, respectively.

**37 kDa - GAPDH**

**58 kDa - PMAT**

**90 kDa – TfR1**

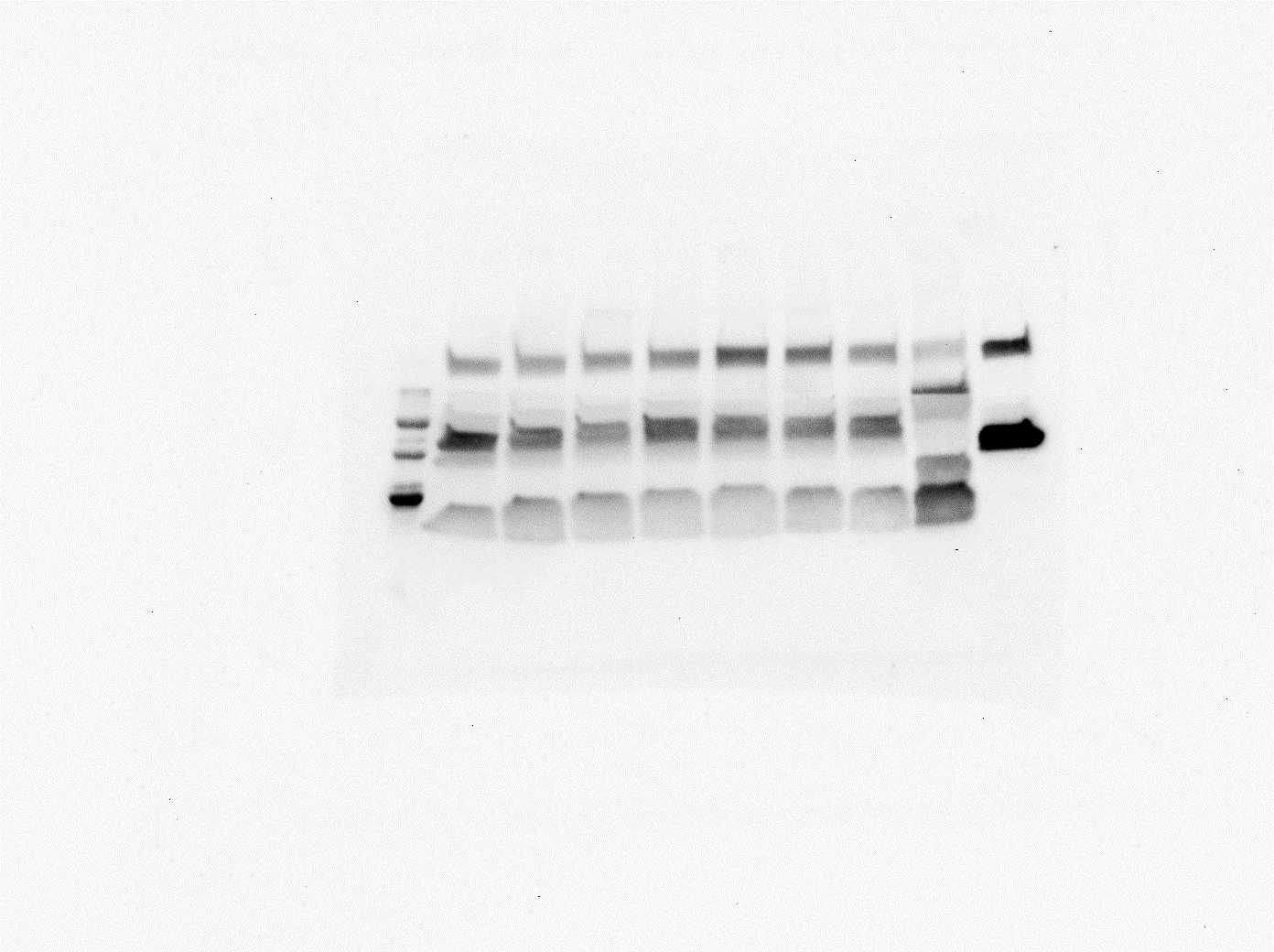

**WT**

**5xFAD**

**Kidney**

**Calu3**

**Supplementary Figure 5:** Brain capillary lysates (40 µg per well were loaded) from C57BL/6 and 5xFAD mice were tested for TfR1 (90 kDa), PMAT (58 kDa) expression, (n=6 WT mice, n=7 5xFAD mice across two membranes, three technical repeats of each, one membrane is presented). GAPDH (37 kDa) was used as a loading control. Antibodies used: anti-TfR1 antibody – 1:1000, #13-6800, Thermo Fisher; anti-PMAT antibody – 1:600, #bs-4176R, Bios; anti-GAPDH antibody – 1:2500, #ab9485, Abcam; secondary anti-rabbit IgG, HRP-linked antibody – 1:1000, #7074, Cell Signalling Technology for PMAT and GAPDH; secondary anti-mouse antibody – 1:3000 (#ab6728, Abcam, UK, RRID:AB_955440) for TfR1.

**Supplementary Figure 6:** Brain capillary lysates (20 µg per well were loaded) from C57BL/6 and 5xFAD mice were tested for GLUT1 (40-60 kDa), (n=5 WT mice, n=4 5xFAD mice across two membranes, three technical repeats of each, except ** samples – which have two repeats, and * – which have one repeat, one membrane is shown). Tubulin (50-60 kDa) was used as a loading control, Caco2 and HEK293 cell lysates were used as a positive and negative control, respectively. Antibodies used: anti-GLUT1 antibody – 1:100 000, #ab115730, Abcam; anti-Tubulin clone DM1A antibody – 1:4500, #05-829, Millipore; secondary anti-rabbit IgG, HRP-linked antibody – 1:2000, #7074, Cell Signalling Technology, secondary anti-mouse antibody IgG, HRP linked antibody – 1:2000, #ab6728, Abcam.

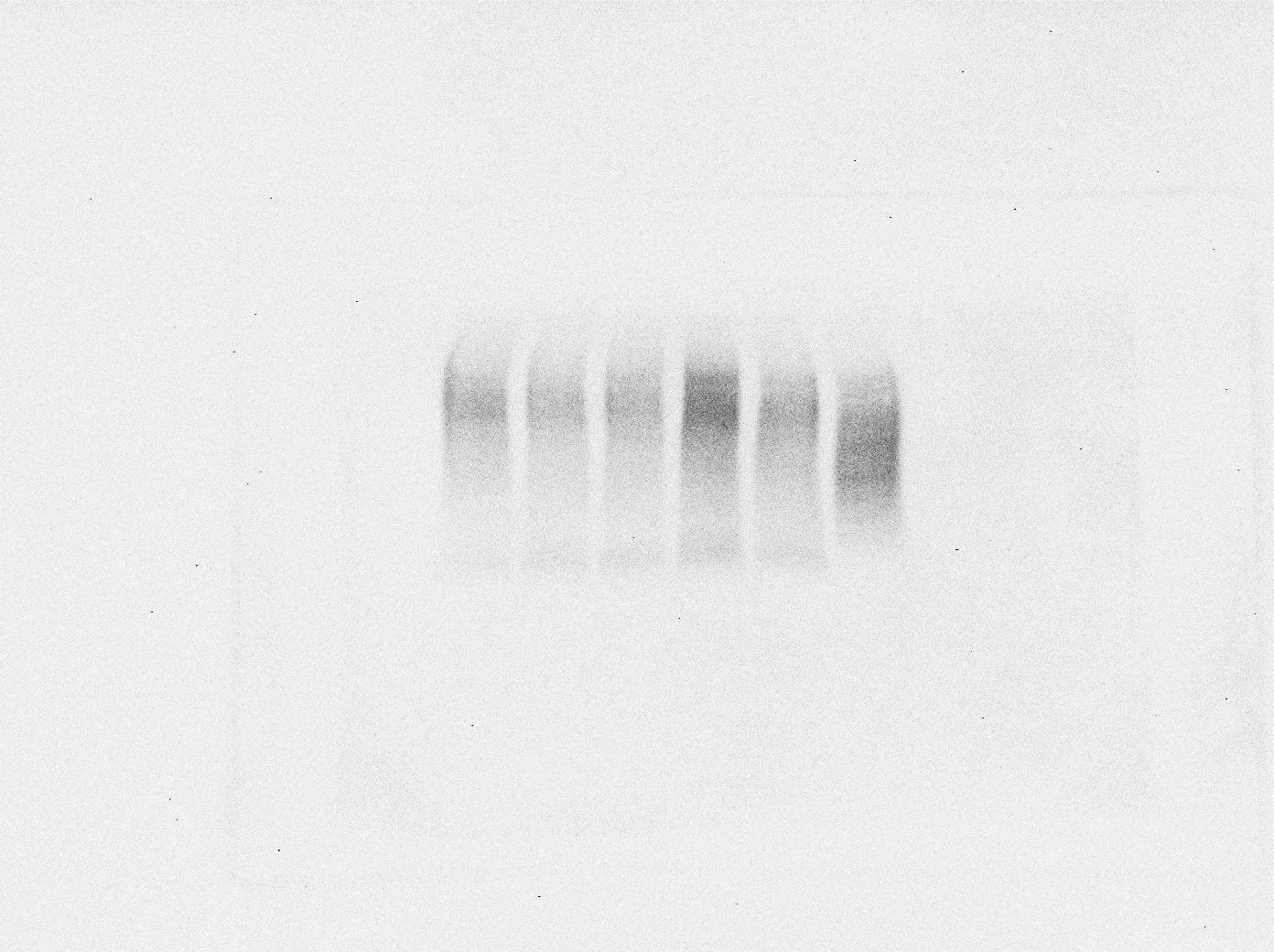

**WT**

**5xFAD**

**Caco2**

**HEK293**

**40-60 kDa – GLUT1**

**50-60 kDa - Tubulin**

******

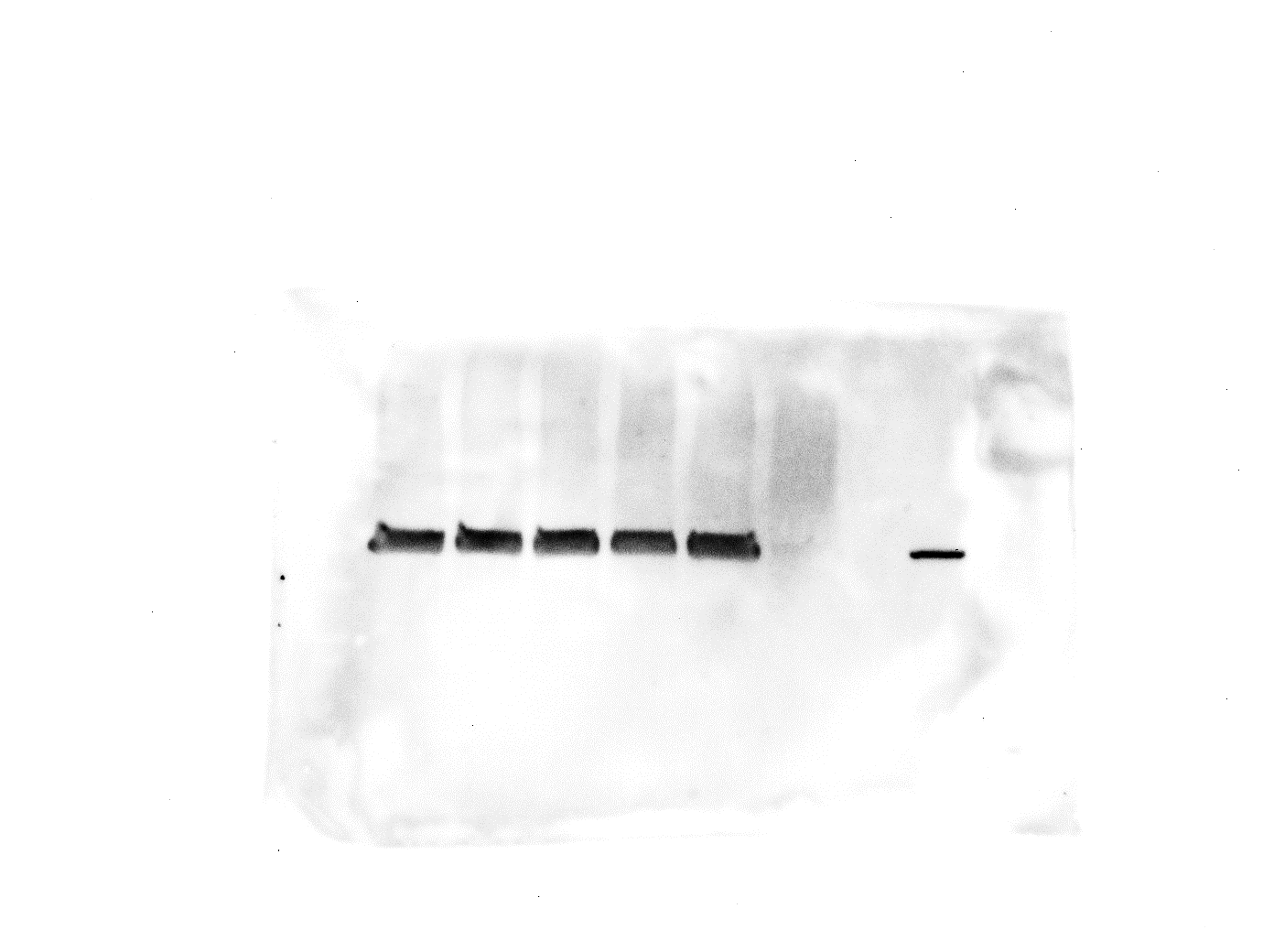

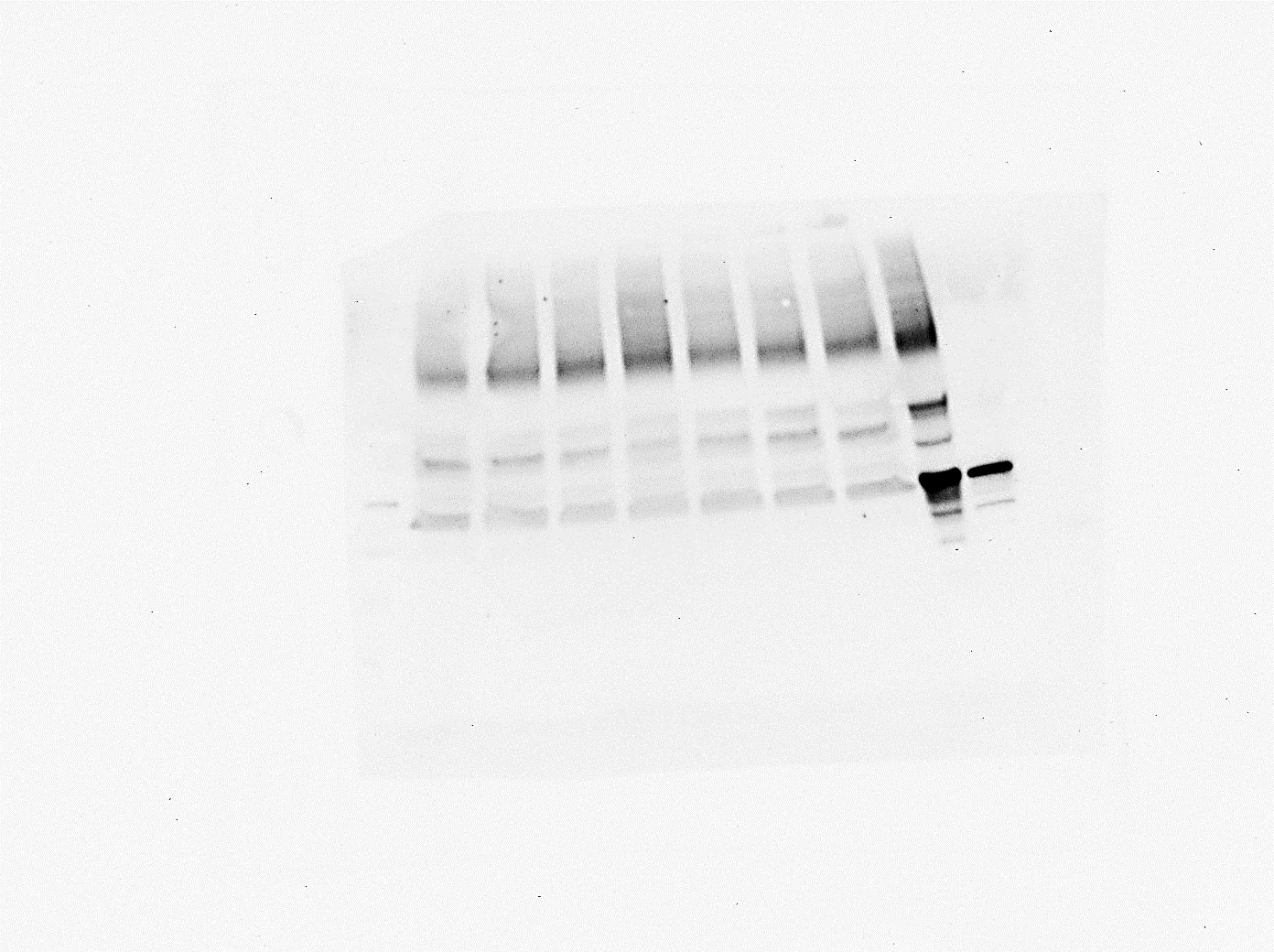

**37 kDa - GAPDH**

**140 kDa - P-gp**

**75 kDa - MATE1**

**HL-60**

**HEK293**

**5xFAD**

**WT**

Supplementary Figure 7: Brain capillary lysates (40 µg per well were loaded) from WT and 5xFAD mice were tested for P-gp (141 kDa), and MATE1 (75 kDa) expression, (n=6 WT and n=7 5xFAD mice per group, three membranes were ran, one membrane is presented). GAPDH (37 kDa) was used as a loading control. Antibodies used: anti-P-gp antibody – 1:1000, #ab170904, Abcam; anti-MATE1 – 1:1000, #ANT-131, Alomone Labs; anti-GAPDH antibody – 1:2500, #ab9485, Abcam; secondary anti-rabbit IgG, HRP-linked antibody – 1:1000, #7074, Cell Signalling Technology. Kidney lysates from a C57BL6 mouse and HEK293 whole cell lysate (#ab7902, Abcam, UK) were used as positive controls for P-gp and MATE1, HL-60 whole cell lysate (#ab7914, Abcam, UK) was used as a negative control.

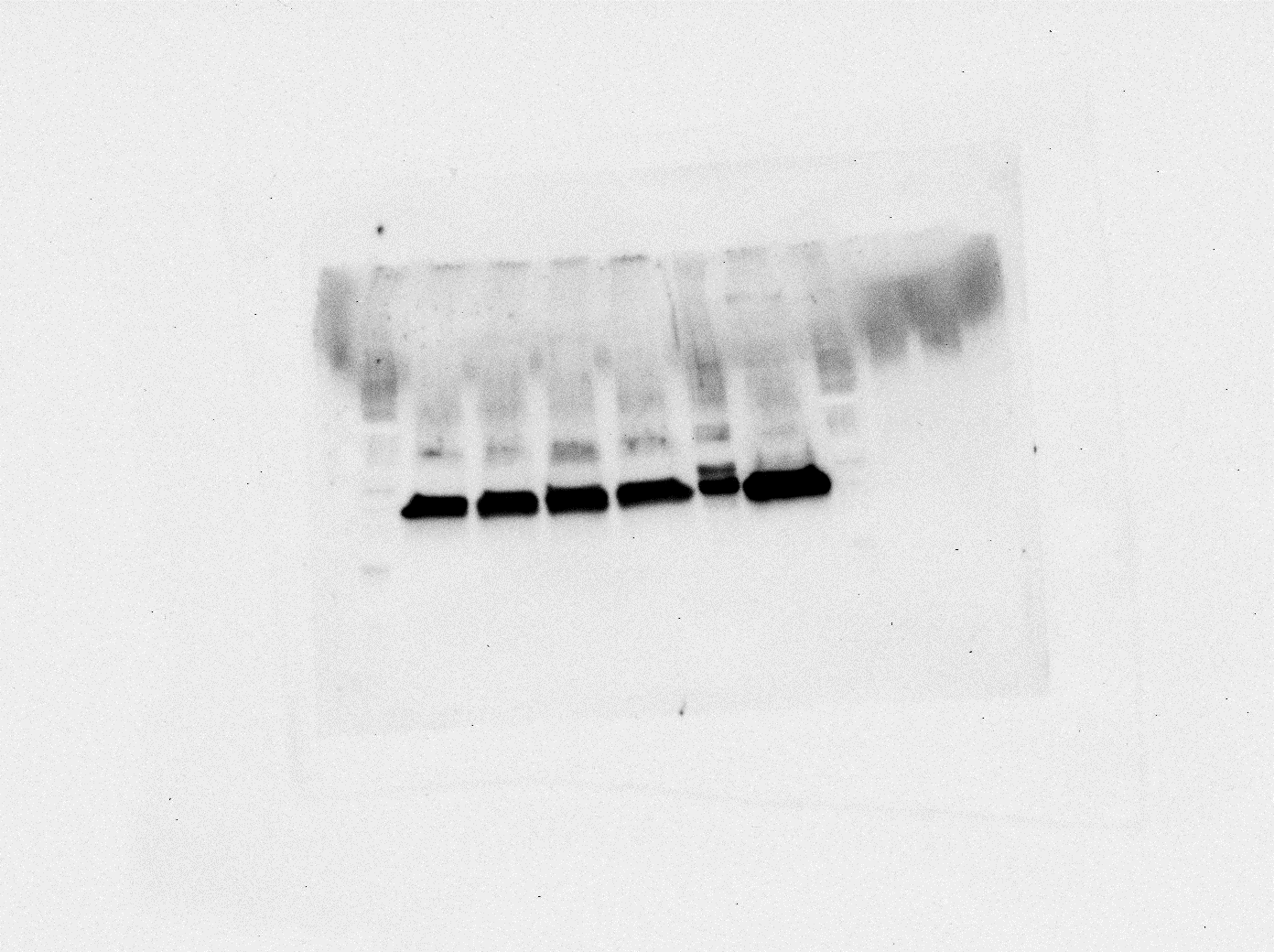

**37 kDa - GAPDH**

**61 kDa - OCT1**

**HL-60**

**Calu3**

**5xFAD**

**WT**

**Supplementary Figure 8:** Mouse brain capillary lysates (40 µg per well were loaded) from WT and 5xFAD mice were tested for OCT1 (61 kDa) expression, (n=5 WT and n=6 5xFAD mice per group, three membranes, one membrane is shown). GAPDH (37 kDa) was used as a loading control. Antibodies used: anti-OCT1 antibody – 1:667, #abab55916, Abcam; anti-GAPDH antibody – 1:2500, #ab9485, Abcam; secondary anti-rabbit IgG, HRP-linked antibody – 1:1000, #7074, Cell Signalling Technology. HL-60 cell line lysates (#ab7914, Abcam, UK) and Calu3 cell lysates were respectively used as a positive and negative control for OCT1.

**Intensity ratio of TfR1, and GLUT1 as well as the SLC transporters PMAT, MATE1, OCT1, and the ABC transporter P-gp in WT and 5xFAD**

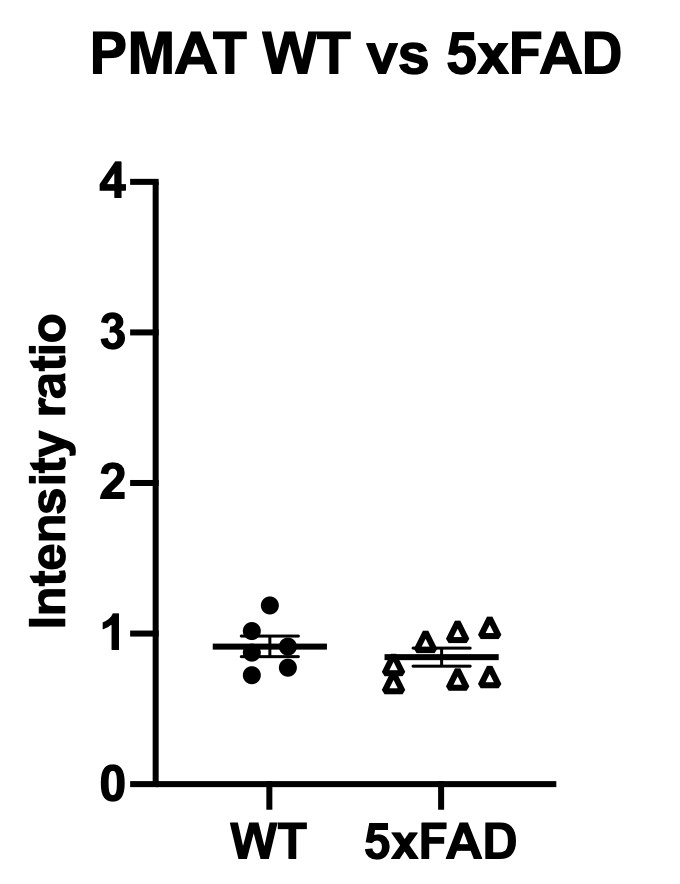

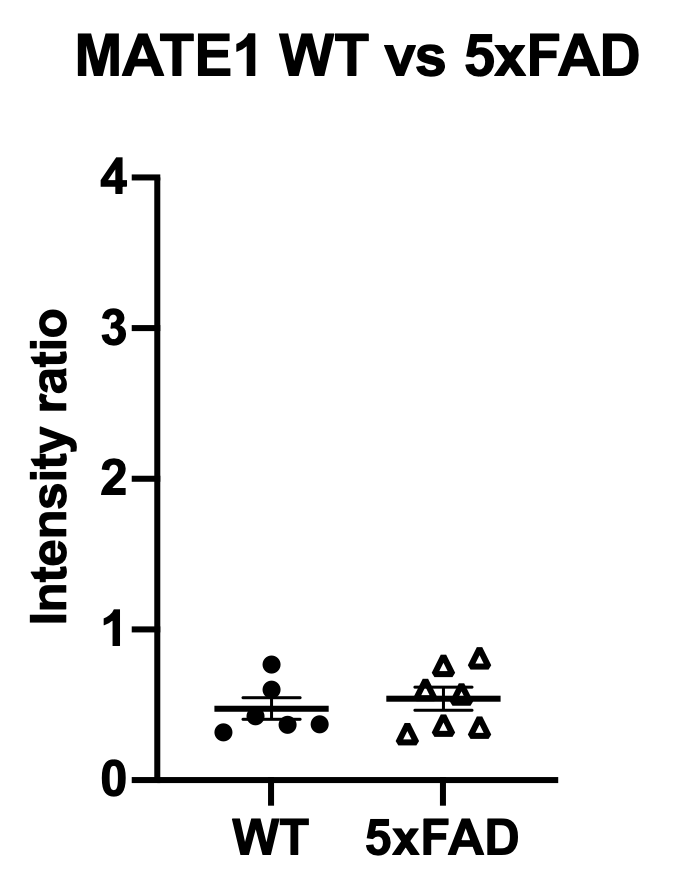

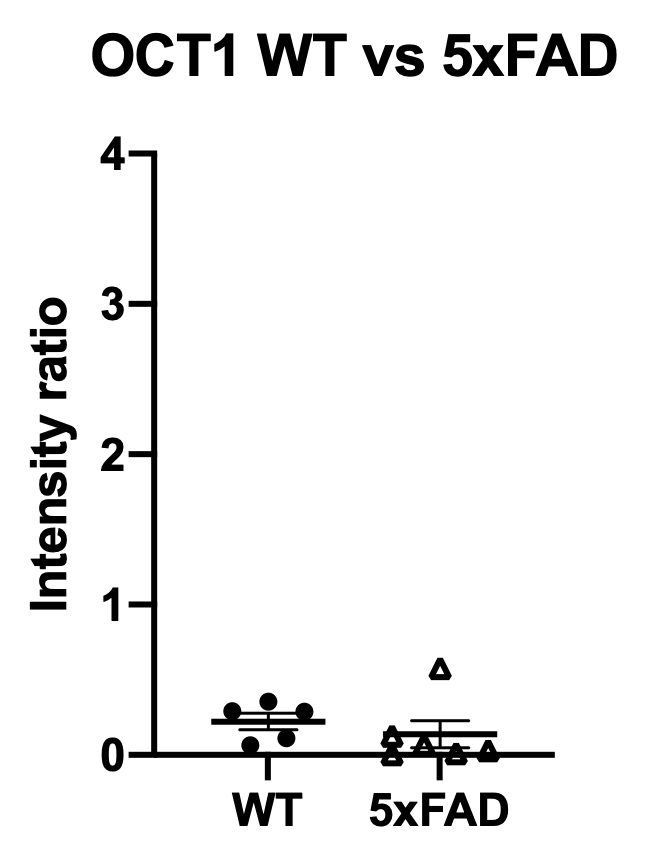

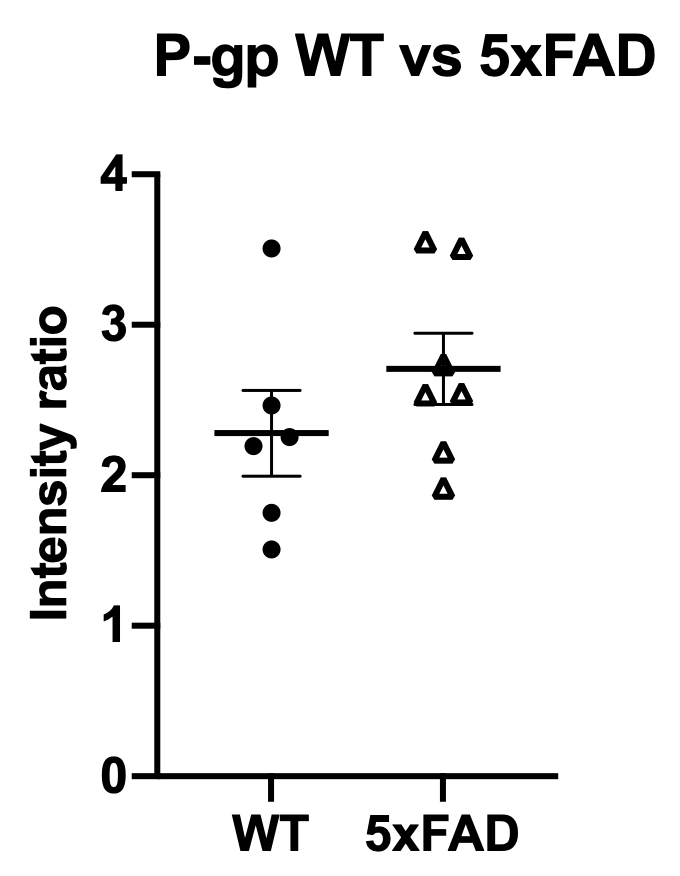

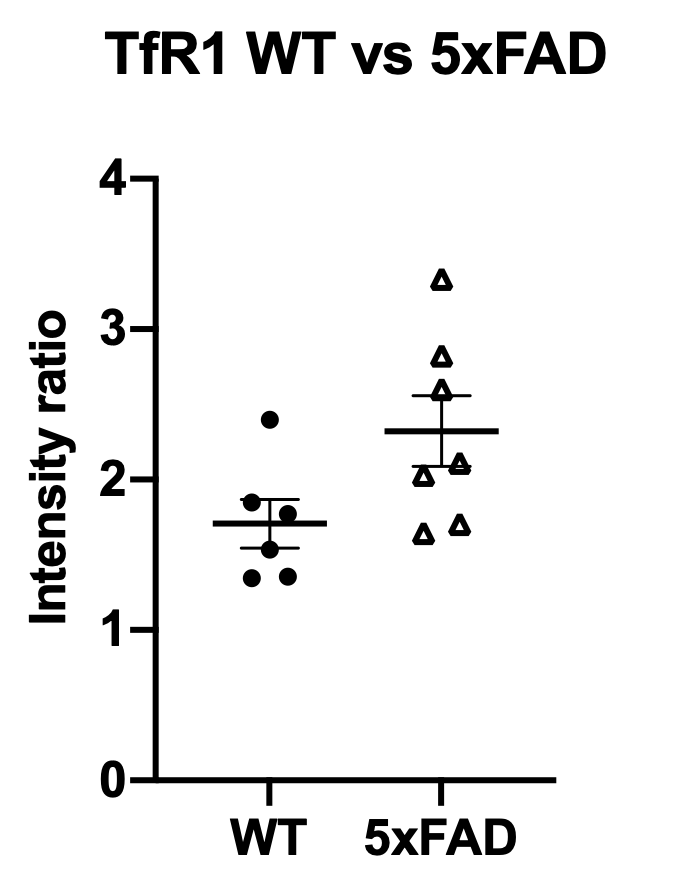

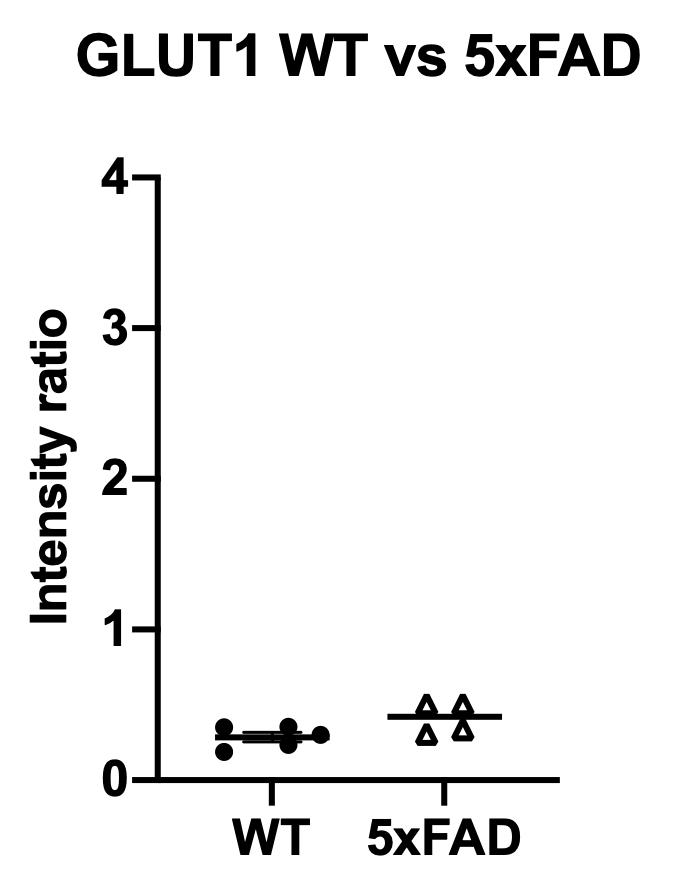

Supplementary Figure 9: Intensity ratio of TfR, and GLUT1 as well as the SLC transporters MCT1, PMAT, MATE1, OCT1, and the ABC transporter P-gp in WT and 5xFAD. Each dot represents data from one mouse, data is presented as mean±SEM. The WB images were analysed with Image J and graph were made with GraphPad Prism 9.0.

**Total protein concentration in the frontal cortex and caudate capillary lysates**

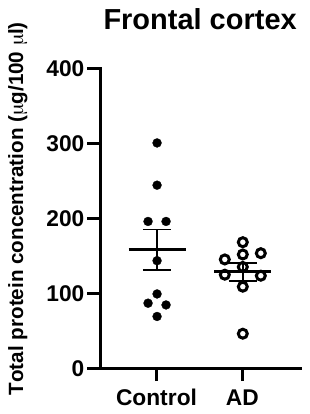

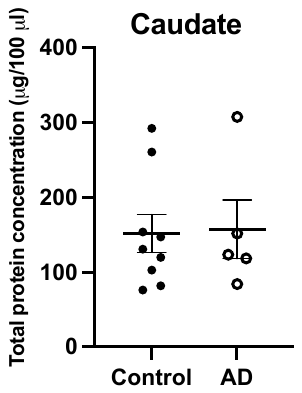

Supplementary Figure 10: Total protein concentration in the frontal cortex and caudate capillary lysates: control and AD cases. Expressed as µg protein per 100 µl lysate. Each dot represents total protein concentration per case in the frontal cortex and the caudate, respectively. Results are presented as mean±SEM, and are analysed with unpaired two-tailed Student’s t-test.

**Representative WBs in control and AD human cases**

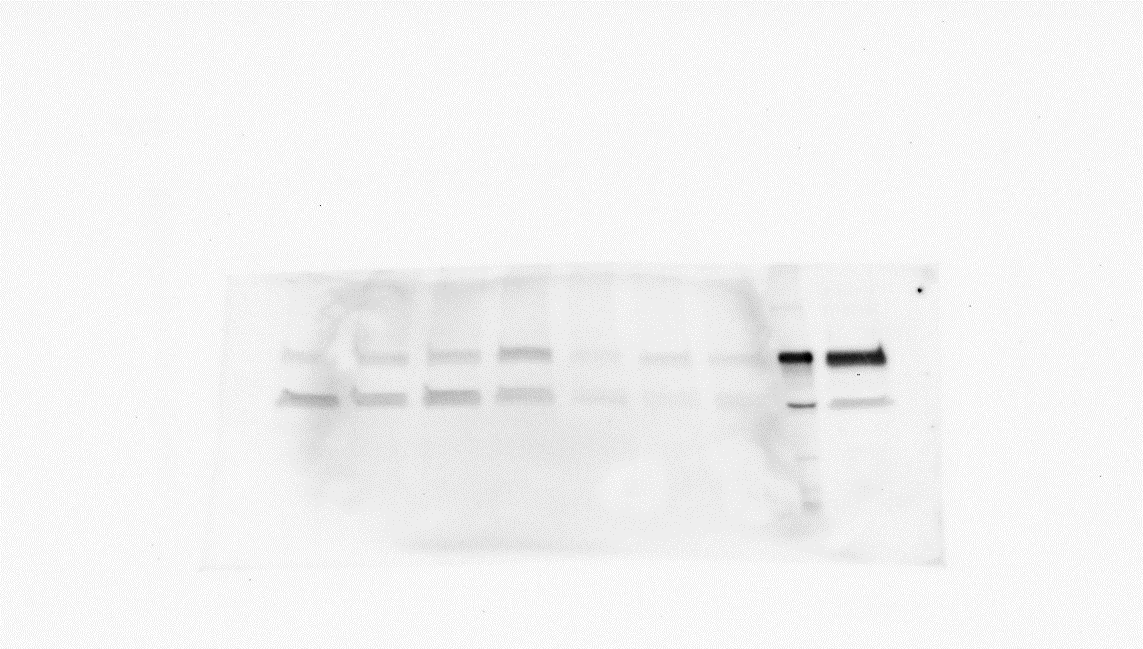

**≈90 kDa**

**Control Caudate**

**AD Caudate**

**HL-60**

**Calu3**

**37 kDa**

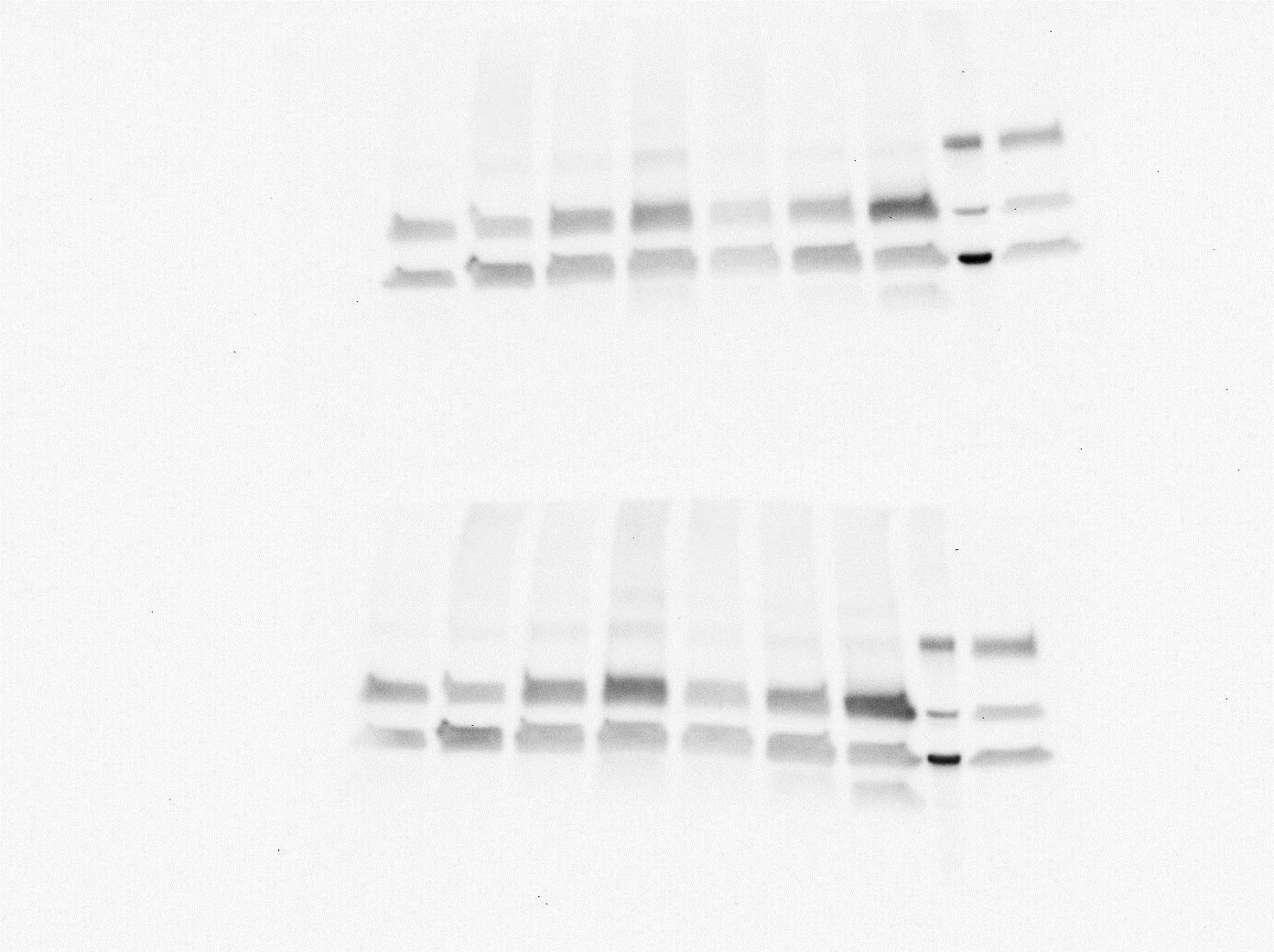

**Calu3**

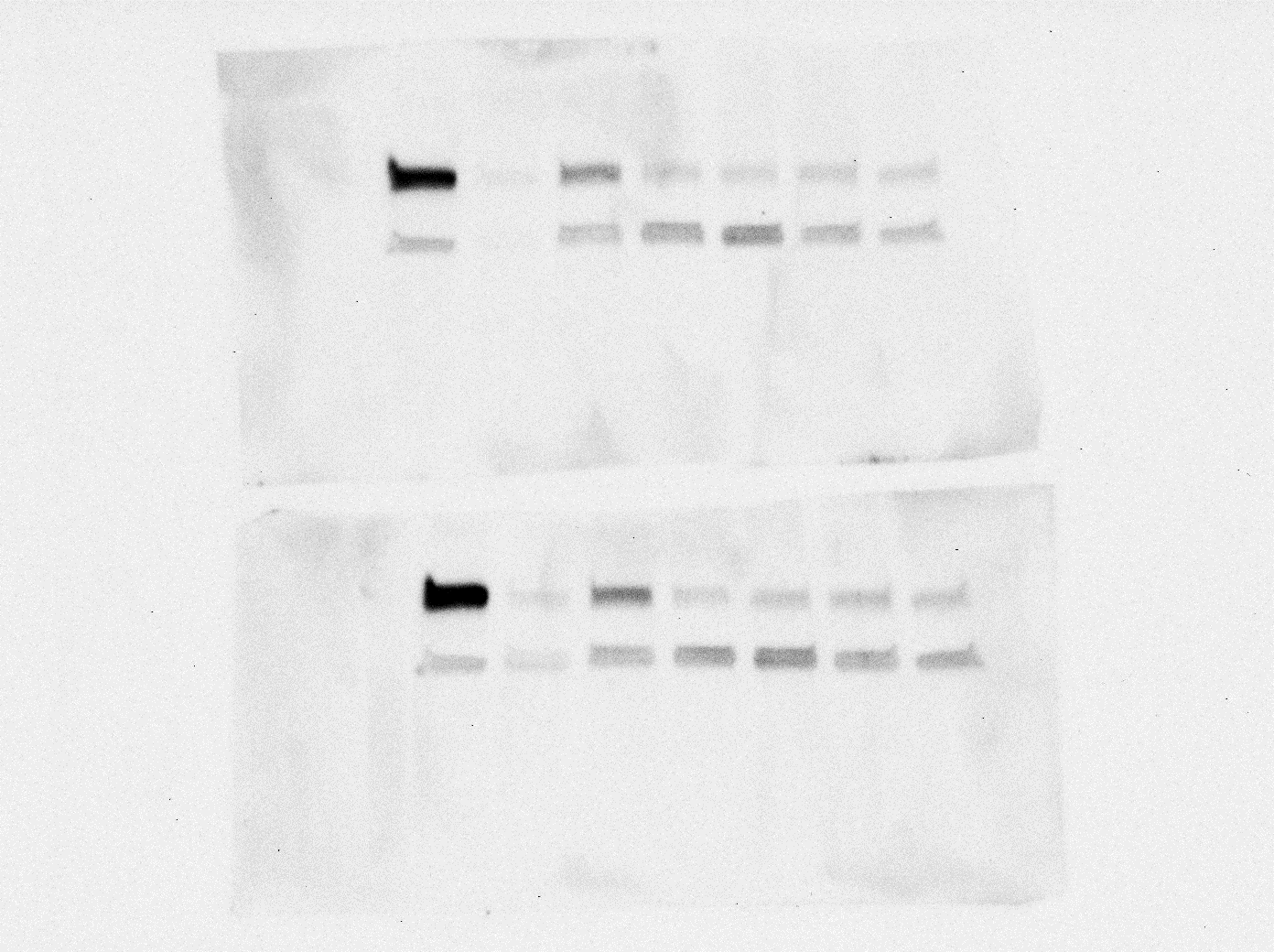

**GAPDH**

**TfR1**

**Control Frontal cortex**

**AD FC**

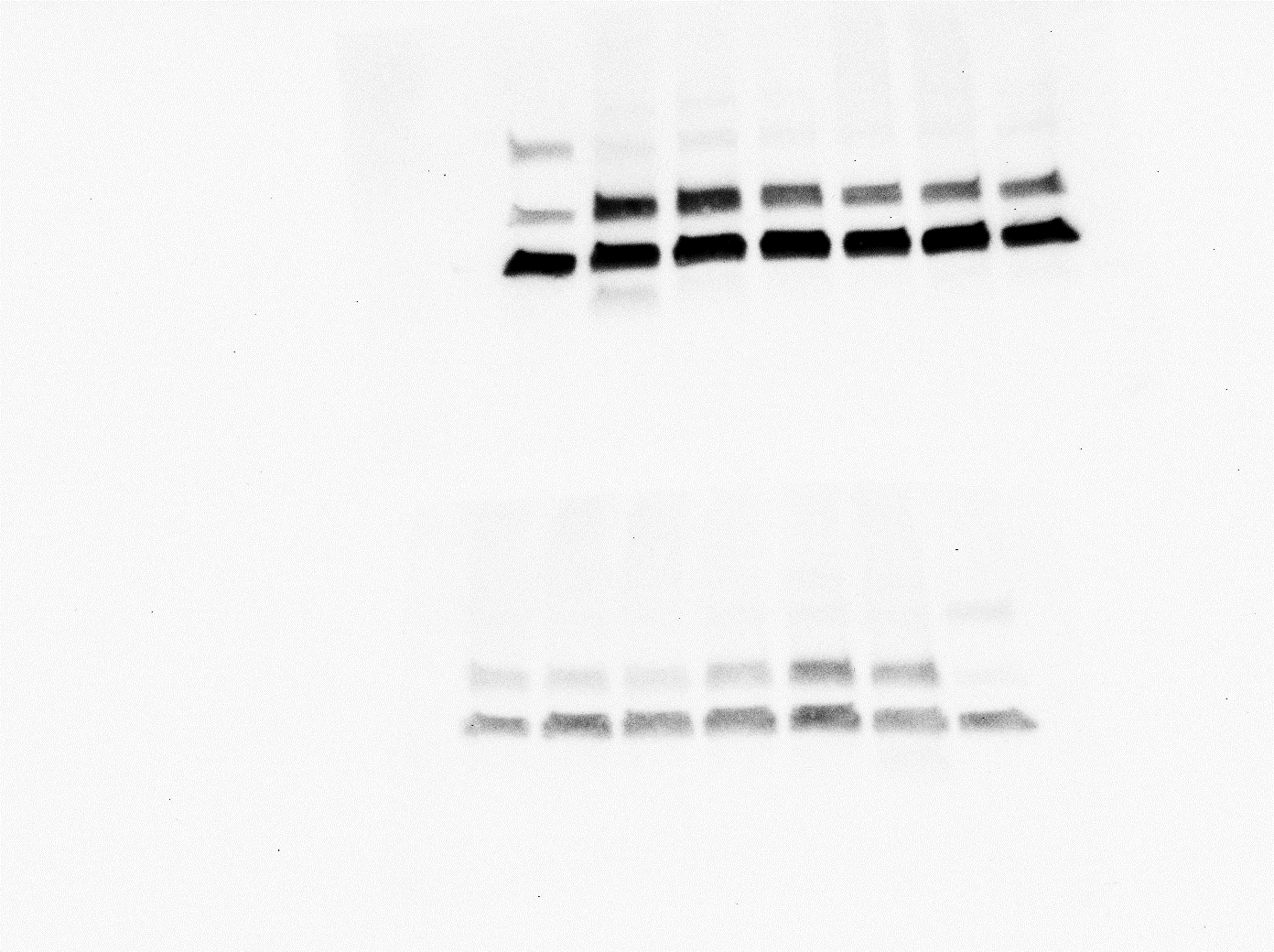

**Calu3**

Supplementary Figure 11: Frontal cortex capillary lysates (25 µg per well were loaded) from control and AD cases were tested for TfR1 (90 kDa) (n=9 Control, n=9 AD cases across 3 membranes, three technical repeats of each). Caudate capillary lysates (25 µg per well were loaded) from control and AD cases were tested for TfR1 (90 kDa), (n=7 Control, n=8 AD cases across three membranes, three technical repeats of each, except for one control sample). GAPDH (37 kDa) was used as a loading control. Antibodies used: anti-TfR1 antibody – 1:1000, #13-6800, Thermo Fisher, secondary anti-mouse IgG H&L HRP-linked antibody – 1:3000, #ab6728, Abcam; anti-GAPDH antibody – 1:2500, #ab9485, secondary anti-rabbit HRP-linked antibody – 1:2000, #7074, Cell Signalling Technology. HL-60 and Calu3 were used as controls for MCT1, ran on the same membrane.

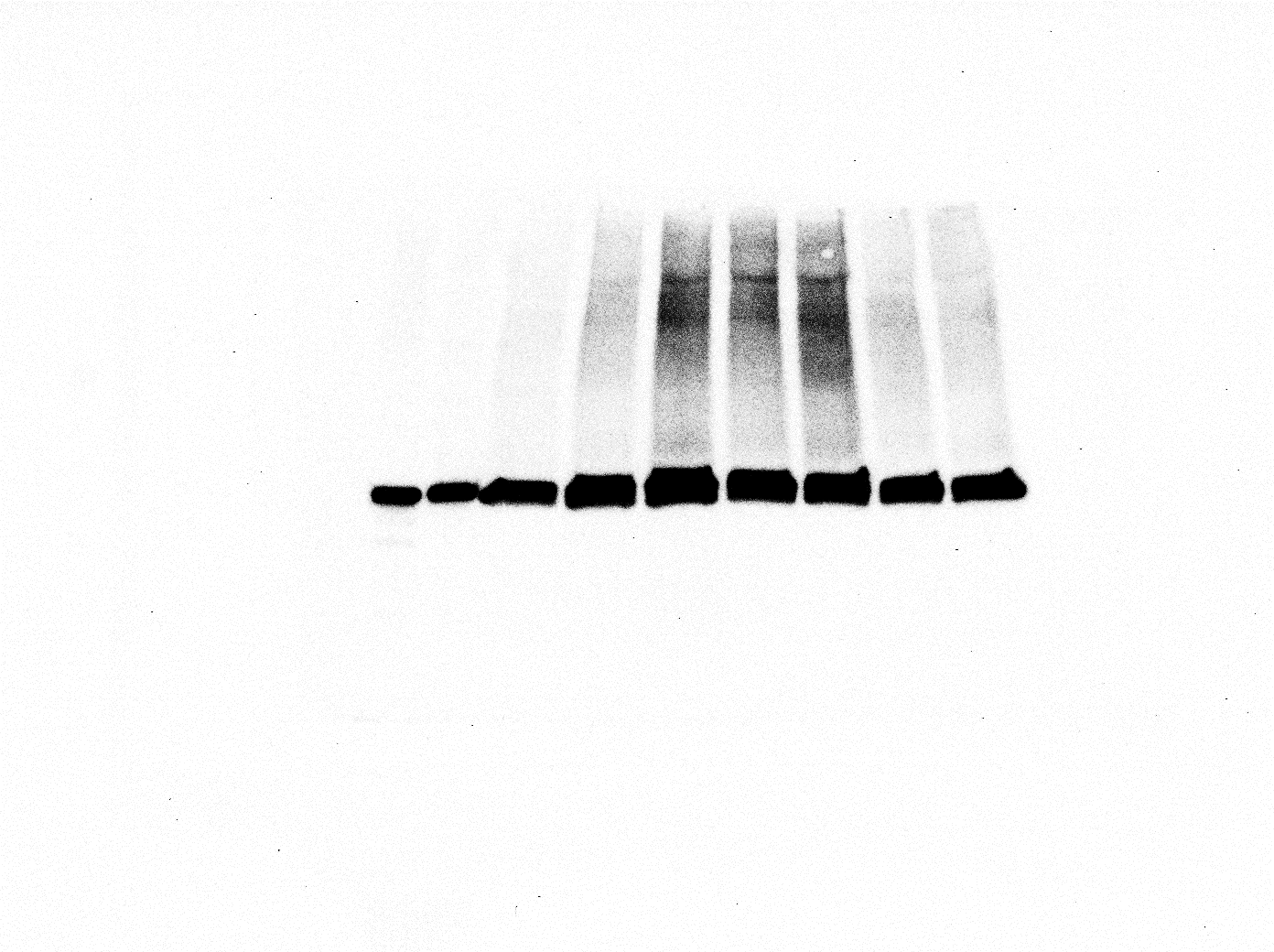

**Control FC**

**AD FC**

**Calu3**

**HEK**

**293**

**HL-60**

**GLUT1**

**GAPDH**

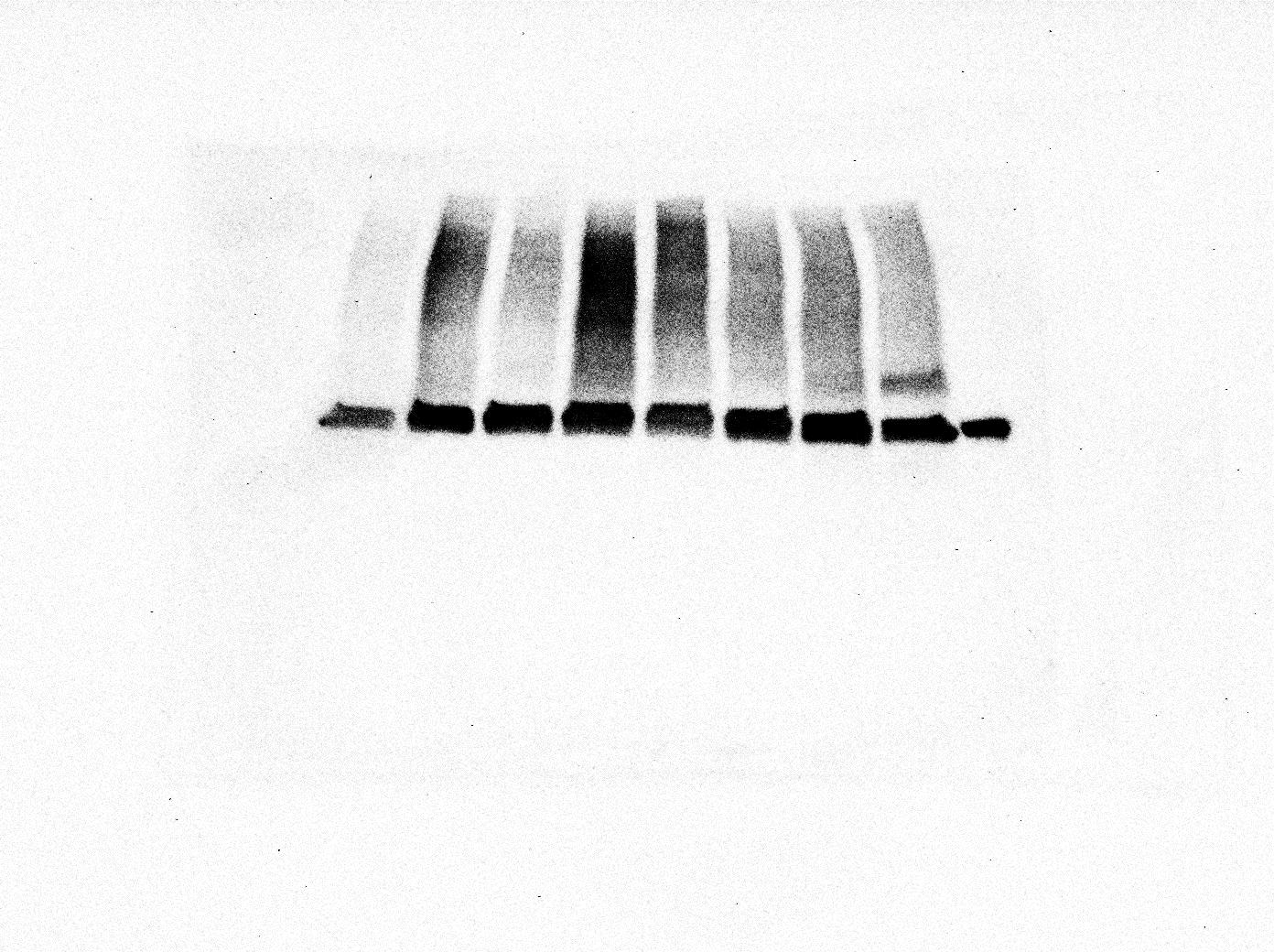

**Control Caudate**

**AD Caudate**

**HEK293**

**40-60 kDa**

**37 kDa**

Supplementary Figure 12: Frontal cortex capillary lysates (20 µg per well were loaded) from control and AD cases were tested for GLUT1 (40-60 kDa), (n=9 Control, n=13 AD cases across four membranes, three technical repeats of each, except for one AD sample). Caudate capillary lysates (20 µg per well were loaded) from control and AD cases were tested for GLUT1 (40-60 kDa), (n=9 Control, n=9 AD cases across three membranes, three technical repeats of each). One membrane is shown. GAPDH (37 kDa) was used as a loading control, Caco2 and Calu3 lysates were used as a positive control, HEK293 cell lysates were used as negative control. Antibodies used: anti-GLUT1 antibody – 1:100 000, #ab115730, Abcam; anti-GAPDH antibody – 1:2500, #ab9485, Secondary anti-rabbit HRP-linked antibody – 1:2000, #7074, Cell Signalling Technology.

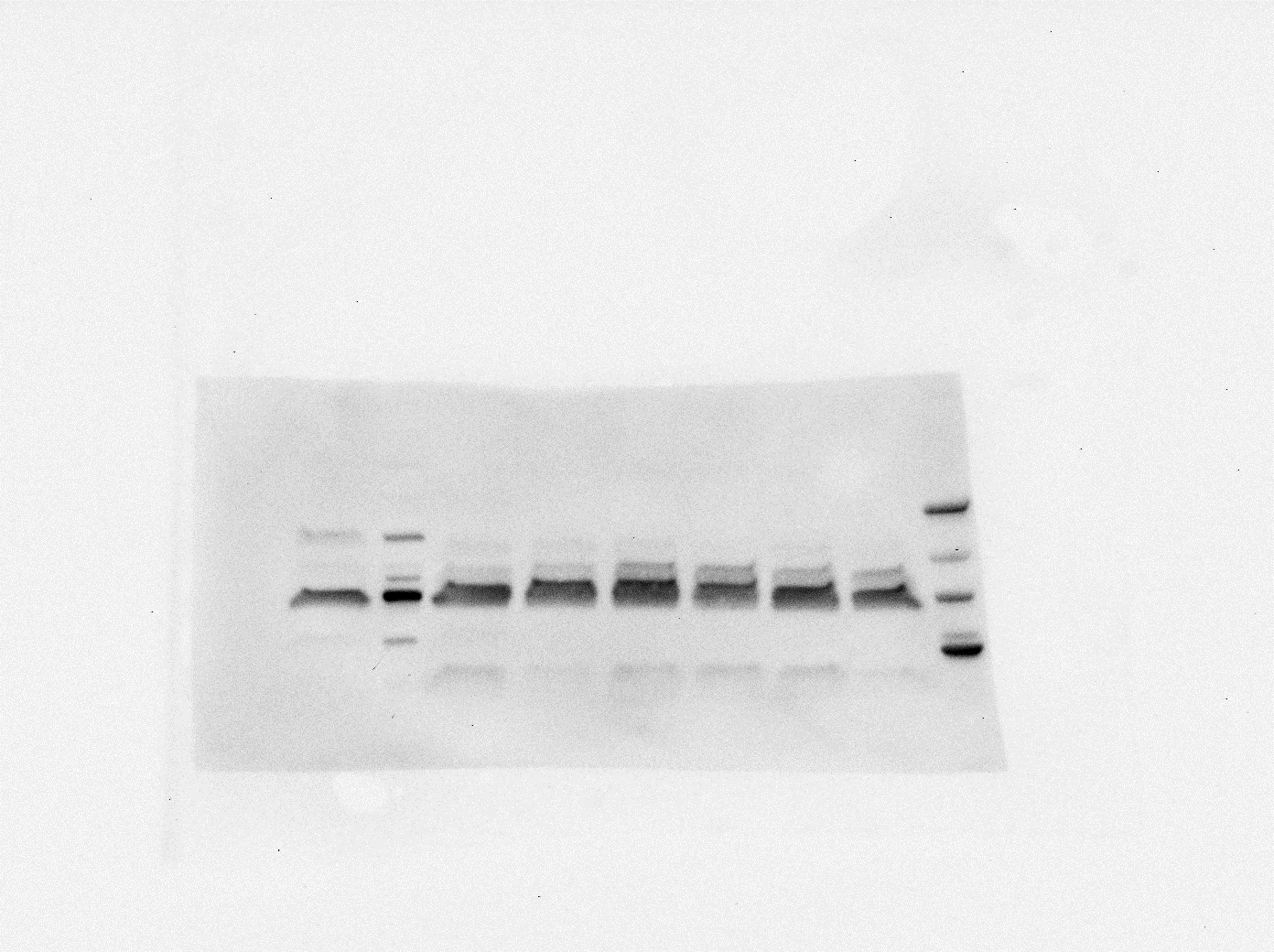

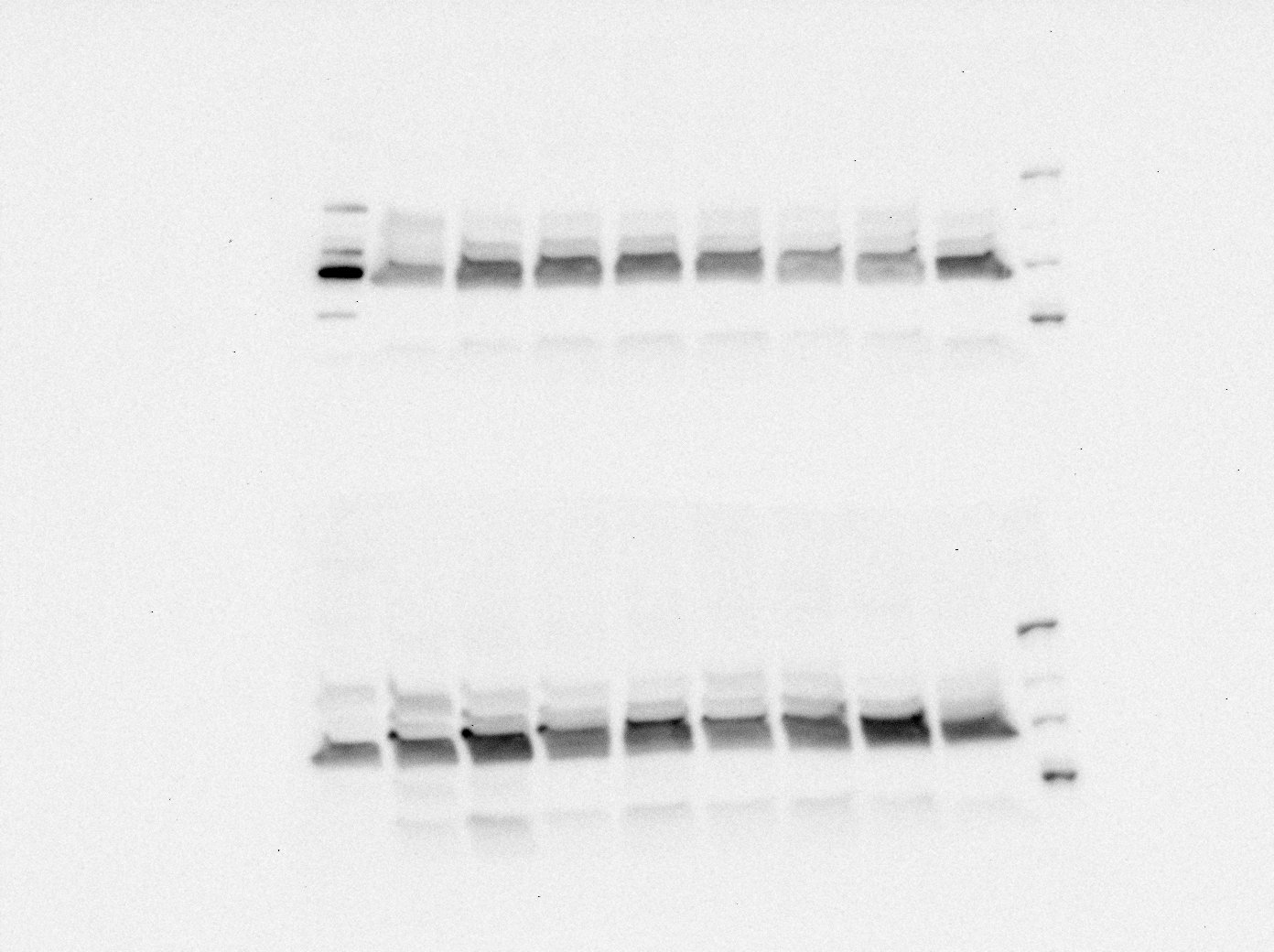

**37 kDa**

**GAPDH**

**58 kDa**

**PMAT**

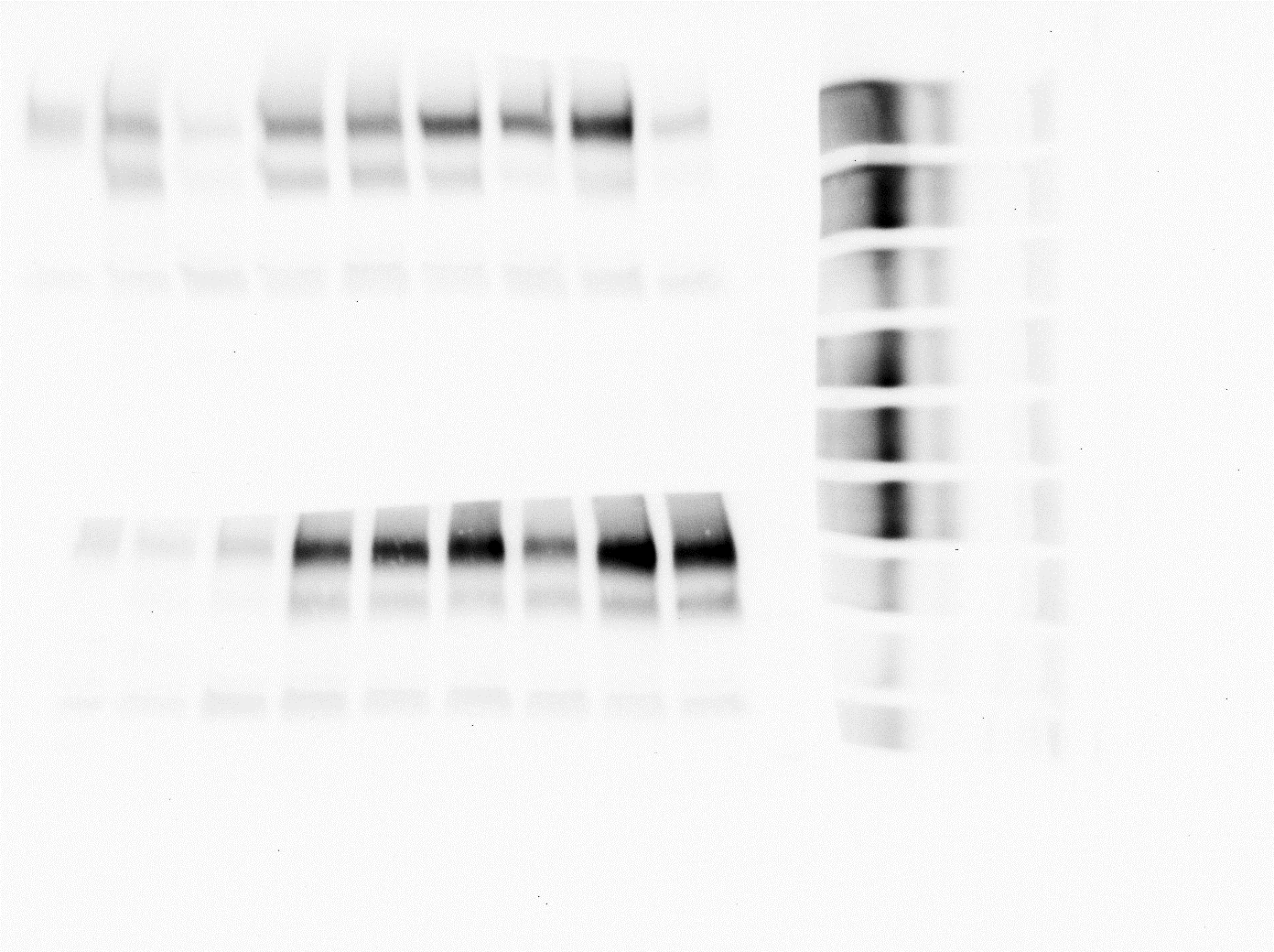

**141 kDa**

**37 kDa**

**Control Caudate**

**HEK293**

**AD Caudate**

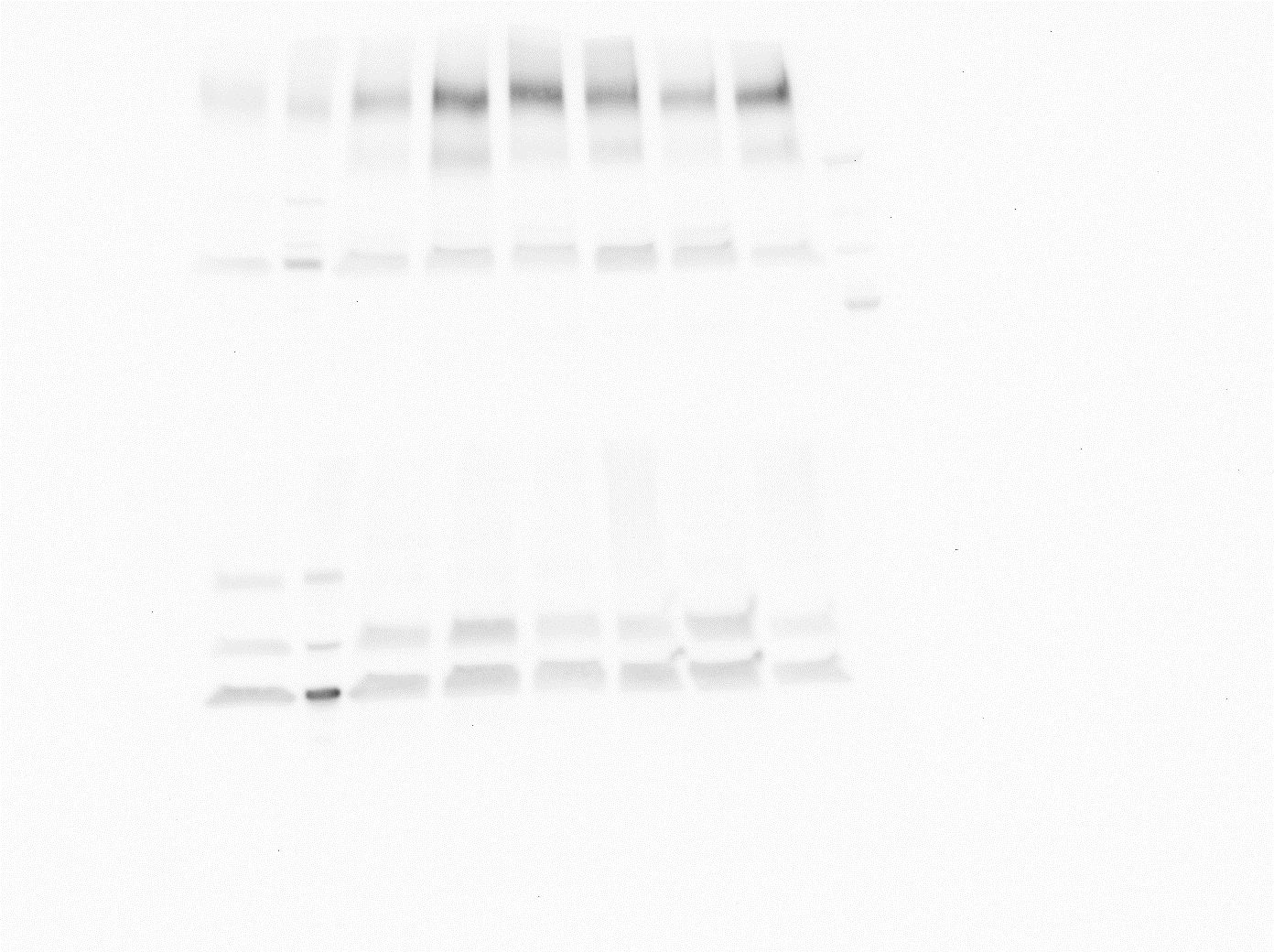

**AD FC**

**Control Frontal cortex**

**HEK293**

**Calu3**

**P-gp**

**GAPDH**

Supplementary Figure 13: Frontal cortex capillary lysates (20 µg per well were loaded) from Control and AD cases were tested for P-gp (141 kDa), (n=9 control, n=9 AD cases across three membranes, three technical repeats of each). Caudate capillary lysates (20 µg per well were loaded) from Control and AD cases were tested for P-gp (141 kDa), (n=9 Control, n=7 AD cases across three membranes, three technical repeats of each). GAPDH (37 kDa) was used as a loading control. HEK293 cell lysate was used as a positive control for P-gp. Antibodies used: anti-P-gp antibody – 1:1000, #ab170904, Abcam, Secondary anti-rabbit HRP-linked antibody – 1:2000, #7074, Cell Signalling Technology; anti-GAPDH antibody – 1:2500, #ab9485, Secondary anti-rabbit HRP-linked antibody – 1:2000, #7074, Cell Signalling Technology. The membrane was also stained for PMAT, PMAT is marked on the image but not discussed in this paper.

**References**

1.	Abcam Primary Antibodies. https://www.abcam.com/nav/primary-antibodies Accessed 21 June 2022.

2.	Cell Signalling Technology. https://www.cellsignal.co.uk/ Accessed 21 June 2022.

3.	Alomone Labs. https://www.alomone.com Accessed 21 June 2022.

4.	Bioss USA. https://www.biossusa.com/ Accessed 21 June 2022.

5.	Thermo Fisher Scientific. https://www.thermofisher.com/uk/en/home.html Accessed 21 June 2022.

6.	UK brain banks network: database of tissue samples. https://brainbanknetwork.ac.uk Accessed 17 November 2019.

7.	Wishart DS, Feunang YD, Guo AC, Lo EJ, Marcu A, Grant JR, et al. DrugBank 5.0: a major update to the DrugBank database for 2018. Nucleic Acids Res. 2018;46(D1):D1074–82.

8.	MarvinSketch version 22.9.0. ChemAxon http://chemaxon.com Accessed April 2022 and April 2023.

9.	Augustin R. The protein family of glucose transport facilitators: It’s not only about glucose after all. IUBMB Life. 2010 May;62(5):315–33.

10.	Mueckler M, Thorens B. The SLC2 (GLUT) family of membrane transporters. Mol Aspects Med. 34(2–3):121–38.

11.	Chiba Y, Murakami R, Matsumoto K, Wakamatsu K, Nonaka W, Uemura N, et al. Glucose, Fructose, and Urate Transporters in the Choroid Plexus Epithelium. Int J Mol Sci. 2020 Sep 30;21(19).

12.	Montel-Hagen A, Kinet S, Manel N, Mongellaz C, Prohaska R, Battini JL, et al. Erythrocyte Glut1 triggers dehydroascorbic acid uptake in mammals unable to synthesize vitamin C. Cell. 2008 Mar 21;132(6):1039–48.

13.	Barbosa AM, Martel F. Targeting Glucose Transporters for Breast Cancer Therapy: The Effect of Natural and Synthetic Compounds. Cancers (Basel). 2020 Jan 8;12(1).

14.	Holman GD. Structure, function and regulation of mammalian glucose transporters of the SLC2 family. Pflugers Arch. 2020;472(9):1155–75.

15.	Zambrano A, Molt M, Uribe E, Salas M. Glut 1 in Cancer Cells and the Inhibitory Action of Resveratrol as A Potential Therapeutic Strategy. Int J Mol Sci. 2019 Jul 9;20(13).

16.	Samec M, Liskova A, Koklesova L, Samuel SM, Zhai K, Buhrmann C, et al. Flavonoids against the Warburg phenotype-concepts of predictive, preventive and personalised medicine to cut the Gordian knot of cancer cell metabolism. EPMA J. 2020 Sep;11(3):377–98.

17.	Zezina E, Sercan-Alp O, Herrmann M, Biesemann N. Glucose transporter 1 in rheumatoid arthritis and autoimmunity. Wiley Interdiscip Rev Syst Biol Med. 2020;12(4):e1483.

18.	Dwyer DS, Pinkofsky HB, Liu Y, Bradley RJ. Antipsychotic drugs affect glucose uptake and the expression of glucose transporters in PC12 cells. Prog Neuropsychopharmacol Biol Psychiatry. 1999 Jan;23(1):69–80.

19.	Dwyer DS, Donohoe D. Induction of hyperglycemia in mice with atypical antipsychotic drugs that inhibit glucose uptake. Pharmacol Biochem Behav. 2003 May;75(2):255–60.

20.	Dwyer DS, Lu XH, Bradley RJ. Cytotoxicity of conventional and atypical antipsychotic drugs in relation to glucose metabolism. Brain Res. 2003 May;971(1):31–9.

21.	Dwyer DS, Liu Y, Bradley RJ. Dopamine receptor antagonists modulate glucose uptake in rat pheochromocytoma (PC12) cells. Neurosci Lett. 1999 Oct;274(3):151–4.

22.	Ardizzone TD, Bradley RJ, Freeman AM, Dwyer DS. Inhibition of glucose transport in PC12 cells by the atypical antipsychotic drugs risperidone and clozapine, and structural analogs of clozapine. Brain Res. 2001 Dec 27;923(1–2):82–90.

23.	de Silva PN. Does the association with diabetes say more about schizophrenia and its treatment?--the GLUT hypothesis. Med Hypotheses. 2011 Oct;77(4):529–31.

24.	Stapel B, Kotsiari A, Scherr M, Hilfiker-Kleiner D, Bleich S, Frieling H, et al. Olanzapine and aripiprazole differentially affect glucose uptake and energy metabolism in human mononuclear blood cells. J Psychiatr Res. 2017;88:18–27.

25.	Babkin P, George Thompson AM, Iancu C v, Walters DE, Choe JY. Antipsychotics inhibit glucose transport: Determination of olanzapine binding site in Staphylococcus epidermidis glucose/H(+) symporter. FEBS Open Bio. 2015;5:335–40.
